# Exploration of mRNA-sized RNA import into *Saccharomyces cerevisiae* mitochondria by a combined synthetic biology and adaptive laboratory evolution approach

**DOI:** 10.1101/2025.08.28.672929

**Authors:** Charlotte C. Koster, Kavish Kohabir, Maxime den Ridder, Marijke A.H. Luttik, Erik de Hulster, Martin Pabst, Pascale Daran-Lapujade

## Abstract

Efficient gene integration using RNA-guided endonucleases has not yet been achieved in the mitochondrial genome. Import of nucleic acids into mitochondria, a controversial feature, is essential for implementation of Cas9-mediated genome engineering of mitochondria. Import of short RNAs naturally occurs in mitochondria, and several putative import mechanisms and determinants have been proposed. However to date, import of gene-length RNA, required for gene integration in the mitochondrial genome, has never been described.

The goal of this study was to devise and test experimental strategies to detect and improve the import of mRNA-sized RNA in mitochondria, using *S. cerevisiae* as model. A first fluorescence-based screening approach, relying on mitochondrial import of a fluorescent protein encoding mRNA was analyzed by fluorescence measurements, western blot and mRNA-FISH. Confounding results obtained with these different techniques made it difficult to unambiguously conclude on the occurrence of import of mRNA-sized RNAs into mitochondria. An adaptive laboratory evolution (ALE) approach, imposing a strong selection pressure for mRNA import to mitochondria, was then designed and tested to improve mitochondrial mRNA import. While the ALE approach did not improve mitochondrial mRNA import in the present study, it is a promising, unambiguous method for future studies testing different RNAs or mutants. The present study highlights remaining challenges in analytical techniques to identify RNA import to mitochondria, and introduces a novel application of ALE for studies on mitochondrial import of short and long RNA species.

## Introduction

The development of the CRISPR/Cas9 technology has revolutionized genome editing [1]. However, even the most genetically accessible eukaryotes harbor a genome that is not yet amenable for RNA-guided endonuclease (RGEN) editing: the mitochondrial genome. Mitochondria perform many (essential) cellular functions including the conservation of metabolic energy in the form of ATP, the maintenance of the cellular redox state, as well as production of co-factors, amino acids and lipids [2]. It is generally accepted that mitochondria have an endosymbiotic origin [3]. As a result, mitochondria have their own transcription and translation machineries, as well as a reduced genome encoding ribosomal subunits, a full set of tRNAs and several subunits of the respiratory chain. 1 % of the mitochondrial proteome is encoded on this mitochondrial genome, the rest is imported from the cytosol [4–6]. Apart from their key role in many eukaryotic cellular processes, mitochondria are also the source of many human diseases: over 250 pathogenic mutations in the mitochondrial DNA (mtDNA) have been identified [7]. Therefore, there have been substantial efforts in engineering mitochondrial genomes mostly focused on therapeutic applications [8, 9]. These efforts have led to the development of protein-based DNA editing strategies including base editors and zinc-finger- or TALE nucleases [10–13], which are limited to base exchange or depletion of mutated heteroplasmic mtDNA variants. More extensive modifications of mtDNA, such as gene integration or targeted deletion, would enable the reprogramming of mitochondrial genomes. This could either be applied to treat diseases, to endow mitochondria with new functions for metabolic engineering purposes, or even use mitochondria as chassis for the construction of synthetic cells. However, the large spectrum of applications of engineering of mitochondrial genomes is impeded by their poor genetic accessibility.

Currently, only microprojectile bombardment of DNA-coated metal particles (biolistic transformation) enables engineering of the mtDNA beyond base editing. This invasive method of mtDNA editing is restricted to two organisms, the yeast *Saccharomyces cerevisiae,* and the microalga *Chlamydomonas reinhardtii* [14–16]. Expanding the mitochondrial genome editing ‘toolbox’ to include the flexible and less invasive CRISPR/RGEN editing has proven very challenging [17, 18]. This is largely explained by the three-component nature of CRISPR/RGEN systems. Unlike the existing mitochondrial programmable endonucleases, RGEN function depends on nucleic acids [19]. A guide RNA (gRNA) is required to direct the RGEN towards the editing site, and the induced double-stranded break can be repaired by a DNA molecule (repair DNA). While targeting proteins to the mitochondrial matrix is achieved by addition of mitochondrial targeting sequences (MTS [20]), targeting DNA and RNA molecules presents a substantial challenge. Mitochondria are not naturally competent for DNA, but native uptake of RNA in mitochondria has been reported for tRNAs in yeasts, plants and mammals (reviewed by Schneider (21)). Also, the ribonucleic subunits of RNases P and MRP, and rRNA were reported to be imported in mammalian mitochondria [22–24]. The mechanistic understanding of mitochondrial RNA import is limited to the import mechanism of yeast tRNA^Lys^(CUU). The tRNA is first recruited to the mitochondrial membrane by binding Eno2p and subsequently bound by the precursor of mitochondrial lysyl-tRNA synthetase (pre-LysRS). Pre-LysRS is imported in mitochondria and likely co-imports the associated tRNA^Lys^(CUU) [25]. This led to the hypothesis that certain RNA-sequences or structures of the RNA promote RNA import. Hence, multiple RNA ‘import sequences’ were derived from tRNA^Lys^(CUU), 5S rRNA, mammalian RNAse P and RNase MRP that reportedly improve RNA import *in vitro* and occasionally *in vivo* [26–30].

Several studies have described successful gRNA targeting to the mitochondria through addition of RNA import sequences, reportedly leading to successful Cas9 editing of mtDNA [31–36]. In most instances, editing was measured by monitoring mtDNA depletion, in lieu of a DNA repair. However, no definitive proof of RGEN-induced DNA-editing in the mitochondria, such as a site-specific shift in heteroplasmy, has been demonstrated to date. Since discrete and reproducible evidence of mitochondrial RNA localization and site-specific Cas9 activity on the mtDNA is lacking, several studies question the occurrence of gRNA import and mtDNA editing by Cas9 altogether [9, 10, 37–39]. These contrasting reports highlight the difficulty of the experimental design required to demonstrate if RNA is indeed imported into mitochondria and if site-specific RGEN-mediated mtDNA cleavage occurs. Even if gRNA import is successfully achieved, it is not sufficient for gene replacement or the addition of new functions in the mtDNA. This requires the integration of gene-length exogenous DNA, which is not natively imported in the mitochondria. However, repair DNA could be generated from mitochondrially-targeted RNA, through a mitochondrially-localized reverse transcriptase [40–42]. The use of such an RNA-mediated repair system would require the targeting of gene-length RNA to the mitochondria. There is little knowledge on the ability of mitochondria to import mRNA-sized RNA, as most studies focus on the import of short tRNA and gRNA (100 -200 nucleotides), while the import of a 1000-nucleotide long mRNA was only reported once [28].

The goal of the present study is to devise and test experimental strategies to detect and enable the import of mRNA-sized RNA in mitochondria, using *S. cerevisiae* as paradigm. *S. cerevisiae* is often used as model organism in mitochondrial biology as it is one of the few model eukaryotes that can survive (extensive) mtDNA damage [43]. Additionally, *S. cerevisiae* is a robust, genetically tractable, and well-characterized model eukaryote, making it a perfect host to explore RNA import to mitochondria and mtDNA editing. RNA import in mitochondria was first explored by screening for mitochondrial localization of an mRNA fused to a library of RNA import signals and its fluorescent translation product. Next to this systematic approach, an adaptive laboratory evolution strategy was devised in an attempt to select for mutants with improved mitochondrial RNA import.

## Materials and methods

### Strains, medium and cultivation

*Saccharomyces cerevisiae* strains used in this study were derived from a CEN.PK genetic background [44]. Unless indicated otherwise, yeast strains (Table 1) were grown aerobically at 30 °C in 500 mL shake flasks containing 100 mL minimal synthetic medium with ammonium as nitrogen source (SM) supplied with vitamins and trace elements, prepared and sterilized as described previously [37], in an Innova incubator shaker (Eppendorf AG, Hamburg, Germany) set at 200 rpm. Media were autoclaved at 121 °C for 20 minutes. Media for respiro-fermentative growth in shake-flasks contained 2% (w/v) D-glucose, for respiring cultures 2% (v/v) ethanol (SME) or 2% (v/v) ethanol and 2% (v/v) glycerol (SMEG) served as carbon sources Glucose solutions (50 % w/v) and glycerol solutions (99 % w/v) were autoclaved separately for 10 minutes at 110 °C. Where relevant, media were complemented with Hygromycin B Gold (Invivogen, San Diego, CA, USA) to a final concentration 200 mg L^-1^. Uracil- or arginine auxotrophic strains were supplemented with a separately sterilized solution of uracil (Ura) to a final concentration of 150 mg L^-1^ [45] or with L-arginine·HCl (Arg) to a final concentration of 450 mg L^-1^ of L-arginine [46]. For cloning, YPD medium, containing 10 g L^−1^ Bacto Yeast extract, 20 g L^−1^ Bacto Peptone and 20 g L^-1^ glucose was used, and when necessary supplemented with 200 mg L^−1^ G418 (Geneticin, Invivogen) or 200 mg L^−1^ Hygromycin. For solid media 2 % (w/v) Bacto™ Agar (Bacton Dickinson (BD), Franklin lakes, NJ, USA) was added to the medium prior to heat sterilization. Single-colony isolates were obtained by re-streaking single colonies three consecutive times on selective medium. Culture densities at any point of this study were determined using a Jenway™ 7200 spectrometer (Cole-Parmer, Staffordshire, UK) measuring the optical density at 660 nm (OD_660_) with a 1 cm light path.

**Table 1.**
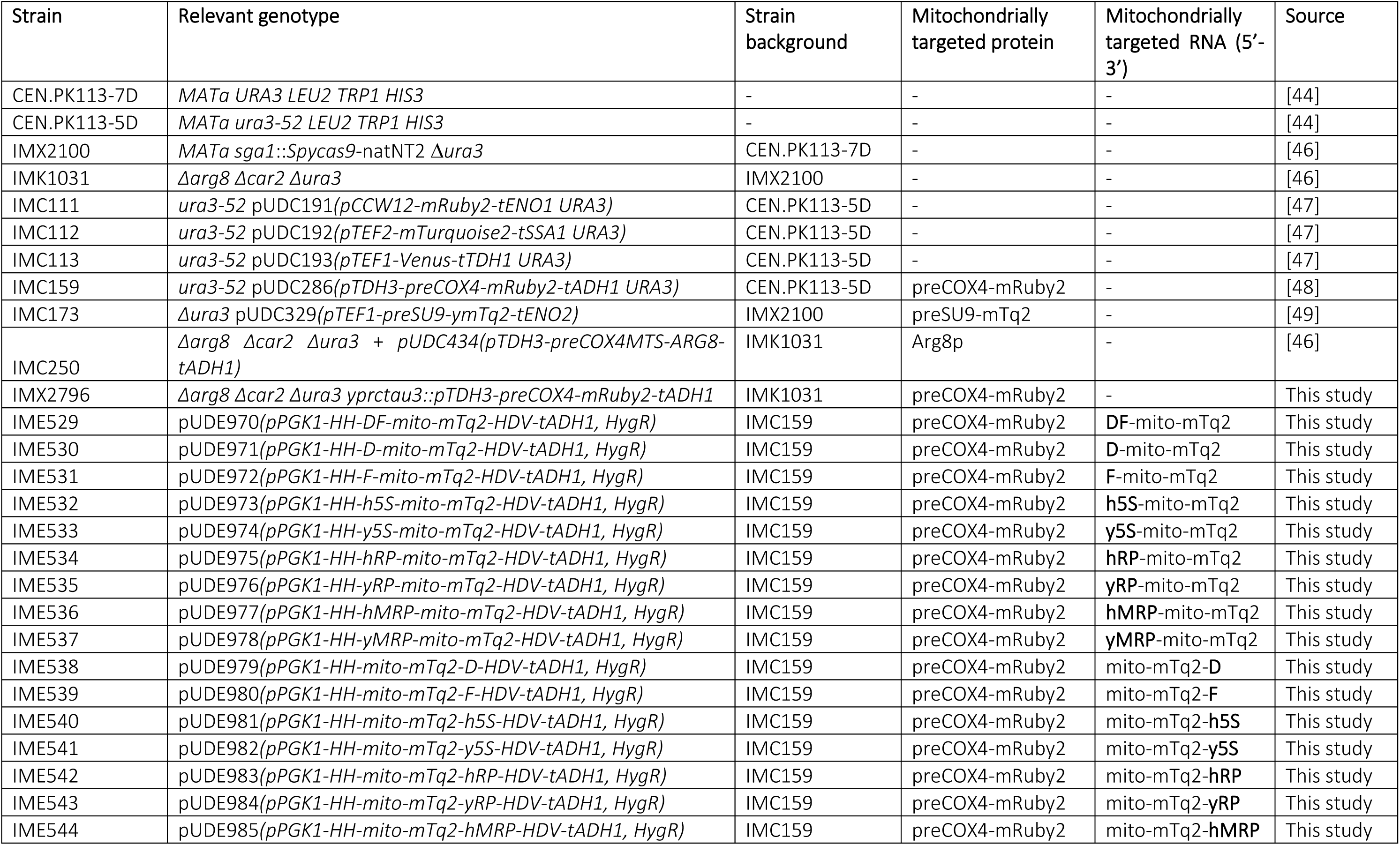
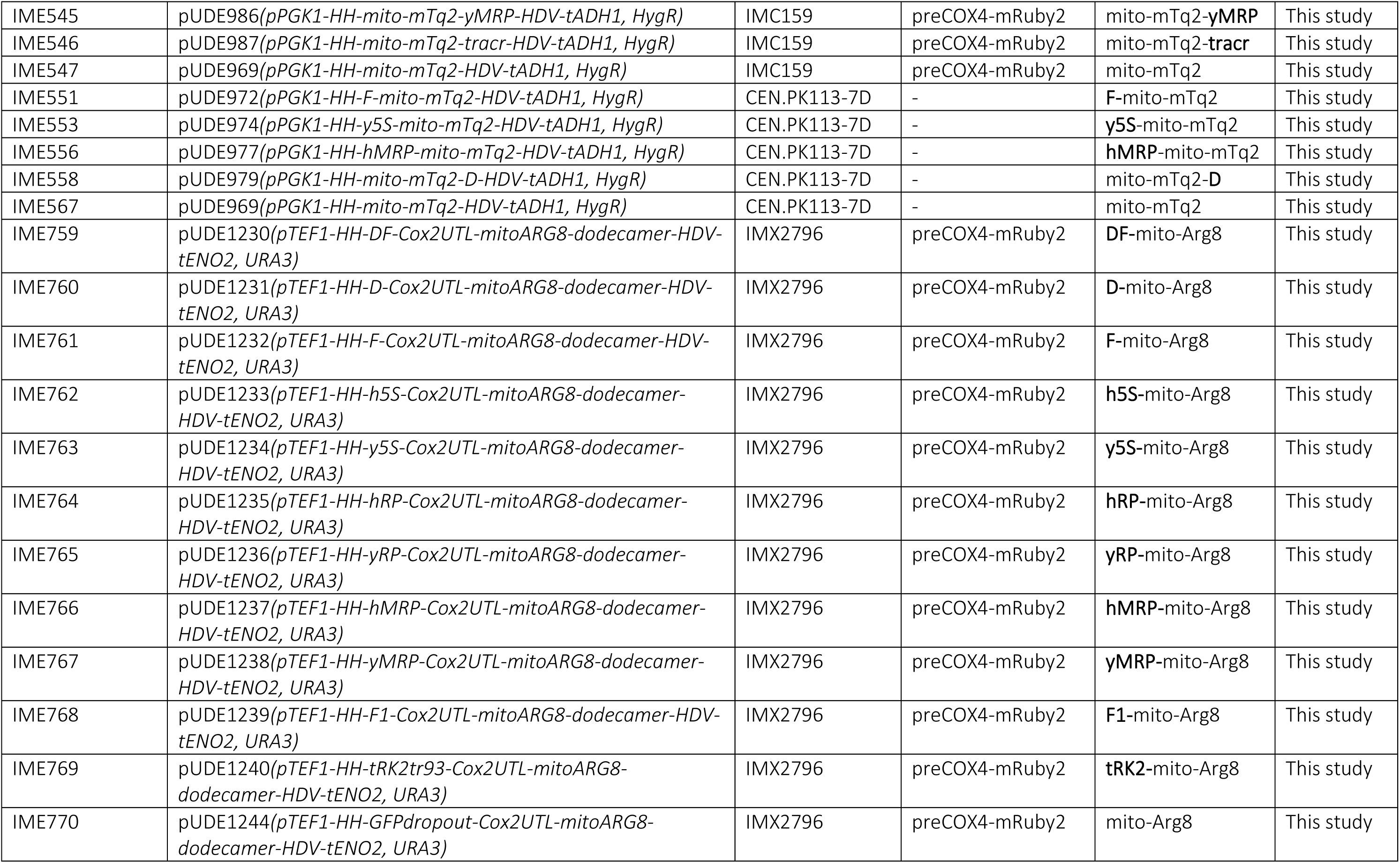
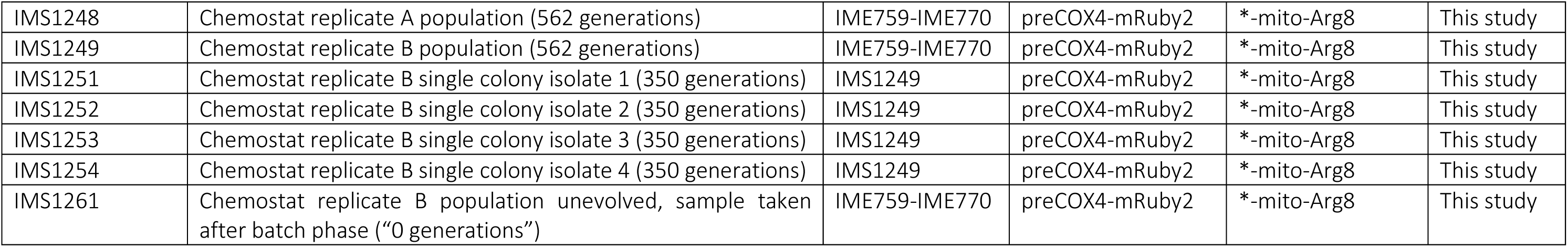
Strains used in this study. The genotype of plasmids is shown between brackets, RNA-targeting signals are indicated in bold. * Indicates a mix of targeting signals.

| Strain | Relevant genotype | Strain background | Mitochondrially targeted protein | Mitochondrially targeted RNA (5'-3') | Source |
| --- | --- | --- | --- | --- | --- |
| CEN.PK113-7D | *MATa URA3 LEU2 TRP1 HIS3* | - | - | - | [44] |
| CEN.PK113-5D | *MATa ura3-52 LEU2 TRP1 HIS3* | - | - | - | [44] |
| IMX2100 | *MATa sga1::Spycas9-natNT2 Δura3* | CEN.PK113-7D | - | - | [46] |
| IMK1031 | *Δarg8 Δcar2 Δura3* | IMX2100 | - | - | [46] |
| IMC111 | *ura3-52 pUDC191(pCCW12-mRuby2-tENO1 URA3)* | CEN.PK113-5D | - | - | [47] |
| IMC112 | *ura3-52 pUDC192(pTEF2-mTurquoise2-tSSA1 URA3)* | CEN.PK113-5D | - | - | [47] |
| IMC113 | *ura3-52 pUDC193(pTEF1-Venus-tTDH1 URA3)* | CEN.PK113-5D | - | - | [47] |
| IMC159 | *ura3-52 pUDC286(pTDH3-preCOX4-mRuby2-tADH1 URA3)* | CEN.PK113-5D | preCOX4-mRuby2 | - | [48] |
| IMC173 | *Δura3 pUDC329(pTEF1-preSU9-ymTq2-tENO2)* | IMX2100 | preSU9-mTq2 | - | [49] |
| IMC250 | *Δarg8 Δcar2 Δura3 + pUDC434(pTDH3-preCOX4MTS-ARG8-tADH1)* | IMK1031 | Arg8p | - | [46] |
| IMX2796 | *Δarg8 Δcar2 Δura3 yprctau3::pTDH3-preCOX4-mRuby2-tADH1* | IMK1031 | preCOX4-mRuby2 | - | This study |
| IME529 | *pUDE970(pPGK1-HH-DF-mito-mTq2-HDV-tADH1, HygR)* | IMC159 | preCOX4-mRuby2 | **DF**-mito-mTq2 | This study |
| IME530 | *pUDE971(pPGK1-HH-D-mito-mTq2-HDV-tADH1, HygR)* | IMC159 | preCOX4-mRuby2 | **D**-mito-mTq2 | This study |
| IME531 | *pUDE972(pPGK1-HH-F-mito-mTq2-HDV-tADH1, HygR)* | IMC159 | preCOX4-mRuby2 | **F**-mito-mTq2 | This study |
| IME532 | *pUDE973(pPGK1-HH-h5S-mito-mTq2-HDV-tADH1, HygR)* | IMC159 | preCOX4-mRuby2 | **h5S**-mito-mTq2 | This study |
| IME533 | *pUDE974(pPGK1-HH-y5S-mito-mTq2-HDV-tADH1, HygR)* | IMC159 | preCOX4-mRuby2 | **y5S**-mito-mTq2 | This study |
| IME534 | *pUDE975(pPGK1-HH-hRP-mito-mTq2-HDV-tADH1, HygR)* | IMC159 | preCOX4-mRuby2 | **hRP**-mito-mTq2 | This study |
| IME535 | *pUDE976(pPGK1-HH-yRP-mito-mTq2-HDV-tADH1, HygR)* | IMC159 | preCOX4-mRuby2 | **yRP**-mito-mTq2 | This study |
| IME536 | *pUDE977(pPGK1-HH-hMRP-mito-mTq2-HDV-tADH1, HygR)* | IMC159 | preCOX4-mRuby2 | **hMRP**-mito-mTq2 | This study |
| IME537 | *pUDE978(pPGK1-HH-yMRP-mito-mTq2-HDV-tADH1, HygR)* | IMC159 | preCOX4-mRuby2 | **yMRP**-mito-mTq2 | This study |
| IME538 | *pUDE979(pPGK1-HH-mito-mTq2-D-HDV-tADH1, HygR)* | IMC159 | preCOX4-mRuby2 | mito-mTq2-**D** | This study |
| IME539 | *pUDE980(pPGK1-HH-mito-mTq2-F-HDV-tADH1, HygR)* | IMC159 | preCOX4-mRuby2 | mito-mTq2-**F** | This study |
| IME540 | *pUDE981(pPGK1-HH-mito-mTq2-h5S-HDV-tADH1, HygR)* | IMC159 | preCOX4-mRuby2 | mito-mTq2-**h5S** | This study |
| IME541 | *pUDE982(pPGK1-HH-mito-mTq2-y5S-HDV-tADH1, HygR)* | IMC159 | preCOX4-mRuby2 | mito-mTq2-**y5S** | This study |
| IME542 | *pUDE983(pPGK1-HH-mito-mTq2-hRP-HDV-tADH1, HygR)* | IMC159 | preCOX4-mRuby2 | mito-mTq2-**hRP** | This study |
| IME543 | *pUDE984(pPGK1-HH-mito-mTq2-yRP-HDV-tADH1, HygR)* | IMC159 | preCOX4-mRuby2 | mito-mTq2-**yRP** | This study |
| IME544 | *pUDE985(pPGK1-HH-mito-mTq2-hMRP-HDV-tADH1, HygR)* | IMC159 | preCOX4-mRuby2 | mito-mTq2-**hMRP** | This study |
| IME545 | pUDE986(*pPGK1-HH-mito-mTq2-yMRP-HDV-tADH1, HygR*) | IMC159 | preCOX4-mRuby2 | mito-mTq2-**yMRP** | This study |
| IME546 | pUDE987(*pPGK1-HH-mito-mTq2-tracr-HDV-tADH1, HygR*) | IMC159 | preCOX4-mRuby2 | mito-mTq2-**tracr** | This study |
| IME547 | pUDE969(*pPGK1-HH-mito-mTq2-HDV-tADH1, HygR*) | IMC159 | preCOX4-mRuby2 | mito-mTq2 | This study |
| IME551 | pUDE972(*pPGK1-HH-F-mito-mTq2-HDV-tADH1, HygR*) | CEN.PK113-7D | - | **F**-mito-mTq2 | This study |
| IME553 | pUDE974(*pPGK1-HH-y5S-mito-mTq2-HDV-tADH1, HygR*) | CEN.PK113-7D | - | **y5S**-mito-mTq2 | This study |
| IME556 | pUDE977(*pPGK1-HH-hMRP-mito-mTq2-HDV-tADH1, HygR*) | CEN.PK113-7D | - | **hMRP**-mito-mTq2 | This study |
| IME558 | pUDE979(*pPGK1-HH-mito-mTq2-D-HDV-tADH1, HygR*) | CEN.PK113-7D | - | mito-mTq2-**D** | This study |
| IME567 | pUDE969(*pPGK1-HH-mito-mTq2-HDV-tADH1, HygR*) | CEN.PK113-7D | - | mito-mTq2 | This study |
| IME759 | pUDE1230(*pTEF1-HH-DF-Cox2UTL-mitoARG8-dodecamer-HDV-tENO2, URA3*) | IMX2796 | preCOX4-mRuby2 | **DF**-mito-Arg8 | This study |
| IME760 | pUDE1231(*pTEF1-HH-D-Cox2UTL-mitoARG8-dodecamer-HDV-tENO2, URA3*) | IMX2796 | preCOX4-mRuby2 | **D**-mito-Arg8 | This study |
| IME761 | pUDE1232(*pTEF1-HH-F-Cox2UTL-mitoARG8-dodecamer-HDV-tENO2, URA3*) | IMX2796 | preCOX4-mRuby2 | **F**-mito-Arg8 | This study |
| IME762 | pUDE1233(*pTEF1-HH-h5S-Cox2UTL-mitoARG8-dodecamer-HDV-tENO2, URA3*) | IMX2796 | preCOX4-mRuby2 | **h5S**-mito-Arg8 | This study |
| IME763 | pUDE1234(*pTEF1-HH-y5S-Cox2UTL-mitoARG8-dodecamer-HDV-tENO2, URA3*) | IMX2796 | preCOX4-mRuby2 | **y5S**-mito-Arg8 | This study |
| IME764 | pUDE1235(*pTEF1-HH-hRP-Cox2UTL-mitoARG8-dodecamer-HDV-tENO2, URA3*) | IMX2796 | preCOX4-mRuby2 | **hRP**-mito-Arg8 | This study |
| IME765 | pUDE1236(*pTEF1-HH-yRP-Cox2UTL-mitoARG8-dodecamer-HDV-tENO2, URA3*) | IMX2796 | preCOX4-mRuby2 | **yRP**-mito-Arg8 | This study |
| IME766 | pUDE1237(*pTEF1-HH-hMRP-Cox2UTL-mitoARG8-dodecamer-HDV-tENO2, URA3*) | IMX2796 | preCOX4-mRuby2 | **hMRP**-mito-Arg8 | This study |
| IME767 | pUDE1238(*pTEF1-HH-yMRP-Cox2UTL-mitoARG8-dodecamer-HDV-tENO2, URA3*) | IMX2796 | preCOX4-mRuby2 | **yMRP**-mito-Arg8 | This study |
| IME768 | pUDE1239(*pTEF1-HH-F1-Cox2UTL-mitoARG8-dodecamer-HDV-tENO2, URA3*) | IMX2796 | preCOX4-mRuby2 | **F1**-mito-Arg8 | This study |
| IME769 | pUDE1240(*pTEF1-HH-tRK2tr93-Cox2UTL-mitoARG8-dodecamer-HDV-tENO2, URA3*) | IMX2796 | preCOX4-mRuby2 | **tRK2**-mito-Arg8 | This study |
| IME770 | pUDE1244(*pTEF1-HH-GFPdropout-Cox2UTL-mitoARG8-dodecamer-HDV-tENO2, URA3*) | IMX2796 | preCOX4-mRuby2 | mito-Arg8 | This study |
| IMS1248 | Chemostat replicate A population (562 generations) | IME759-IME770 | preCOX4-mRuby2 | *-mito-Arg8 | This study |
| IMS1249 | Chemostat replicate B population (562 generations) | IME759-IME770 | preCOX4-mRuby2 | *-mito-Arg8 | This study |
| IMS1251 | Chemostat replicate B single colony isolate 1 (350 generations) | IMS1249 | preCOX4-mRuby2 | *-mito-Arg8 | This study |
| IMS1252 | Chemostat replicate B single colony isolate 2 (350 generations) | IMS1249 | preCOX4-mRuby2 | *-mito-Arg8 | This study |
| IMS1253 | Chemostat replicate B single colony isolate 3 (350 generations) | IMS1249 | preCOX4-mRuby2 | *-mito-Arg8 | This study |
| IMS1254 | Chemostat replicate B single colony isolate 4 (350 generations) | IMS1249 | preCOX4-mRuby2 | *-mito-Arg8 | This study |
| IMS1261 | Chemostat replicate B population unevolved, sample taken after batch phase ("0 generations") | IME759-IME770 | preCOX4-mRuby2 | *-mito-Arg8 | This study |

*Escherichia coli* XL1-Blue Subcloning Grade Competent Cells (Agilent Genomics, Santa Clara, US) were used for molecular cloning purposes and plasmid proliferation. Bacterial cultures were grown in Lysogeny Broth (LB) media (10 g·L^−1^ Bacto™ Tryptone, 5 g·L^−1^ Bacto™ Yeast Extract (BD), 5 g·L^−1^ NaCl). LB media were autoclaved at 121 °C for 20 minutes. Where relevant, media were supplemented with ampicillin or chloramphenicol to a final concentration 10 mg L^-1^. 5 mL liquid cultures were grown in a 15 mL CELLSTAR® tube (Greiner Bio-One, Kremsmünster, Austria) at 37 °C while shaking 200 rpm in an Innova 4000 Incubator Shaker (Eppendorf).

Frozen stocks were prepared by addition of sterile glycerol (30% v/v) to late exponential phase shake-flask cultures of *S. cerevisiae* or *E. coli* and 1 mL aliquots were stored aseptically at −80 °C.

### General molecular biology techniques

Golden Gate assembly was done with 30 fmol of each fragment and 10 fmol of backbone. Additionally, the reaction mixtures contained 0.5 µL BsmbI or BsaI-HF®v2 (New England Biolabs (NEB), Ipswich, MA, USA); 0.5 µL T7 DNA ligase (NEB); 1 µL T4 DNA ligase buffer (NEB) and was filled to 10 µL with nuclease-free water. When cloning with BsmbI or BsaI-HF®v2, respective digestion temperatures of 42 °C and 37.5 °C were used. Ligation steps were always performed at 16 °C. The Golden Gate protocol used consisted of 25 cycles of 2 minutes digestion and 5 minutes ligation, followed by a final digestion step at 60 °C and a heat inactivation step at 80 °C, each lasting 10 minutes. Gibson isothermal assembly was performed using NEBuilder HiFi DNA Assembly Master Mix (NEB) with 10 fmol of backbone and 20 fmol of insert, and incubated at 50 °C for 1 hour. High fidelity PCR reactions were executed with Phusion® Hot Start II High Fidelity Polymerase (Thermo Fisher Scientific) and PAGE-purified primers (Sigma-Aldrich, St. Louis, MO, USA) according to manufacturer’s instructions. Gibson assembly products and Golden Gate assembly products were heat-shock transformed to chemical competent *E. coli* XL1-Blue cells, prepared following the supplier’s instructions (Agilent Genomics). Plasmids were harvested from overnight bacterial cultures using GeneJET Plasmid Miniprep Kit (Thermo Fisher Scientific) according to the manufacturer’s instructions. Fidelity of plasmids was assessed through diagnostic PCR reactions, done using DreamTaq Master Mix (Thermo Fisher Scientific) and desalted primers (Sigma-Aldrich, Table S2) according to manufacturer’s instructions. Plasmids were transformed into yeast following the LiAc/SS carrier DNA/PEG method [50].

### Plasmid and strain construction

#### Plasmids and strains expressing mito-mTq2 mRNA

Plasmids pUDE969 – pUDE987 expressing mito-mTq2 mRNA were cloned using Golden Gate assembly according to the Yeast Toolkit (YTK) design of Lee, DeLoache (51). An extensive cloning strategy including primer numbers is depicted in Figure S 1. A list of plasmids and part plasmids is described in Table S1. First, all RNA targeting signals were ordered as complementary ssDNA primer pairs containing YTK-type 3a BsaI flanks for 5’signals or type YTK-type 4a BsaI flanks for 3’signals respectively, their sequences are listed in Table S2. *In vitro* annealing of the signals was done by heating a 1:1 ratio of complementary ssDNA oligos to 100 °C for 5 minutes and letting it cool down on the bench at room temperature. Recoded mito-mTq2 (see Supplementary information for sequence) was ordered as a synthetic gene fragment at GeneArt (Thermo Fisher Scientific, Landsmeer, The Netherlands), and was amplified with primers containing YTK-type 3b BsaI flanks for 5’signals or YTK-type 3 BsaI flanks for 3’ or no signals respectively (Table S2). All parts also contained BsmBI flanks which were used to clone the signals as part plasmids in backbone pYTK001 using Golden Gate cloning as described above, yielding part plasmids pGGKp265-pGGKp291 [51](Table S 1). Backbone pUDE929 (*pPGK1-HH-dKanMX-HDV-tADH1, bla, ColE1, HygR,2µ)* was linearized by PCR-amplification with primers annealing to the HH and HDV with YTK-type 3 and 4(b) BsaI flanks, respectively (detailed construction strategy is shown in Figure S 1). The different part plasmids with import signals and/or mito-mTq2 (pGGKp265 - pGGKp291) were then assembled in linearized backbone pUDE929 using Golden Gate assembly, yielding plasmids pUDE969 – pUDE987 (Table S 1). For co-expression with mRuby2, plasmids were transformed in yeast strain IMC159 [48], yielding strains IME529 – IME547. For sole expression of mito-mTq2 mRNA, plasmids were transformed in CEN.PK113-5D, yielding strains IME551 – IME567. For co-expression with preCOX4-Arg8, the plasmids were transformed in strain IMC250 [46], yielding strains IME797 – IME802.

#### Plasmids and strains expressing mito-ARG8 mRNA

The plasmids expressing mito-ARG8 mRNA (pUDE1230 - pUDE1241) were constructed from the construct pUD1202 (*HH-GFPdropout-Cox2UTL-mitoARG8-dodecamer-HDV),* which was ordered as a synthetic gene fragment at GeneArt (Thermo Fischer Scientific), and the fragment was PCR amplified with primers annealing to the HH/HDV sequences of the fragment. A backbone was PCR-amplified from plasmid pUDC329. using primers annealing to the promoter and terminator with homology flanks to the HH/HDV ribozyme sequences, respectively (Figure S 1). The fragment and the backbone were cloned together by Gibson isothermal assembly, resulting in plasmid pUDC365. The full construct including promoter and terminator (*pTEF1*-*HH-GFPdropout-Cox2UTL-mitoARG8-dodecamer-HDV-tENO2*) was amplified and inserted via Gibson assembly into a PCR-amplified backbone of pUD518, which is a multi-copy vector, yielding plasmid pUDE1241 (Figure S1). The 5’ signals were subsequently cloned into the GFP-dropout site of the plasmid as described by Lee, DeLoache (51) to generate a library of mito-Arg8 mRNA with different signals (plasmids pUDE1230 - pUDE1240, Figure S1,Table S1). Plasmids were transformed into yeast yielding strains IME759 – IME770 (Table 1).

#### Genomic integration

For genomic integration of the preCOX4-mRuby2 in the YPRCtau3 locus in strain IMX2796, the preCOX4-mRuby expression cassette repair fragment was amplified from plasmid pUDC286 using high-fidelity PCR with primers that attached 60 bp homology flanks of the YPRCtau3 locus of CEN.PK113-7D to the repair fragment (Table S1,Table S2). The repair fragment was co-transformed with plasmid pUDR514 expressing two gRNA sequences targeting the YPRCtau3 locus in *SpyCas9*-expressing strain IMK1031 as described [52, 53], following the LiAc/SS carrier DNA/PEG method.

### mRNA-FISH

Probe sets for single-molecule RNA FISH were designed using Stellaris® Probe Designer (version 4.2; Biosearch™ Technologies, Inc., Petaluma, CA). The sets targeting mito-mTq2 mRNA and *COX3* mRNA respectively consisted out of 25 Quasar-570® labelled probes and 24 CAL Fluor® Red 635 labelled probes. All probes were 22-mers, designed to anneal with a minimal spacing of 1 nt. Probes were diluted and stored as instructed by the manufacturer.

The protocol for RNA FISH was adopted from Schwabe and Bruggeman (54) with the following modifications. 20 mL starter cultures (SMEG + HygB) were inoculated and after 6-8 hours and transferred to 100 mL fresh medium. Approximately 1·10^9^ cells were sampled from exponentially growing cultures and fixed in 4% (w/v) paraformaldehyde in SM. Formaldehyde fixation was done while shaking 200 rpm at 30 °C for at least 30 minutes, but for no longer than 1 hour. Fixed cells were stored at least 1 night at 4 °C. All consecutive steps entailed RNase-free work, for which a separate bench was dedicated for work with ambidextrous glove protection. Where possible, chemicals and consumables were autoclaved at 121 °C for at least 45 minutes or otherwise treated with RNase*Zap* decontamination solution (Sigma-Aldrich). Prior to spheroplasting, cells were washed twice with cooled spheroplasting buffer (1.2 M sorbitol, 0.1 M pH 7.5 potassium phosphate buffer (KPB)). Cells were resuspended in 1.25 mL spheroplasting buffer containing approximately 800 U lyticase from *Arthrobacter luteus* (L4025-50KU; Sigma-Aldrich) and incubated at 30 °C while shaking gently. Progression of spheroplast formation was monitored by phase-contrast microscopy, aiming for roughly 50% of the cells to appear phase-contrast dark (i.e. after 50 – 70 min). At any succeeding point of the protocol, centrifugation of spheroplasts was done at 380 × *g* for 5 minutes at room temperature and resuspension of spheroplasts was done by gently flicking the tube. The spheroplasting reaction was terminated by 2 cycles of spinning down the spheroplasts and washing with 1 mL cold spheroplasting buffer. Obtained 1 mL suspensions were distributed in 200 µL aliquots and pelleted. Spheroplasts were resuspended in 200 µL 70% EtOH for storage at 4 °C until further use. Prior to hybridization, spheroplasts were spun down, ethanol supernatants were aspirated, and pellets were resuspended in 1 mL room-temperature (RT) wash buffer (2x saline sodium citrate (SSC) buffer, 10% [v/v] formamide). After a 5-minute incubation on bench, samples were centrifuged, aspirated, and resuspended in 50 µL hybridization solution (100 g·L^-1^ dextran sulfate sodium salt from *Leuconostoc* spp., 1 g·L^-1^ *E. coli* tRNA, 2 mM vanadyl ribonucleoside complex (VRC), 200 mg·L^-1^ bovine serum albumin (BSA), 2x SSC buffer, 10% formamide, 125 nM probe) at RT. Hybridization was done overnight, protected from light, in a heat block set at 30 °C and 300 rpm. After hybridization, spheroplasts were washed with 1 mL wash buffer at RT, centrifuged and aspirated. Subsequently, pellets were resuspended in 1 mL wash buffer (RT) containing 500 ng·mL^-1^ DAPI (Sigma-Aldrich) and incubated for 30 minutes at 30 °C, after which the samples were pelleted and aspirated. Spheroplasts were resuspended in 1 mL wash buffer and incubated for 30 minutes at 30 °C.

Spheroplasts were resuspended in 100 µL gelvatol (155 g·L^-1^ polyvinyl alcohol [PVA], 21% [v/v] glycerol, 53% [v/v] Tris buffer (0.2 M Trizma® base (Sigma-Aldrich), pH 8.5), a few crystals of sodium azide) as a mounting agent. 5 µL of this suspension was placed on a SuperFrost objective slide (Menzel-Gläzer, Thermo Fisher Scientific) and covered with a #1.5 high precision cover slip (Paul Marienfeld & Co., Lauda-Köningshofen, Germany). Coverslips were sealed to objective slides using transparent nail polish (product no. 11244545; HEMA, Amsterdam, The Netherlands). Slides were labelled and stored at room temperature while protected from light until required for microscopy.

### Microscopy

Brightfield, phase-contrast and fluorescence widefield microscopy were done using Zeiss Axio Imager Z1 Upright Microscope (Carl Zeiss, Oberkochen, Germany), equipped with a HAL 100 Halogen Illuminator (Carl Zeiss) and an HBO 100 illuminating system (Carl Zeiss). Acquisitions were done with an AxioCam HRm Rev3 detector (60N-C 1’’ 1.0x) (Carl Zeiss), connected to Zen 2 v10.0.0.910 (Carl Zeiss) software. For live-cell microscopy, strains were grown as a pre-culture overnight at 30 °C, in shake flasks containing SMD supplied with appropriate antibiotics. The next morning, cultures were diluted 1:50 in a fresh shake flask with SMD and grown for another 3-4 hours to an OD of approximately 1-2, to ensure strains were in a mid-exponential growth phase. 1 mL from a liquid culture was centrifuged for 3 minutes 3000 × *g* at RT, re-suspended in sterile dH_2_O and kept on ice until imaging. 4 µL was imaged using a 100x EC Plan-NeoFLUAR lateral magnification oil-immersion objective with a numerical aperture (NA) of 1.3 (Carl Zeiss) after oil immersion with Immersol™ 518F type F immersion oil (Carl Zeiss). mTurquoise fluorescence was detected using filter set 47 (Carl Zeiss AG; excitation bandpass (BP) filter 436/20 nm, beam splitter filter 455 nm, emission filter BP 480/40 nm, Figure S2). The exposure time was always set at 1200 ms for all strains expressing mito-mTq2 mRNA. mRuby2 fluorescence was detected using filter set 14 (Carl Zeiss, excitation BP 535/25, emission Long Pass (LP) 590, Figure S2), with an exposure time of 500 ms. Exposure times were set such that the red fluorescence was clearly visible, and no blue fluorescence (caused by i.e. auto-fluorescence or overexposure of the cells) was observed in a negative control strain not expressing mTurquoise2.

For image acquisition of prepared slides as described for FISH experiments, fluorescence widefield microscopy was performed using an Olympus IX 81 inverted microscope (Olympus, Tokyo, Japan), equipped with an Andor™ AMH-200-F6S high-power metal halide lamp (Oxford Instruments, Abingdon, UK) and an Andor™ Luca R EM-CCD camera (Oxford Instruments) connected to Andor™ iQ3 control and acquisition software (Oxford Instruments). Acquisitions were done using a 100x UplanFLN lateral magnification objective with an NA of 1.3 after oil immersion with IMMOIL-F30CC low auto-fluorescence type F immersion oil (Olympus).

### Image analysis

Image analysis was performed using the FIJI package of ImageJ [55]. Mean co-localized fluorescence of mRuby and mTurquoise2 in the mitochondria was analysed by thresholding the red fluorescence channel of each image using the minimal algorithm to determine the location of the mitochondria in the image. Based on the thresholded red fluorescence image, a binary mask was generated indicating the regions of interest (ROI, i.e. mitochondrial structures) of the image. Subsequently, the mask was overlaid on the red and blue fluorescent channels of the image and the mean fluorescence for each ROI was determined using FIJI, resulting in a mean mitochondrial fluorescence per cell in both the red and blue channel. ANOVA one-way analysis were calculated using R-based analysis in RStudio (RStudio, Boston, MA, USA) using the ggpubr package of R.

When there was no fluorescent protein targeted to the mitochondrial matrix, masking was on the brightfield channel of the microscopy data. The cell boundary was determined in the brightfield image using auto-thresholding with the Otsu algorithm. The background value was determined by measuring the fluorescence in the non-masked area (i.e. without cells) Subsequently, within the cell area, the background value was subtracted and the mean fluorescence in the whole cell was determined. For strains expressing mito-mTq2 mRNA in combination with preCox4-Arg8, a fluorescence threshold was set at 1500, as this was found to be the value above which auto-fluorescence could not be detected in the negative controls.

FISH images were subject to deconvolution. The generation of Point Spread Functions (PSFs) for spectral deconvolution was done in ImageJ, using the Parallel Spectral Deconvolution (v1.9) plugin. All settings were kept constant, except for stack height in number of slices, spacing between slices in nm and emission wavelength (λ_EM_) of each registered channel. For each of these variations, a new PSF needed to be generated to subsequently deconvolve the image accordingly. All images were normalized by setting the sum of pixels to 1. Algorithmic deconvolution was done in ImageJ using the Deconvolutionlab2 (v2.1.2) plugin. Each channel was deconvolved in grey-scale separately and finally combined to a multi-channel, multidimensional acquisition. Deconvolution was done using Richardson-Lucy deconvolution, commonly used for deconvolution when the PSF is known, but no or little information is known about the noise. All deconvolutions for this study were done with 35 iterations.

### Growth characterization of mito-ARG8 mRNA mutants

Growth rate analysis of mutants expressing mito-ARG8 mRNA was performed in 96-wells microtiter plates at 30°C and 250 rpm using a Growth Profiler 960 (EnzyScreen BV, Heemstede, The Netherlands), essentially as described previously [46]. Frozen glycerol stocks were inoculated in 100 mL SMD + Arg and grown overnight. Cultures were supplied with uracil when necessary. 0.5 mL of the overnight culture was transferred to 100 mL SMD + Arg and grown until the OD_660_ had doubled at least once to ensure exponential growth. The cultures were spun down and washed twice in sterile dH_2_O, then diluted to an OD_660_ of 15. A 96-wells microtiter plate (EnzyScreen, type CR1496dl) containing SMD or SMEG with final working volumes of 250 μL was inoculated with a starting OD_660_ of 0.3, approximating 300 cells. Growth rate analysis and the calculation of OD equivalents was performed as described by Boonekamp, Knibbe (56).

### Laboratory evolution and bioreactor cultivation

#### Chemostat cultures

Pre-cultures of IME759 – IME770 strains expressing mito-ARG8 mRNA and preCOX4-mRuby2 were grown overnight in 20 mL volume of SMD + Arg in 100 mL shake flasks. A total of 20 OD_660_ units of each strain was combined in a single 50 mL centrifuge tube so an equal number of cells per strains would be inoculated in each bioreactor. The combined cultures were inoculated for automated evolutionary engineering in duplicate parallel 2 L Applikon Bio laboratory bioreactors (Getinge, Delft, the Netherlands) with a 1 L working volume [57], containing arginine-limited SME (SM supplemented with 20 g L^-1^ ethanol, 0.3 g L^-1^ antifoam Pluronic PE 6100 (BASF, Ludwigshafen, Germany) and 60 mg L^-1^ L-Arginine). Cultures were stirred at 800 RPM, the temperature was controlled at 30 °C, and the pH was maintained at 5.0 through automated addition of 2.0 M KOH. The cultures were sparged with air at a flow rate of 475 mL min^−1^. The strains were grown in batch until a stabilization of CO_2_ indicated arginine depletion. Upon arginine depletion, evolution in continuous culture set-up was started by switching on medium pumps to obtain a constant in-flow of arginine-limited SME. The culture volume was kept constant at 1 L using an effluent pump that was controlled by an electric level sensor, resulting in a stable dilution rate of 0.1 h^-1^. Evaporation of water and volatile metabolites was minimized by cooling the outlet gas of bioreactors to 4°C in a condenser. CO_2_ and O_2_ concentrations were measured continuously in in the reactor outlet gas which was dried with a PermaPure PD-50T-12MPP dryer (Permapure, Lakewood, NJ) prior to analysis, and concentrations in the outlet gas were measured with an NGA 2000 Rosemount gas analyzer (Emerson, St. Louis, MO) calibrated with reference gas containing 3.03% CO_2_ and N6-grade N_2_ (Linde Gas Benelux). All process parameters and output were continuously monitored *in situ* using Lucullus software (v3.8.2, Getinge) and through regular sampling and off-line measurements described below. Two independent bioreactors named A and B, corresponding to two evolution lines, were run in parallel. Glycerol stocks of the chemostat populations were taken at regular intervals at the same time as off-line measurements. The samples used for analysis were taken at the start of the chemostat phase right after the batch phase ended (IMS1261) and at the very end of the chemostat cultivation prior to ending the experiment (562 generations, IMS1248 and IMS1249). Single-colony isolates IMS1250 - IMS1254 were isolated from samples taken from bioreactor B after approximately 350 generations. To obtain the isolates, a sample of bioreactor B was plated on SME without arginine, and four colonies were picked and re-streaked on medium without arginine three consecutive times to obtain single-colony isolates.

#### Characterization of IMS1254 physiology in batch culture

Single-colony isolate IMS1254 was characterized in batch culture. The strain was grown overnight in shake flasks containing 100 mL SMD + Arg, washed with sterile dH_2_O and inoculated in three replicate bioreactors run in parallel, one containing SME supplemented with 20 g L^-1^ ethanol, and two containing SM supplemented with 20 g L^-1^ glucose. Aeration, pH, temperature, and dissolved oxygen concentration thresholds were the same as in the chemostat cultivations. The cultures were grown in batch mode until OD and CO_2_ stabilized to deplete any leftover arginine in the medium, upon which an empty-refill cycle with the respective medium (SMD or SME, omitting arginine) was started manually. The cultures were then characterized in batch mode by online monitoring of O_2_ and CO_2_ as described for chemostat cultures and by off-line measurements. After approximately 220 hours, the batch culture on SME was spiked with additional ethanol to a concentration of 20 g L^-1^ to ensure sufficient ethanol supply despite evaporation.

### Analytical methods for cultures in bioreactors

The bioreactors were sampled at regular intervals and several parameters were measured off-line. Biomass dry weight measurements of the bioreactor experiments were performed as previously described, using 10 mL of culture volume [58]. Metabolite concentrations in culture supernatants, obtained by centrifugation of 1 mL of culture for 3 min at 13.000 × *g,* were analyzed by high-performance liquid chromatography (HPLC) on an Agilent 1260 HPLC system (Agilent Technologies, Santa Clara, CA) fitted with a Bio-Rad HPX 87 H column (Bio-Rad, Hercules, CA) operated at 60 °C with 5 mM H_2_SO_4_ as the mobile phase with a flow rate of 0.6 mL min^-1^ and detection by refractive index and wavelength absorbance detectors at 214 nm. OD_660_ was determined on culture samples as described above. Colony forming units (CFU) were determined by making a 0 ×, 100 ×, 1000 × and 10.000 × serial dilutions of biomass samples in sterile dH_2_O and plating on solid SMD, SMD + Arg, SME and SME + Arg and counting CFU manually.

### Whole genome sequencing (WGS)

To obtain DNA for WGS, glycerol stocks from evolved strains IMS1251-1254 were grown overnight in 100 mL SMD + Arg to late exponential phase (OD_660_ > 10) and harvested by centrifugation. Biomass of evolved bioreactor populations IMS1248 and IMS1249 was harvested directly from the outflow of the bioreactors and spun down. Genomic DNA was extracted with the Qiagen Blood & Cell Culture DNA kit (Qiagen, Germantown, MD), following manufacturer’s specifications. WGS was performed by Macrogen Europe (Amsterdam, the Netherlands) on a Novaseq 6000 sequencer (Illumina, San Diego, CA) to obtain 151 cycle paired-end libraries with an insert-size of 550 bp using TruSeq Nano DNA library preparation, yielding 2 Gigabases in total per sample. All Illumina sequencing data are available at NCBI (https://www.ncbi.nlm.nih.gov/) under the Bioproject accession number PRJNA923502. Reads were mapped using BWA (version 0.7.15) [59] to a IMX2600 reference [60] Alignments were processed using SAMtools (version 1.3.1) [61], and variants were called by applying Pilon (version 1.18) [62]. Sequencing output was visualized using the Integrative Genomics Viewer [63] and chromosome copy numbers were determined using Magnolya [64].

### Mitochondria isolation

Mitochondria required for Western blotting or proteomics were isolated from shake-flask cultures. Glycerol stocks were grown overnight on SM medium supplemented glucose as a carbon source overnight, supplemented with HygB for strains expressing mito-mTq2 mRNA and supplemented with arginine for strains obtained from bioreactor experiments. The pre-cultures were transferred to fresh medium to an OD_660_ of 0.1 in a total of 300 mL of SMEG distributed over 3 shake flasks, supplied with arginine or HygB when required. The strains were then grown to mid-exponential phase (OD_660_ of 4-8) and 300 mL of culture were harvested by centrifugation. Mitochondrial isolation was performed as described in Koster, Kleefeldt (49). Mitochondria isolation was performed in biological duplicate experiments for each strain. The mitochondria-enriched fraction was immediately processed for Western blot analysis. For proteome analysis, the mitochondria were resuspended in 5 mL ice-cold Sorbitol-HEPES buffer (0.6 M sorbitol, 20 mM HEPES-KOH, 2 mM MgCl, cOmplete Protease Inhibitors Cocktail (Roche Diagnostics, Rotkreuz, Switzerland)), and distributed in 500 μL aliquots, equating approximately 1 -2 mg total protein consisting for 50-70 % of mitochondrial protein (Figure S13). The aliquots were flash-frozen in liquid nitrogen and stored at -80 °C for proteome analysis.

### Western blotting

For protein expression analysis by western blot, 50 μL of isolated mitochondria were resuspended in 200 µL ice-cold lysis buffer (50 mM HEPES pH 7.5, 150 mM NaCl, 2.5 mM EDTA, 1% v/v Triton X-100, cOmplete Mini Protease Inhibitors (Roche Diagnostics, Rotkreuz, Switzerland)), and thoroughly vortexed to release proteins. The supernatant was cleared by centrifugation (10.000 ×*g* for 10 min at 4 °C) and protein concentrations were determined using the Quickstart Bradford Protein Assay (Bio-rad Laboratories, Inc, Hercules, CA, USA). 25 µg of total protein were loaded on two separate 4-15% Mini-Protean TGX Stain-Free Protein gels (Bio-rad) at 250 V for 20 min. Gels were activated using the stain free protocol on a ChemiDoc MP Imager (Bio-rad) to verify separation of the protein extracts. Proteins were transferred to PVDF membranes using a Turboblotter system (Bio-rad). Successful transfer of proteins was verified using the ChemiDoc MP Imager. Membranes were blocked for 1 hour in TBS + 1% Casein (Blocking Buffer, Bio-rad) and washed once with PBSt (Phosphate saline buffer pH 7.2 with 0.05% Tween20). Washed membranes were incubated overnight, shaken at 4 °C, in PBSt. For detection of Subunit 3 of Cytochrome *c* oxidase, anti-Cox3p antibody (from mouse, #459300, Invitrogen, Waltham, MA, USA) was added to one of the membranes of each replicate, diluted 1 in 1000 in PBSt. After overnight incubation, membranes were washed in PBSt. Membranes with anti-Cox3p were incubated with the secondary anti-mouse HRP antibody, diluted 1 in 2000 (from goat, Dako Agilent, Santa Clara, CA, USA) for 3 hours at 4 °C. The other membranes were incubated for detection of mTurquoise2 with anti-GFP Alexa Fluor 488 conjugated antibody (from rabbit, #A21311, Invitrogen) for 1 h at room temperature. HRP antibody levels were detected with enhanced chemiluminescence using the Clarity Max Western ECL Substrate (Bio-rad) and chemiluminescence- and fluorescence signals were detected using a ChemiDoc MP Imager (Bio-Rad).

### Proteomic analysis

Proteome extraction and analysis was performed essentially as described in den Ridder, van den Brandeler (65), with modifications to accommodate proteome extraction of isolated mitochondria. Two biological replicates were used for each experiment. Mitochondrial aliquots (500μL, -80 °C) containing 1-2 mg protein consisting of 50-70 % mitochondrial protein (Figure S13), were thawed on ice and resuspended in lysis buffer composed of 100 mM Triethylammonium bicarbonate (TEAB) containing 1% SDS and phosphatase/protease inhibitors. Mitochondrial fractions were lysed by adding glass beads to the mitochondria and vortexing in 3 cycles of 1 minute, alternated with 1 min rest on ice. Proteins were reduced by addition of 5 mM DTT and incubation for 1 hour at 37°C. Subsequently, the proteins were alkylated for 60 min at room temperature in the dark by addition of 50 mM iodoacetamide. Protein precipitation was performed by addition of four volumes of ice-cold acetone (-20°C), followed by 1 hour freezing at -20°C. The proteins were solubilized using 100 mM ammonium bicarbonate. Proteolytic digestion was performed by Trypsin (Promega, Madison, WI), 1:100 enzyme to protein ratio, and incubated at 37°C overnight. Solid phase extraction was performed with an Oasis HLB 96-well μElution plate (Waters, Milford, USA) to desalt the mixture. Eluates were dried using a SpeedVac vacuum concentrator at 50 °C and frozen at -80 °C. An aliquot corresponding to approximately 1 µg protein digest was analyzed using a one-dimensional shot-gun proteomics approach in three technical replicate experiment for each biological replicate [65]. Data were analyzed against the proteome database from *Saccharomyces cerevisiae* (UniProt, strain ATCC 204508 / S288C, Tax ID: 559292, July 2020) using PEAKS Studio X (Bioinformatics Solutions Inc., Waterloo, Canada) as described [65]. The significance score for evaluating the observed abundance changes was calculated using a one-way ANOVA and expressed as the -10·*log*_10_(p), where p is the significance testing p-value, which represents the likelihood that the observed change is caused by random chance. Based on the average of the duplicate quantified proteins, the fold change of each protein in a specific condition was calculated relative to the unevolved sample. The average fold changes of the technical replicates were subsequently used to determine the standard deviations of the biological replicates. Results from the analysis are attached in SI file 2.

### GO term enrichment

A functional term enrichment analysis was performed to determine whether specific Gene Ontology (GO [66]) terms or were shared between proteins that were significantly more- or less abundant in two out of three or all three strains. GO-term analysis was performed using the GO::TermFinder [67] accessed through the GO Term Finder webpage hosted by the *Saccharomyces* genome data base (SGD, https://www.yeastgenome.org, [68]) using GO version 2023-01-01. A list of genes of interest (e.g. mitochondrial proteins more abundant in all strains), was analyzed against a custom reference list containing all *S. cerevisiae* mitochondrial proteins (SI file 2). The reference list of mitochondrial proteins was obtained by exporting a list of all genes with “mitochondrion” as cellular component (GO:0005739) from SGD. The enrichment strength was calculated by dividing the number of proteins in the entered dataset that associated with a GO-term by the number of proteins associated with the same GO-term in the reference list.

## Results

### Developing a fluorescence-based screening of RNA import in yeast mitochondria

An array of putative RNA structures of 20 – 80 nucleotides that aid import of RNA in mitochondria have been proposed previously [27, 31, 69]. A subset of these RNA structures was selected to test whether their proposed ability to import RNA molecules extends to longer RNA molecules (Table 2). Four tRNA-derived import signals (DF, D-arm, F-stem, F1-stem) identified in *S. cerevisiae* were selected based on their described ability to effectively import RNA in yeast mitochondria. Additionally, three signals described to aid import in human mitochondria (5s rRNA, RNase P and RNase MRP) were tested, as well as their three yeast homologues (Table 2, Figure S3). Lastly, since a tracrRNA structure was also described to aid RNA import [31], this RNA structure was also included, resulting in a library of eleven import signals, that could be cloned at either the 5’ end or the 3’ end of an mRNA molecule.

The RNA import capacity of eleven RNA structures was first investigated by a fluorescence-based screening approach (Figure 1), relying on the alternative codon usage of yeast mitochondria. In mitochondria, the UGA codon encodes a tryptophan (Trp) while it encodes a STOP codon when translated in the cytosol [70]. Nuclear gene expression with the UGA codon ensures that full length mRNAs are only translated in the mitochondria, while translation in the cytosol yields truncated proteins. A 720-nucleotide mRNA sequence encoding the blue fluorescent protein mTurquoise2 (mTq2) was chosen as mRNA molecule for mitochondrial import, due to its bright fluorescence, stability, short maturation time and presence of two Trp-residues in the amino acid sequence [71]. The mTq2 mRNA was recoded for mitochondrial translation by replacing these two native Trp-codons with the UGA codon. Subsequently, translation of the recoded mTq2 mRNA (referred to as mito-mTq2 mRNA) in mitochondria would result in a 238 amino acid-long fluorescent protein, while cytosolic translation would yield truncated, non-fluorescent proteins of 65 and 161 amino acids (Figure S4). The selected RNA import signals were cloned at either the 5’ or 3’ end of the recoded mito-mTq2 mRNA in a multicopy plasmid under control of a strong constitutive promoter. This resulted in eighteen different strains, each expressing mito-mTq2 with a putative RNA import signal attached at either the 5’ or the 3’-end. A control omitting an import signal was included, leading to a total of nineteen plasmids (pUDE969-987, Table S 1). To prevent potential interference with mitochondrial translation, the 5’-cap and poly(A)-tail added during transcription in the nucleus were removed by framing the 5’ and 3’ of the RNA construct with self-cleaving ribozymes (Figure S1) [72, 73]. Constitutive fluorescent labelling of mitochondria was achieved by targeting the cytosolically translated red fluorescent protein mRuby2 to mitochondria (preCOX4-mRuby2, Figure 1). The nineteen mito-mTq2 mRNA plasmids were expressed in the presence (strains IME529 – IME546) or absence of preCOX4-mRuby2 (strains IME551 – IME567), and localization of the red and blue fluorescent proteins was assessed by fluorescence microscopy.

**Table 2.**
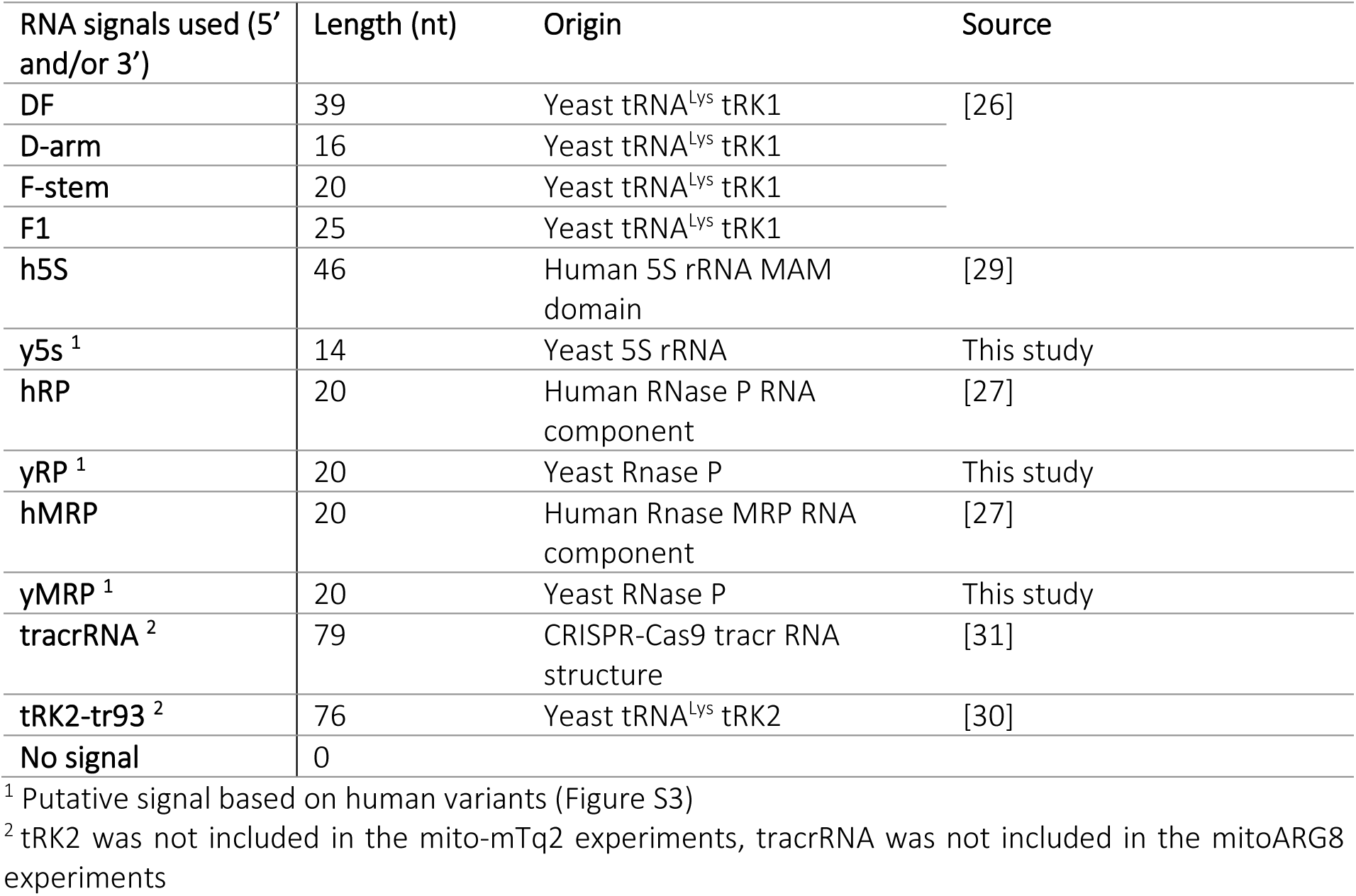
RNA targeting signals used in this study.

| RNA signals used (5' and/or 3') | Length (nt) | Origin | Source |
| --- | --- | --- | --- |
| DF | 39 | Yeast tRNA<sup>Lys</sup> tRK1 | [26] |
| D-arm | 16 | Yeast tRNA<sup>Lys</sup> tRK1 |  |
| F-stem | 20 | Yeast tRNA<sup>Lys</sup> tRK1 |  |
| F1 | 25 | Yeast tRNA<sup>Lys</sup> tRK1 |  |
| h5S | 46 | Human 5S rRNA MAM domain | [29] |
| y5s <sup>1</sup> | 14 | Yeast 5S rRNA | This study |
| hRP | 20 | Human RNase P RNA component | [27] |
| yRP <sup>1</sup> | 20 | Yeast Rnase P | This study |
| hMRP | 20 | Human Rnase MRP RNA component | [27] |
| yMRP <sup>1</sup> | 20 | Yeast RNase P | This study |
| tracrRNA <sup>2</sup> | 79 | CRISPR-Cas9 tracr RNA structure | [31] |
| tRK2-tr93 <sup>2</sup> | 76 | Yeast tRNA<sup>Lys</sup> tRK2 | [30] |
| No signal | 0 |  |  |

<sup>1</sup> Putative signal based on human variants (Figure S3)
<sup>2</sup> tRK2 was not included in the mito-mTq2 experiments, tracrRNA was not included in the mitoARG8 experiments

**Figure 1.**
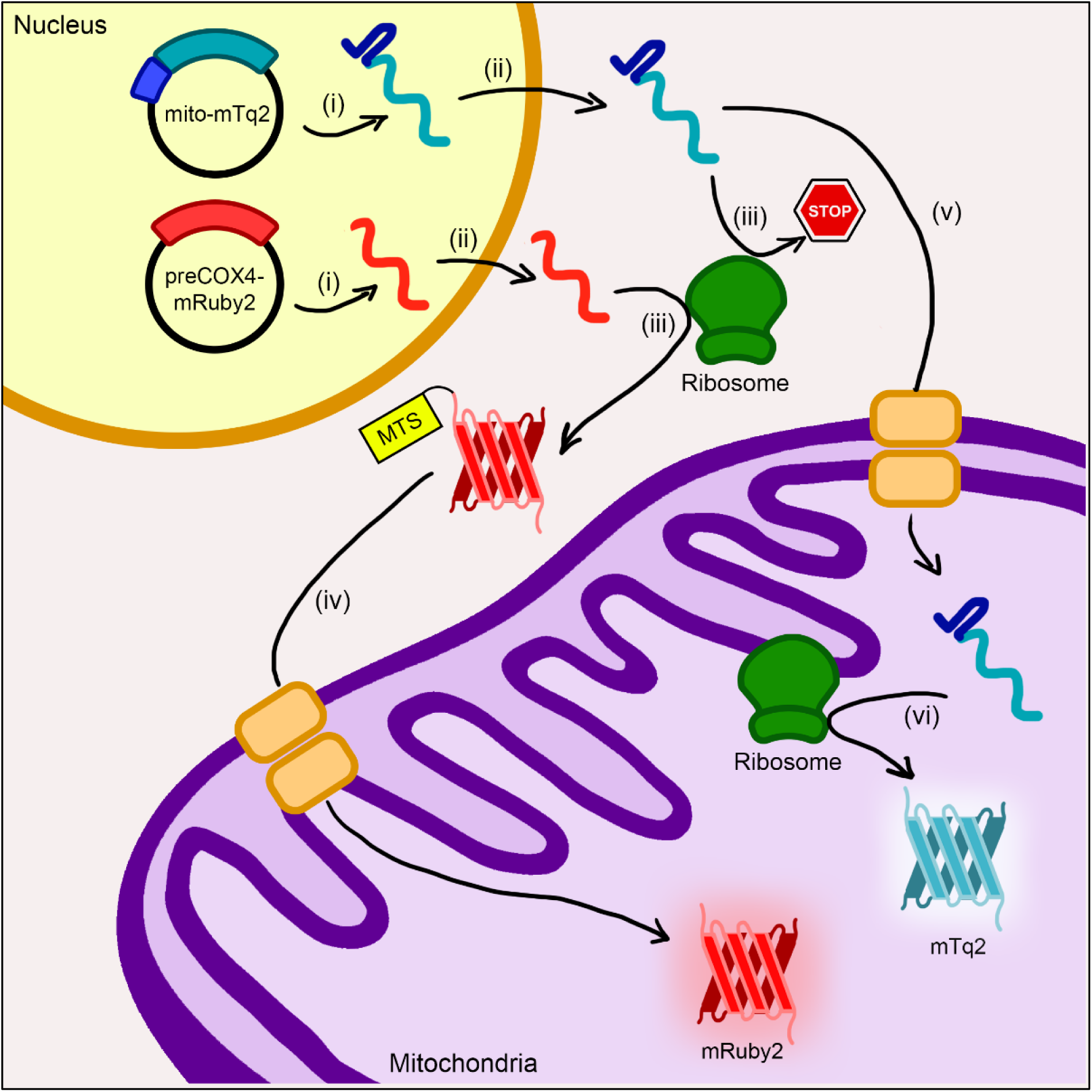
Schematic overview of the screening method to detect RNA import in mitochondria. (i) mitochondrially recoded mTurquoise2 (mito-mTq2) fused to an RNA targeting signal (dark blue) is expressed from a plasmid, as well as mRuby (standard codon usage) fused to mitochondrial targeting signal preCOX4. (ii) Both mRNAs are expressed using Polymerase II promoters and therefore contain a 5’ cap and poly(A)-tail, ensuring export from the nucleus, which are cleaved off by ribozymes attached to the 5’ and 3’ end (not pictured). (iii) mRuby is translated by cytosolic ribosomes, the mito-mTq2 mRNA is mitochondrially recoded and therefore contains internal STOP codons, yielding a truncated protein when expressed in the cytosol. (iv) mRuby2 is fused to a mitochondrial targeting signal (MTS) preCOX4, leading to translocation of the protein upon translation to the mitochondrial matrix, which results in localized red fluorescence in the mitochondria. (v, vi) mito-mTq2 mRNA that is imported into mitochondria can be translated into the full mTq2 protein by the mitochondrial ribosomes, resulting in localized blue fluorescence in the mitochondria. Co-localization of both fluorescent signals indicate successful mitochondrial RNA import.

To benchmark the fluorescence assays, two negative controls, the parental strain CEN.PK113-5D expressing no fluorescent protein and IMC159 only expressing preCOX4-mRuby2, and the positive control IMC173 expressing the mTq2 protein targeted to the mitochondria with a preSU9-MTS (called preSU9-mTq2), but no mRuby2 protein, were included. As expected, the negative controls IMC159 and CEN.PK123-5D did not show any visible fluorescence in mitochondria in the blue channel, while the positive control IMC173 did, and only IMC159 showed red fluorescence, localized to the mitochondria (Figure 2A).

**Figure 2.**
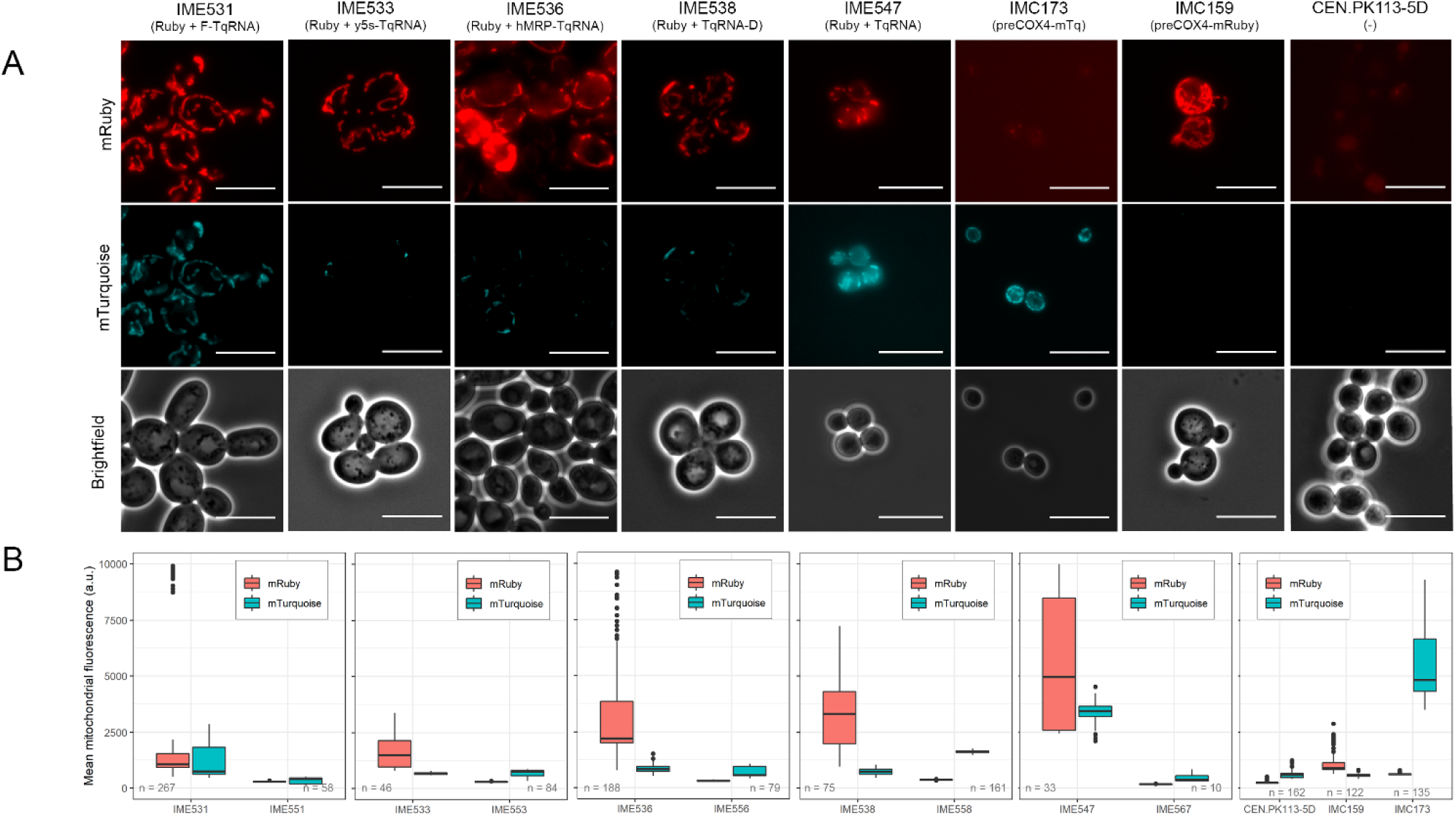
Analysis of colocalization of red (mRuby2) and blue (mTq2) fluorescence in strains expressing preCOX4-mRuby and/or mito-mTq2. Shown is a representative subset of data, the full dataset is shown in the supplementary data (Figure S 5, Figure S 6). A) Fluorescence microscopy images (1000 x) in red (mRuby), blue (mTurquoise) and brightfield images of strains expressing preCOX4-mRuby2 and mito-mTq2 fused to different mitochondrial RNA import signals (IME531-IME547), positive control IMC173, negative control IMC159 and parental strain CEN.PK113-5D. Scale bars represent 10 μm. B) Distribution of the mean mitochondrial fluorescence in the different strains in the mRuby2 (red) and mTurquoise2 (blue) channels. The distribution is presented as boxplots, where the line represents the median value, and the edges of the boxes represent the 25^th^ and 75^th^ percentile. Outliers are shown as dots. Each plot contains mito-mTq2 plus an import signal in the presence (IME531-IME547) or absence (IME551-IME567) of preCox4-mRuby2.

### mTq2 fluorescence appears dependent on the presence of mRuby2

First inspection by fluorescence microscopy of strains expressing mito-mTq2 revealed blue fluorescence specifically localized to mitochondria. This localization was confirmed by the co-localization with nuclear-encoded preCOX4-mRuby2. While these results suggested import and translation of mito-mTq2 mRNA in the mitochondria, a number of confounding observations clouded this conclusion.

First of all, blue fluorescence localized to the mitochondria could only be detected in strains co-expressing both fluorescent proteins, and this blue fluorescence systematically co-localized with red fluorescence (Figure 2A, Figure S5). No blue fluorescence was observed in the absence of preCOX4-mRuby2. For most strains, the mean blue fluorescence intensity was in the range of the negative control strains IMC159 and CEN.PK113-5D that did not express mTq2 (Figure 2B, Figure S6), and only a few strains showed fluorescence above the negative controls (IME531, IME532 and IME547). There is little spectral overlap between mTurquoise2 and mRuby2 (Fig. S2), and the filter set used for excitation and emission of mTurquoise2 do not allow for detection of mRuby2. Accordingly, IMC159, the control strain expressing mRuby2 alone, only showed red and no blue fluorescence in mitochondria. Bleed-through of mRuby2 in the blue channel can therefore be excluded. Consequently, the mRuby2-dependent blue fluorescence is therefore unlikely to be emitted by mRuby2, and probably originates from mTurquoise2. As mTurquoise2 can only be functionally translated in mitochondria, the results suggested that copies of mTurquoise mRNA were localized to and translated in mitochondria.

Heterogeneity in the blue channel was observed not only between strains, but also between cells of the same strain. For instance, IME531 (F-stem) and IME547 (No RNA import signal. Fig. 2) showed blue fluorescence above background at population level, while a fraction of the cell of IME536 (hMRP, Figure 2B), IME537 and IME541 (5’-yMRP and 3’-y5S, resp. Figure S6) showed substantial fluorescence. Remarkably, IME547 was expressing mito-mTq2 without any import signal. The faint and heterogeneous expression of mito-mTq2, together with the lack of requirement of RNA import signal, suggested that the mito-mTq2 mRNA import might be a stochastic process. The connection between mRuby2 and mTurquoise2 fluorescence was elusive. The two fluorescent proteins were expressed with very different expression systems, with preCOX4-mRuby2 integrated in the genome and mito-mTq2 expressed from a multicopy plasmid with strong promoter. Genetic interactions between the two genes is therefore not expected. Alternative explanations like the ability of mRuby2 itself to enhances mTq2 mRNA import to the mitochondria, or the impact of strong overexpression of a mitochondrially-targeted protein on mRNA import are also unlikely, but cannot be entirely excluded.

### Alternative methods to probe mRNA localization in the mitochondria

Fluorescence imaging suggested the presence of functional mTurquoise2 in mitochondria, and thereby the presence of mTq mRNA in mitochondria, however the low fluorescence intensity in the blue channel and its dependence on co-expression with mRuby2 made the fluorescence microscopy results hard to interpret. To find more tangible evidence of the presence of mitochondrial mTq2, RNA FISH and western blotting were performed, the former to detect the presence of mTq2 mRNA and the latter to probe the presence of the mTurquoise2 protein in mitochondria.

For RNA-FISH, fluorescent probes targeting mito-mTq2 mRNA were designed, and labelled cell samples were analyzed by fluorescence microscopy. DAPI was used as marker of cellular DNA content, and *COX3* mRNA, which is transcribed from the mitochondrial genome, was targeted by fluorescent probes and used as indicator of mitochondria-localized mRNA (Figure 3A, Figure S7). The mRuby2 protein was omitted to prevent spectral overlap with the *COX3*-probe. As expected, the control strain that did not express mito-mTq2 did not show fluorescence in the mTq2 probe channel. All other strains expressing mito-mTq2, with or without mitochondrial targeting signal, displayed fluorescence in the mTq2 channel. This fluorescence was substantially more intense than the fluorescence observed with *COX3*-targetting probes, and displayed both a diffuse background and intense foci (Figure 3A, Figure S7). As Mito-mTq2 was expressed from a multicopy plasmid and with a strong promoter in the nucleus, a high number of Mito-mTq2 mRNA copies was expected in the cells, in line with the strong fluorescence intensity observed in the mTq2 probe channel. Notably, the mito-mTq2 probes foci only partially overlapped with the *COX3* foci. However, due to the scattered fluorescence of mito-mTq2 probes through the entirety of the cell and partial co-localization with *COX3* probes fluorescence, the subcellular localization of mTq2 mRNA to mitochondria could not be concluded (Figure 3A, Figure S7).

**Figure 3.**
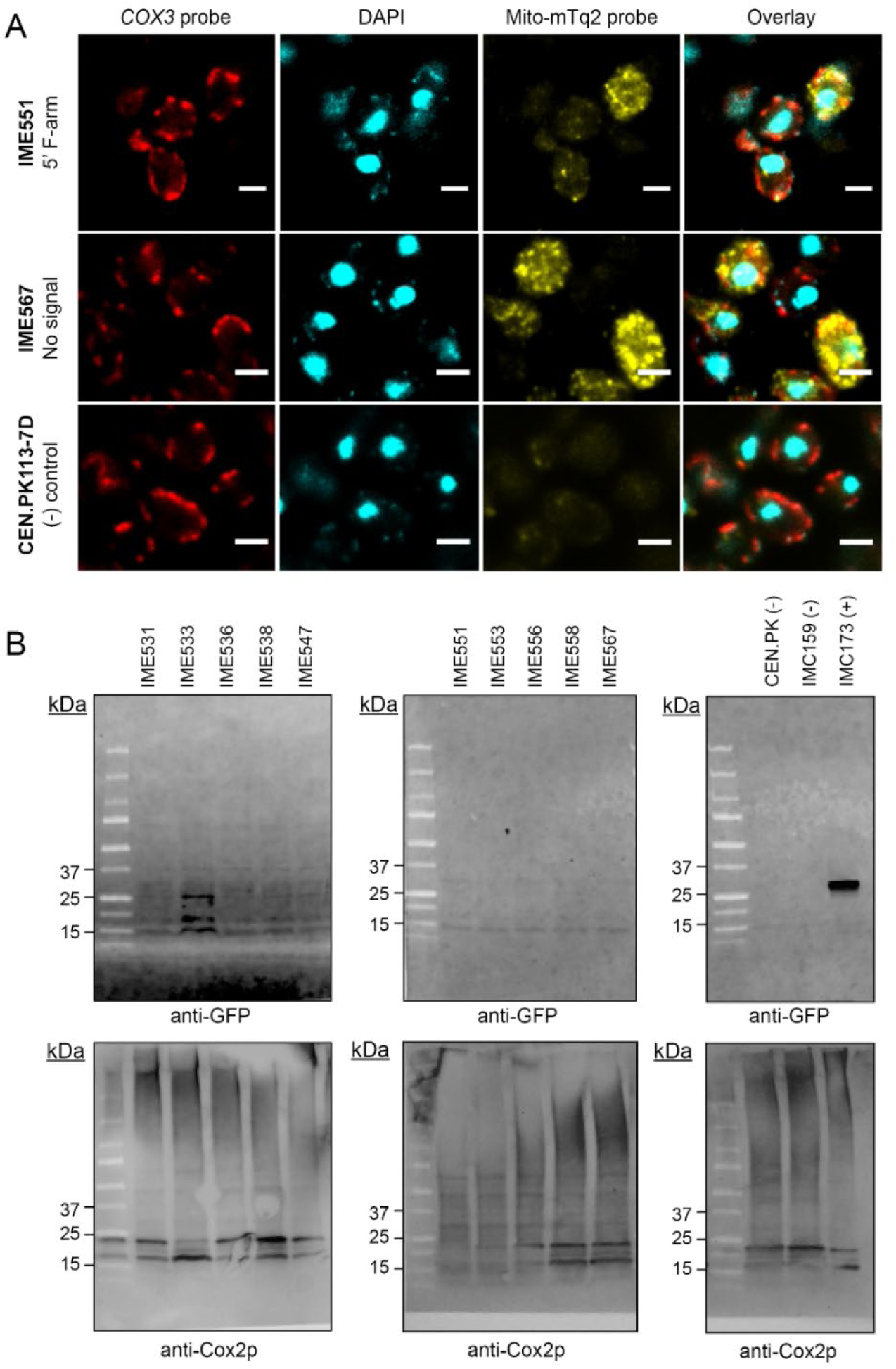
Analysis of transcription and translation of mito-mTq2 mRNA by mRNA-FISH and Western blot. A) Representative figures of mRNA-FISH analysis to determine mito-mTq2 localization, the full dataset is shown in Figure S7. Mitochondrially transcribed *COX3* mRNA was targeted with a specific probe labeled with CAL Fluor Red 635, while mito-mTq2 mRNA was targeted with a specific probe labeled with Quasar-570. DNA was stained with a fluorescent DNA-specific DAPI stain. Strain IME551 expressed mito-mTq2 mRNA fused to the 5’ F-arm RNA signal, IME567 expressed mito-mTq2 with no import signal and negative control CEN.PK113-7D did not express mito-mTq2 mRNA. Acquisition in the *COX3* probe channel (red), mito-mTq2 probe channel (yellow) and the DAPI channel (blue) were respectively done with exposure times of 1000, 600 and 100 seconds, at a magnification of 1000x. Every image was derived from a single slice in a Z-stack after iterative deconvolution. The white scale bar indicates 5 µm. B) Western blot analysis of mTurquoise2 expression in strains expressing mito-mTq2 mRNA in the presence (IME531-547) and absence (IME551-567) of the of the mitochondrially localized mRuby2 protein. Negative controls (-) include CEN.PK113-7D (no fluorescent protein) and IMC159 (preCOX4-mRuby2), the positive control IMC173 (+) expressed preSU9-mTq2. Mito-mTq2 mRNA fused to different RNA import signals were tested - IME531/551: 5’ F-stem, IME533/553: 5’ y5S, IME535/556: 5’ hMRP, IME538/558: 3’ D-loop, IME547/567: no signal. The top row shows blots treated with a fluorescent anti-GFP antibody. The lower row shows the same strains treated with an anti-Cox3p imaged using chemiluminescence. Cox3p is a mitochondrial protein used as loading control. Approximate protein sizes of the observed bands in kDa are indicated on the left. Each lane was loaded with 25 μg protein, except for IMC173 (10 μg protein). Expected protein sizes are 26.9 kDa for mTq2 and 30 kDa for Cox3p, but earlier experiments show that this protein runs lower and suffers from unspecific binding in isolated mitochondria [49]. Uncropped blots are shown in SI figures (Figure S9)

The next detection method took advantage of the difference between mitochondrial and cytosolic translation mechanisms. As indicated above, the mito-mTq2 gene was designed to produce the full 26,9 kDa mTurquoise2 if translated in mitochondria, and truncated translation products of 6 or 18.5 kDa if translated in the cytosol (Figure S4). It is unknown to what extent the cytosolic translation occurs, as the mRNA cap and poly(A)-tail, required for cytosolic translation, are cleaved off by ribozymes upon nuclear export (Figure 1). Western blotting was performed to detect mTurquoise2 presence and to quantify its size. To reduce contamination by cytosolic proteins, Western blots were performed on isolated mitochondria. As GFP has 95 % amino acid sequence similarity with mTq2, but low similarity to mRuby2, antibodies with affinity to GFP were used to specifically detect mTq2 (Figure S8). Accordingly, the anti-GFP probe gave no signal in the negative control strains CEN.PK113-7D (no fluorescent protein) and IMC159 (expressing mRuby2 only, Figure 3B, Figure S9). The positive control IMC173 expressing only preSU9-mTq2 translated in the cytosol and imported into mitochondria did show the expected band of 26,9 KDa (Figure 3B). A subset of strains that expressed mito-mTq2 mRNA only (IME551 – IME567), or mito-mTq2 mRNA in combination with the mRuby2 protein (IME531 – IME547) were tested by Western blotting. Antibodies targeting the mitochondrial protein Cox2p were included as a loading control. The strains expressing both mito-mTq2 and preCOX4-mRuby2 (IME531-IME547, Figure 3B) displayed three faint bands. One of these bands was visible at a height corresponding to the mitochondrially translated mito-mTq2 (ca. 27 kDa). A second band was visible at a height just over 15 kDa. This band could correspond to the 18.5 kDa cytosolic translation product of mito-mTq2 that may have been carried over during isolation of mitochondria, indicating that limited cytosolic translation may still occur. These bands were particularly intense for strain IME533 harboring the 5’ y5S signal fused to mito-mTq2(Figure 3). These bands were not visible in the strains not expressing preSU9-mRuby2 (IME551-IME567, Figure 3). In all tested mito-mTq2 expressing strains, a faint band of just below 15 kDa was visible, both in presence and absence of preSU9-mRuby2 (IME531-IME567). This size does not correspond to any of the predicting peptide resulting from mito-mTq2 translation. It is unclear what this originates from and could be the result of incomplete translation, protein degradation or unspecific interaction with another protein. As observed with fluorescence imaging of the mTq2 protein, Western blotting suggested the presence of mTq2 in the mitochondria with the expected size, and the dependence of mTq2 localization into mitochondria on co-expression with mRuby2.

### Exploring Adaptive Laboratory evolution (ALE) as tool to enhance mRNA import in mitochondria

The three different analytical techniques implemented to assess whether mRNA were imported in mitochondria could not unambiguously confirm mRNA import. The technical difficulty to assess mRNA import in mitochondria with high confidence might simply be caused by the lack of import, but it could also be explained by the low number of mTurquoise2 mRNAs and proteins copies present in mitochondria. In a last attempt to identify mitochondrial import of mRNAs to mitochondria, we decided not to rely on fluorescent detection but to rather use a growth-based screen. This decision was motivated by two factors, the first is the simplicity of screening for mRNA import by monitoring growth, the second it the possibility to use Adaptive Laboratory Evolution (ALE) to improve weak mRNA import. While never used so far for this specific purpose, ALE has already been successfully used to improve complex cellular features (e.g. temperature or chemical tolerance [74]). Starting from the assumption that some of the constructed strains might be able to import and translate a few mRNA copies in mitochondria, a well-designed ALE strategy could drive the selection of mutants with enhanced mitochondrial mRNA import of strategically chosen protein. Acetylornithine aminotransferase, a mitochondrial protein essential for arginine biosynthesis encoded by *ARG8*, was chosen as mRNA cargo. Arg8 is natively translated in the cytosol and imported into the mitochondria, but previous research has shown that an *ARG8* deletion can be complemented by integration of *ARG8* in the mitochondrial genome where it can be transcribed and translated into a functional protein [75]. The novel strategy developed in the present study is to complement an *ARG8* deletion by promoting import of *ARG8* mRNA in the mitochondria where it can be translated into acetylornithine aminotransferase. While *arg8*Δ strains are auxotrophic for arginine, mutants of *arg8*Δ able to import *ARG8* mRNA into mitochondria would be endowed with the ability to grow without arginine supplementation and would therefore outcompete their *arg8*Δ unevolved parent in arginine-limited environment. In other words, a growth-based competition assay would select for *arg8*Δ-derived mutants proficient for mRNA mitochondrial import.

This assay was performed in an arginine-limited chemostat supplied with both ammonium sulfate and arginine. *arg8*Δ strains t are unable to use ammonium for the synthesis of arginine and therefore fully rely on the arginine supplied in the medium for growth. Mutants able to import and translate *ARG8* mRNA in the mitochondria would be able to use ammonium as nitrogen source for the synthesis of arginine and would therefore have a selective advantage and gradually take over the culture. An increase in biomass yield would be an indicator arginine biosynthesis by evolved strains and therefore of *ARG8* mRNA mitochondrial import. Ethanol was used as sole carbon source to increase mitochondrial mass, retain a mitochondrial membrane potential and prevent loss of mitochondrial DNA [76, 77]. –– The chemostat cultures were started by a mixture of *arg8*Δ*car2*Δ mutants equipped with *ARG8* mRNA variants targeted to mitochondria. *CAR2* encoding L-ornithine transaminase was deleted in addition to *ARG8* to prevent the occurrence of arginine prototrophic revertants during long-term culture [46]. The parental strain IMX2796 (*arg8*Δ*car2*Δ) also constitutively expressed preCox4-mRuby2 from the nuclear DNA, as mRuby2 allowed to monitor mitochondrial morphology and potential contaminations during laboratory evolution. Twelve multicopy plasmids carrying the mito-*ARG8* gene without or with one of eleven different 5’ RNA import signals were transformed to the *arg8*Δ*car2*Δ strain IMX2796, resulting in strains IME759-770. None of these strains were readily able to grow in the absence of arginine (Figure S10), indicating that RNA import or translation of the mito-Arg8 mRNA did not occur or were not efficient enough to sustain arginine requirements for growth.

Two bioreactors were inoculated with a mixture of the twelve strains IME759-770 present in equal cell number, leading to two parallel evolution lines named A and B (Figure 4A, Figure S11). The cultures were maintained at a dilution rate (equivalent to the specific growth rate) of 0.1 h^-1^. The theoretical maximum biomass concentration calculated solely based on arginine consumption (60 mg L^-1^) was ca. 2 g L^-1^ [78]. Full consumption of both arginine and ammonium would result in a maximum biomass concentration of 12 g L^-1^, with complete consumption of the supplied ethanol (435 mM). The batch phase preceding the chemostat culture reached a biomass concentration of 3 g L^-1^, which was higher than expected and likely due to residual arginine carried-over from the preculture (performed with 450 mg L^-1^ arginine). During the first 600 hours (*ca.* 86 generations), the chemostat cultures displayed rather steady biomass and residual ethanol concentrations around 1.42 ± 0.14 g L^-1^ and 351 ± 7.2 mM, respectively (Figure 4A, Figure S11). Carbon dioxide and oxygen concentrations however oscillated during this phase. Such oscillations, distinct from fast metabolic oscillations [79], have been previously reported for chemostat cultures of *S. cerevisiae* and might reflect cellular stress due to the limited arginine supply and periodic shifts in intracellular arginine pools [80–84]. After approximately 750 hours (*ca*. 107 generations), the oscillations disappeared while biomass concentration increased to 2.57 ± 0.09 g L^-1^. Concomitantly, carbon dioxide concentration in the off-gas increased by 40 – 50 % and the residual ethanol concentration decreased by 10 % (Figure 5A,B). While the increase in biomass concentration was not as high as expected if all supplied ammonium was consumed, a 60 % increase was observed (Figure 4A, Figure S11).

**Figure 4.**
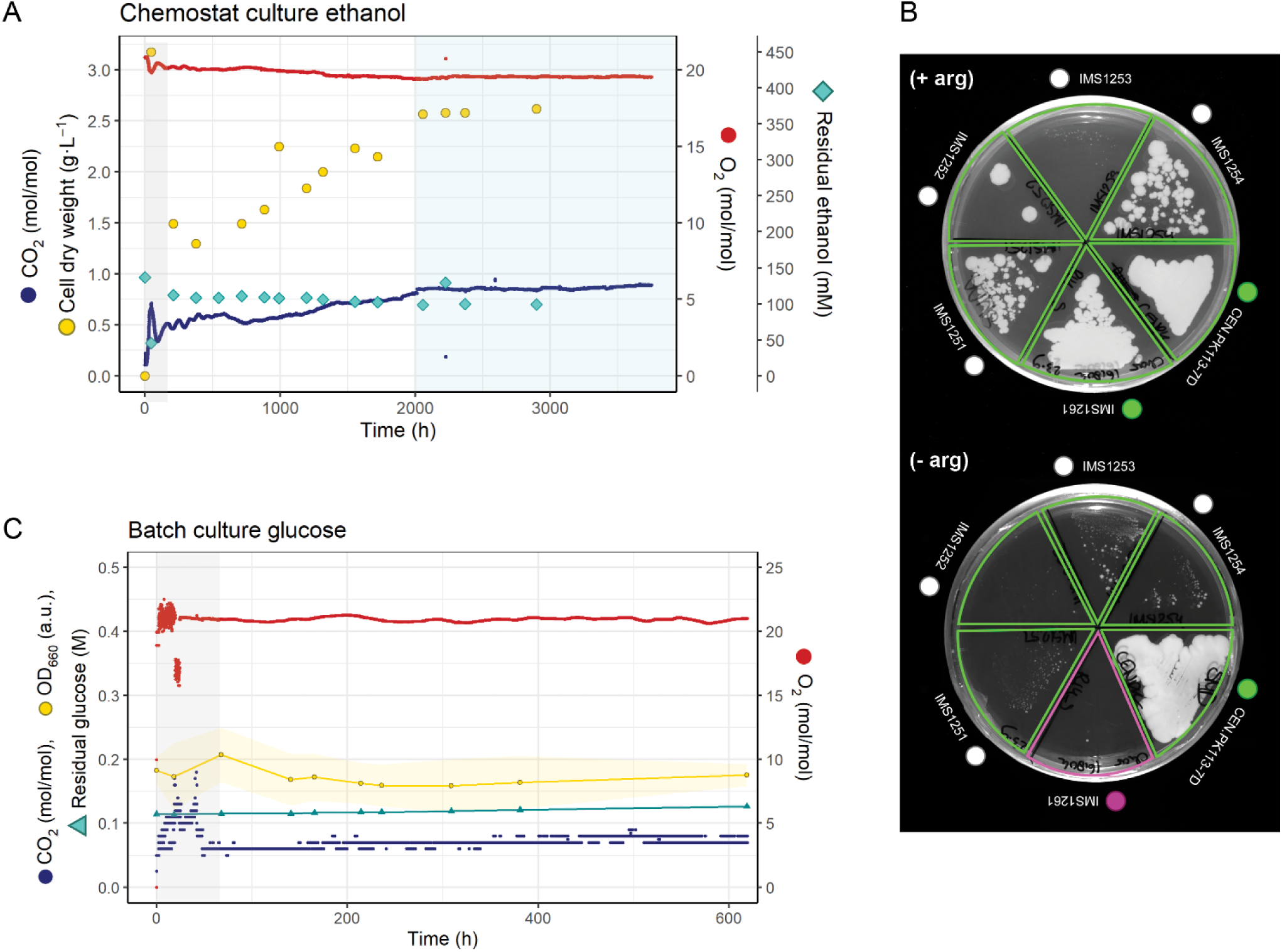
Prolonged chemostat culture of *S. cerevisiae* to evolve towards mito-ARG8 mRNA import in mitochondria and growth characterization of single-colony isolates. A) Profile of one of two replicate bioreactors (Replicate B), inoculated with a mixture of IMC224-IMC235, arginine auxotrophic strains expressing mito-ARG8 mRNA and the PreCOX4-mRuby2 protein. The chemostats were operated at a dilution rate of 0.1 h^-1^, under nitrogen (arginine) limitation and ethanol as source carbon and energy source. The grey area represents the batch phase, the blue area highlights the phase with relatively steady biomass concentration. Profiles of replicate A are shown in Figure S11. B) Growth of four single-colony isolates IMS1251-IMS1254 from bioreactor replicate B on medium with arginine (top)) and without arginine (bottom), along with control strain CEN.PK113-7D (prototroph) and the population of bioreactor B at the end of the batch phase (IMS1261). C) Profiles of the evolved single colony isolate IMS1254 in batch cultivation in a bioreactor in the absence of arginine. Growth in synthetic medium supplied with ammonium as sole nitrogen source and glucose as sole carbon source. CO_2_ and O_2_ concentrations were measured continuously in the off-gas of the bioreactor. The averaged data of two bioreactors is shown, standard deviation is shown as shaded areas. Individual replicates are shown in Figure S12, as well as a culture grown on ethanol as carbon source. Optical density (OD_660_), cell count and the residual glucose concentrations were measured offline. The pre-cultures were grown in shake flasks in the presence of arginine (450 mg L^-1^). After inoculation of the bioreactor with this pre-culture, a first batch was run to deplete the residual arginine carried over from the pre-culture (grey area). At the end of this first batch, the biomass was let to settle by stopping stirring and aeration and the bioreactor was partially emptied and refilled with fresh, arginine-free medium. The white background reflects the complete absence of arginine.

### Characterization of the yeast strains evolved for arginine prototrophy

The increase in biomass concentration observed in chemostat revealed that the yeast population evolved towards more efficient use of the limited supply of arginine. This efficiency might come from the consumption of part of the supplied arginine, but could also result from optimization of ammonium assimilation, for example by enhanced recycling of cellular nitrogen. To test the ability of the evolved strains to synthetize arginine, samples from the chemostat population as well as single colony isolates were plated on minimal, chemically-defined medium with and without arginine and with glucose as carbon source. While cells from evolution line A were not able to grow in the absence of arginine, whole population samples and three out of the four single colony isolates from evolution line B formed colonies in arginine-free medium (Figure 4B). Growth was however extremely slow, with approximately 100 colonies formed per 10^7^ cells plated after 3 weeks. While these results suggested that a fraction of the cell population in evolution line B had acquired the ability to synthetize arginine, growth could not be observed in liquid medium. Even after 40 days, cultures in bioreactors using glucose or ethanol as carbon source did not support growth of the evolved strains in the absence of arginine (Figure 4C, Figure S12). These contradictory results were inconclusive regarding the prototrophy of the evolved strains for arginine, and thereby on the potential mitochondrial localization of Arg8p.

To obtain more solid data regarding the presence of Arg8p in mitochondria of the evolved strains, proteomics analysis was performed on isolated mitochondria of the starting population IMS1261 and the evolved strains IMS1251, IMS2152 and IMS1254. IMS1253 was excluded from this analysis as it grew poorly even in the presence of arginine. The strains were grown in minimal synthetic medium supplemented with arginine and with ethanol as carbon source. None of the peptides of the Arg8p protein were detected in the proteome of the isolated mitochondria of evolved strains (SI file 2), indicating that the mito-Arg8 mRNA was most probably not imported or translated in mitochondria.

The observed increased in biomass concentration in chemostat could therefore be explained by a more efficient use of the supplied arginine by the evolved strains. Accordingly, comparing the proteome of evolved strains IMS1251, IMS2152 and IMS1254 to the non-evolved, starting culture IMS1261 revealed among the 927 proteins detected in all samples a decreased abundance for proteins involved in respiration, nitrogen metabolism and protein synthesis in the evolved strains (Figures S14-S16).

Additionally, analysis of whole genome sequence data of the evolved strains from evolution line B and from the evolved population sampled at the end of the evolution lines A and B (IMS1248 and IMS1249, respectively) also identified modifications in nitrogen metabolism. A set of 38 non-synonymous mutations were found in the genome of one or more of the evolved isolates (Tables S3 and S4, SI file 1). As culture samples from evolution line A did not show any ability to grow in the absence of arginine, mutations also found in this evolution line were discarded, resulting in a set of five genes (*UBR2*, *ASG1*, *ECM21*,*TIF3* and YOR389w) that might form the molecular basis of the improved growth in arginine-limited chemostats. *UBR2* displayed the same mutation in all four evolved strains, *ASG1* and *ECM21* were mutated in a single evolved strain, *TIF3* and YOR389w harbored the same mutation in two evolved strains (Table S3). Four out of these five proteins are involved in protein turn-over, either via synthesis (*TIF3* encodes the translation initiation factor eIF-4B and *ASG1* is a transcriptional regulator) or degradation through the proteasome (*ECM21* and *UBR2*).

The evolution lines were started with a mixture of strains, each with a different putative RNA import signal attached to the mito-Arg8 mRNA. All four evolved isolates contained the same plasmid, encoding the 72-base pair tr93-tRK2 import signal, a modified version of the full tRNA^Lys^(CUU) tRK2 that was described to be imported in mitochondria [85]. The fact that many mutations were shared between the four evolved isolates suggests that they share a common ancestor, rather than indicating that tr93-tRK2 is an optimal RNA import signal. No mutations that could enhance mitochondrial mRNA import or translation were detected in the plasmid sequences.

Altogether these results suggested that the evolved strains became more efficient for growth under low arginine supply, potentially by altering protein turn-over. The sequencing data did not provide information on alternative routes for arginine synthesis, nor did they lead to the identification of potential candidates for improvement of RNA import to mitochondria.

## Discussion

To date, mitochondrial import of RNA remains a controversial topic, and has so far been only explored for short RNAs such as tRNAs or gRNAs (100 - 200 nucleotides). gRNA localization is typically demonstrated through depletion of mtDNA by targeting Cas9 and a gRNA to the mitochondrial matrix. Although several mitochondrial CRISPR-studies demonstrated a 25 – 50 % reduction of mtDNA abundance in the presence of a Cas9 protein and a gRNA [31, 86, 87], there is as of yet no direct evidence of gRNA targeting to the mitochondrial matrix. A study by Schmiderer, Yudovich (38) uncovered that a mitochondrially targeted Cas9 base-editor had a 1 % editing efficiency on the mtDNA, regardless of the sequence of the targeted gRNA. This data suggested that Cas9 can have off-target effects on the mtDNA regardless of the presence of the gRNA, which may lead to mutations in mtDNA or trigger maintenance systems such as mitophagy [37, 88, 89], resulting in a reduced number of mtDNA copies. Depletion of mtDNA is therefore not a conclusive readout of gRNA targeting. Other studies investigating (t)RNA import often use a combination of mitochondrial subfractionation and subsequent qPCR or Northern blot to demonstrate mitochondrial RNA localization. However, cytosolic RNA adheres to the mitochondrial surface [10, 38, 90–93], and isolated mitochondria, even in highly purified or mitoplast form, still contain circa 20 - 80 % contamination by cytosolic material [90, 94]. Therefore, detection of mRNA import is challenging with these sensitive methods as they do not explicitly demonstrate RNA import, but only RNase protection of the targeted RNA by association to the mitochondrial membrane or mitochondrial proteins [37].

With these challenges in mind, different methods are direly needed to reproducibly demonstrate RNA import, without relying on Cas9 activity, and to prevent potential false-positives caused by cytosolic contamination. In this study, this was attempted by targeting a mitochondrially recoded mRNA to the mitochondrial matrix. Mitochondrially recoded genes can only be translated in the mitochondria, offering a powerful screening method of mRNA import to mitochondria, and has been applied once before to demonstrate mitochondrial mRNA import [28, 95]. However, the present study demonstrated that, even though mitochondrial translation is highly specific to the mitochondrial matrix, this method is still sensitive to false positive outcomes. When targeting an mRNA encoding a mitochondrially encoded fluorescent protein, import of mRNA in the mitochondria was detected in the form of mitochondrially localized fluorescence. However, the presence or absence of an RNA import signal did not have a measurable effect on mRNA import in the mitochondria. Random import of RNA regardless of its sequence is highly unlikely, as it would result in uncontrolled accumulation of any RNA species in the mitochondria. Additionally, mRNA import appeared to be dependent on the co-import of the mRuby2 protein. This fluorescence could in theory not originate from the co-imported mRuby2 protein, because of the lack of spectral overlap between mRuby and mTurquoise2, and a strain solely expressing mRuby2 did not show blue fluorescence localized in the mitochondria. Even though the TIM and TOM protein import complexes are likely involved in RNA import to mitochondria [25], high nuclear expression and mitochondrial import of an additional protein should not affect RNA import. Approximately 1,000 proteins are natively imported into mitochondria, and are sorted either in the mitochondrial membrane or in the matrix based on their localization signal [2, 96], causing intensive protein trafficking across the mitochondrial membrane in active cells. Even though mitochondrial co-import of RNA and specific proteins has been described in multiple occasions in yeast and mammalian cell lines, these mechanisms are often highly disputed [10, 27, 28, 38, 97–99]. It is therefore unlikely that expressing and importing yet another protein affects RNA import. The mechanism leading to blue fluorescent signal in the mitochondria expressing mito-mTq2 therefore remains elusive. Considering the low likelihood of random RNA import or mRuby2-assisted import, it is probably an experimental artifact, possibly caused by auto-fluorescence of mitochondrial components such as mitochondrial NADH, which has similar spectral properties as mTurquoise2 [100]. These results stress the need for extremely sensitive and specific methods to detect mitochondrial localization of RNAs.

To circumvent the occurrence of false positives connected to fluorescent detection, our second approach relied on enzyme activity-based growth screening. However, high expression of mitochondrially targeted *ARG8* mRNA did not restore the arginine prototrophy of an Δ*arg8* mutant (Figure S 10). This result suggested either that Arg8 mRNA was not readily imported into mitochondria or that Arg8p expression, if present, was too low to sustain growth requirements. The latter hypothesis served as basis for an adaptive laboratory evolution approach. The ALE experiment was carefully designed with a strong selection pressure favoring mutants with enhanced mitochondrial import of mito-*ARG8* mRNA. However, the mito-Arg8 protein was not detected in the proteome of the evolved strains indicating mito-*ARG8* mRNA was not imported or not translated. In line with this result, whole genome sequencing and mitochondrial proteome analysis of the evolved strains did not reveal the occurrence of mechanisms enhancing RNA import to mitochondria. Under the strong selection pressure applied in arginine-limited chemostat, *S. cerevisiae* evolved towards a more efficient utilization of arginine, and did not accumulate mutations enabling mitochondrial RNA import (Figures S14-S16, Tables S3 and S4). The absence of mito-Arg8 mRNA import in mitochondria despite the strong selection pressure for arginine synthesis reveals that RNA import is particularly difficult to evolve for. The lack of RNA import may be explained by different factors such as the length of the RNA species used in this work (1272 and1344 nt) while RNA import is typically described for short RNA molecules (< 200 nt), the structure or charge of the transported RNA or simply the fact that the signal sequences used cannot lead to RNA import. Even if not successful in the present study, the designed ALE strategy is a very valuable tool in efforts to engineer mRNA import into mitochondria. As arginine limitation results in cellular optimization to arginine-poor conditions during prolonged chemostat cultivation, it is advisable to perform a control chemostat with an arginine auxotrophic strain devoid of mitochondrial *ARG8* mRNA import. This control will enable to disentangle yeast native responses to limited arginine supply from effective import of mito-*ARG8* mRNA.

Despite the implementation of innovative approaches based on mitochondria-specific translation and ALE, import of mRNA-sized RNAs was not achieved in the present study. The pitfalls of standard methods to demonstrate RNA import of non-native mRNA into mitochondria, encountered in this study as well as other studies in RNA import, strongly advocate to further develop innovative and more specific approaches to detect RNA import. The import of a long mRNA (up to 1272 nucleotides for *ARG8*) may be too ambitious, but a similar approach might be successful with shorter RNAs. Designing an ALE experiment with selection pressure for the import of a short RNA in mitochondria is challenging, but not impossible. For instance, selecting for complementation of mitochondrially-encoded tRNA, upon removal of the native tRNA encoded by the mtDNA, might offer a powerful ALE strategy.

## Author contributions

C.C.K., K.K., and P.D-L. conceptualized the study and provided scientific input. C.C.K., K.K., M.L. and E.d.H. designed and performed experiments, M.d.R. and M.P. performed proteomic analysis and processed proteome data.

## Acknowledgements

The authors thank Dr. M.M.M. Bisschops for experimental support on the mRNA-FISH and Western blot experiments. This work was funded by the ‘BaSyC—Building a Synthetic Cell’ Gravitation grant (NWO, grant number 024.003.019) and by an Aspasia grant awarded to Pascale Daran-Lapujade by the Dutch Research Council (NWO, grant number 015.014.007).

## List of supplementary information

*Koster_2025_Supplementary-information.pdf*

### Sequences

Putative RNA import sequences used in thi*s study*

Sequences of recoded mito-mRNA

### Supplementary tables

**Table S1.** Plasmids used in this study.

**Table S2.** Primers used for plasmid and strain construction in this study

**Table S3.** All non-silent mutations observed in IMS1248, IMS1249 and IMS1251-1254

**Table S4.** Non-synonymous mutations found in IMS1248, IMS1249 and IMS1251 - IMS1254.

### Supplementary figures

**Figure S1.** Construction strategy of mito-mTq2 mRNA and mito-Arg8 mRNA with different RNA import signals.

**Figure S2.** Spectral overview of fluorophores and filters used for *in vivo* co-localization assays.

**Figure S3.** Structural comparison between human and *S. cerevisiae* putative RNA import signals.

**Figure S4.** Translation frames of mito-mTq2 mRNA.

**Figure S5.** Fluorescence microscopy images of strains expressing mito-mTq2 fused to different mitochondrial RNA import signals.

**Figure S7.** FISH analysis of strains expressing mito-mTq2 mRNA with different RNA signals and negative control CEN.PK113-7D, omitting mito-mTq2 mRNA.

**Figure S8.** Western blot of fluorescent proteins using anti-GFP antibody.

**Figure S9** Western blot analysis of mTurquoise2 expression in strains expressing mito-mTq2 mRNA in with the presence or absence of the preCox4-mRuby2 protein.

**Figure S10** Growth of strains with mito-ARG8 mRNA targeted to mitochondria, on synthetic medium without arginine in microtiterplates.

**Figure S11.** Prolonged chemostat culture of *S. cerevisiae* to evolve towards mito-ARG8 mRNA import in mitochondria

**Figure S12.** Physiological characterization of the evolved single colony isolate IMS1254 in batch cultivation in a bioreactor in the absence of arginine.

**Figure S13.** Fraction of mitochondrial protein detected in each proteome.

**Figure S14.** Global proteome changes between the isolated mitochondria of evolved strains IMS1251, IMS1252 and IMS1254.

**Figure S15.** Proteome response of mitochondrial proteins detected in IMS1251, IMS1252 and IMS1254 compared to unevolved parental population IMS1261.

**Figure S16.** GO-term analysis of 78 proteins with decreased abundance in both strains IMS1251 and IMS1254 as compared to the unevolved parental population IMS1261.

### Supplementary files

**SI File 1** (*SI_1_WGSdata.xlsx)* – Whole genome sequencing data analysis, containing all annotated mutations found in the evolved strains

**SI File 2** (*SI_2_Proteomics_GOterm.xlsx*) – Abundances and annotated proteins found by proteomics analysis of evolved strains, including GO term analysis.

