## Supplementary information for "Exploration of mRNA-sized RNA import into *Saccharomyces cerevisiae* mitochondria by a combined synthetic biology and adaptive laboratory evolution approach"

#### Sequences

*Putative RNA import sequences used in thi*s study

DF-stem (retrieved from [1])
1 gcgcaatcgg tagcgcttcg agccccctac agggctcca

D-arm (retrieved from [1])
1 gcgcaatcgg tagcgc

F-arm (retrieved from [1])
1 gagcccccta cagggctctt

F1 (retrieved from [1])
1 ggttcgagcc ccctacaggg ctcca

tRK2-tr93 (retrieved from [1])
1  gccttgttag ctcagttggt agagcgttcg gcttttaacc gaaatgtcag ggttcgagc
61 cccctatgag gct

Human 5S rRNA (retrieved from [2])
1 ggcctggtta gtacttggat gggggaccgc caaggaatac cgggtg

Yeast 5S rRNA (Figure S 3)
1 gaccgagtag tgtagtgggt gaccatacgc gaaactcagg tg

Human ribonuclease P stem (retrieved from [3])
1 tctccctgag cttcagggag

Yeast ribonuclease P stem (Figure S 3)
1 actggggaac cagt

Human mitochondrial RNA processing ribonuclease stem (retrieved from [3])
1 agaagcgtat cccgctgagc

Yeast mitochondrial RNA processing ribonuclease stem (Figure S 3)
1 gtacctctat tgcagggtac

tracrRNA (from pMEL13 [4])
1   gttttagagc tagaaatagc aagttaaaat aaggctagtc cgttatcaac ttgaaaaagt
61  ggcaccgagt cggtggtgc

*Sequences of recoded mito-mRNA*

#### Mito-mTq2

*The sequence for mTq2 was retrieved from [5], and used for codon-optimization for yeast mitochondrial translation.*

```
1   atggtatcaa aaggtgaaga attatttaca ggtgtagtac ctattttagt agaattagat
61  ggtgatgtaa atggtcataa attttcagta tcaggtgaag gtgaaggtga tgctacatat
121 ggtaaattaa cattaaaatt tatttgtaca acaggtaaat tacctgtacc ttgacctaca
181 ttagtaacaa cattatcatg aggtgtacaa tgttttgcta gatatcctga tcatatgaaa
241 caacatgatt tttttaaatc agctatgcct gaaggttatg tacaagaaag aacaattttt
301 tttaaagatg atggtaatta taaaacaaga gctgaagtaa aatttgaagg tgatacatta
361 gtaaatagaa ttgaattaaa aggtattgat tttaaagaag atggtaatat tttaggtcat
421 aaattagaat ataattattt ttcagataat gtatatatta cagctgataa acaaaaaaat
481 ggtattaaag ctaattttaa aattagacat aatattgaag atggtggtgt acaattagct
541 gatcattatc aacaaaatac acctattggt gatggtcctg tattattacc tgataatcat
601 tatttatcaa cacaatcaaa attatcaaaa gatcctaatg aaaaaagaga tcatatggta
661 ttattagaat ttgtaacagc tgctggtatt acattaggta tggatgaatt atataaataa
```

#### Mito-Arg8 (incl COX2-UTL and dodecamer, underlined)

```
1    agtattaaca tattataaat agacaaaaga gtctaaaggt taagatttat taaaatgttt
61   aaaagatatt tatcatcaac atcatcaaga agatttacat caattttaga agaaaaagct
121  tttcaagtaa caacatattc aagacctgaa gatttatgta ttacaagagg taaaaatgct
181  aaattatatg atgatgtaaa tggtaaagaa tatattgatt ttacagctgg tattgctgta
241  acagctttag gtcatgctaa tcctaaagta gctgaaattt tacatcatca agctaataaa
301  ttagtacatt catcaaattt atattttaca aaagaatgtt tagatttatc agaaaaaatt
361  gtagaaaaaa caaaacaatt tggtggtcaa catgatgctt caagagtatt tttatgtaat
421  tcaggtacag aagctaatga agctgcttta aaatttgcta aaaaacatgg tattatgaaa
481  aatccttcaa aacaaggtat tgtagctttt gaaaattcat ttcatggtag aacaatgggt
541  gctttatcag taacatgaaa ttcaaaatat agaacacctt ttggtgattt agtacctcat
601  gtatcatttt taaatttaaa tgatgaaatg acaaaattac aatcatatat tgaaacaaaa
661  aaagatgaaa ttgctggttt aattgtagaa cctattcaag gtgaaggtgg tgtatttcct
721  gtagaagtag aaaaattaac aggtttaaaa aaaatttgtc aagataatga tgtaattgta
781  attcatgatg aaattcaatg tggtttaggt agatcaggta aattatgagc tcatgcttat
841  ttaccttcag aagctcatcc tgatattttt acatcagcta aagctttagg taatggtttt
901  cctattgctg ctacaattgt aaatgaaaaa gtaaataatg ctttaagagt aggtgatcat
961  ggtacaacat atggtggtaa tcctttagct tgttcagtat caaattatgt attagataca
1021 attgctgatg aagctttttt aaaacaagta tcaaaaaaat cagatatttt acaaaaaaga
1081 ttaagagaaa ttcaagctaa atatcctaat caaattaaaa caattagagg taaaggttta
1141 atgttaggtg ctgaatttgt agaacctcct acagaagtaa ttaaaaaagc tagagaatta
1201 ggtttattaa ttattacagc tggtaaatca acagtaagat ttgtacctgc tttaacaatt
1261 gaagatgaat taattgaaga aggtatggat gcttttgaaa aagctattga agctgtatat
1321 gcttaattat aataatattc ttaa
```

### Supplementary tables

Table S1. Plasmids used in this study. "YTK type #" refers to yeast Toolkit parts as described by Lee, DeLoache (6)

| Name | Relevant characteristics | Source |
|---|---|---|
| | Insert(s) | |
| pUD518 | GFP-dropout, *KanR, ColE1* | [7] |
| pUD1094 | mitochondrial codon optimized mTq2 (mito-mTq2 mRNA), *bla, ColE1* | Synthetic (GeneArt) |
| pUD1202 | *HH-GFPdropout-Cox2UTL-mitoARG8-dodecamer-HDV, bla, ColE1* | Synthetic (GeneArt) |
| pYTK001 | *CamR, ColE1* | [6] |
| pGGkp265 | mito-mTq2 mRNA (YTK type 3), *CamR, ColE1* | This study |
| pGGkp267 | DF-stem import signal (YTK type 3a), *CamR, ColE1* | This study |
| pGGkp268 | D-arm signal (YTK type 3a), *CamR, ColE1* | This study |
| pGGkp269 | F-arm signal (YTK type 3a), *CamR, ColE1* | This study |
| pGGkp270 | h5S rRNA signal (YTK type 3a), *CamR, ColE1* | This study |
| pGGkp271 | y5S rRNA signal (YTK type 3a), *CamR, ColE1* | This study |
| pGGkp272 | hRP-stem signal (YTK type 3a), *CamR, ColE1* | This study |
| pGGkp273 | Putative yRP signal (YTK type 3a), *CamR, ColE1* | This study |
| pGGkp274 | hMRP RP-stem signal (YTK type 3a), *CamR, ColE1* | This study |
| pGGkp275 | Putative yMRP signal (YTK type 3a), *CamR, ColE1* | This study |
| pGGkp280 | Mito-mTq2 mRNA ((YTK type 3b), *CamR, ColE1* | This study |
| pGGkp283 | D-arm signal (YTK type 4a), *CamR, ColE1* | This study |
| pGGkp284 | F-arm signal (YTK type 4a), *CamR, ColE1* | This study |
| pGGkp285 | h5S rRNA signal (YTK type 4a), *CamR, ColE1* | This study |
| pGGkp286 | y5S rRNA signal (YTK type 4a), *CamR, ColE1* | This study |
| pGGkp287 | hRP-stem signal (YTK type 4a), *CamR, ColE1* | This study |
| pGGkp288 | Putative yRP signal (YTK type 4a), *CamR, ColE1* | This study |
| pGGkp289 | hMRP RP-stem signal (YTK type 4a), *CamR, ColE1* | This study |
| pGGkp290 | Putative yMRP signal (YTK type 4a), *CamR, ColE1* | This study |
| pGGkp291 | tracrRNA signal (YTK type 4a), *CamR, ColE1* | This study |
| pUDC191 | *pCCW12-mRuby2-tENO1 URA3, CEN6/ARS4, bla, ColE1* | |
| pUDC192 | *pTEF2-mTurquoise2-tSSA1 URA3, CEN6/ARS4, bla, ColE1* | |
| pUDC193 | *pTEF1-Venus-tTDH1 URA3, CEN6/ARS4, bla, ColE1* | |
| pUDC286 | *pTDH3-preCOX4-mRuby2-tADH1 URA3, CEN6/ARS4, bla, ColE1* | [8] |
| pUDC329 | *pTEF1*-preSU9-ymTq2-*tENO2, URA3, CEN6/ARS4, bla, ColE1* | [9] |
| pUDC365 | *pTEF1-HH-GFPdropout-Cox2UTL-mitoARG8-dodecamer-HDV-tENO2, URA3, CEN6/ARS4, KanR, ColE1* | This study |
| pUDC434 | *pTDH3-preCOX4MTS-ARG8-tADH1, URA3, CEN6/ARS4, bla, ColE1* | [10] |
| pUDE929 | *pPGK1*-HH-*dKanMX*-HDV-*tADH1, HygR, 2µ, bla, ColE1* | This study |
| pUDE969 | *pPGK1*-HH-mito-mTq2-HDV-*tADH1, HygR, 2µ, bla, ColE1* | This study |
| pUDE970 | *pPGK1*-HH-DF-mito-mTq2-HDV-*tADH1, HygR, 2µ, bla, ColE1* | This study |
| pUDE971 | *pPGK1*-HH-D-mito-mTq2-HDV-*tADH1, HygR, 2µ, bla, ColE1* | This study |
| pUDE972 | *pPGK1*-HH-F-mito-mTq2-HDV-*tADH1, HygR, 2µ, bla, ColE1* | This study |
| pUDE973 | *pPGK1*-HH-h5S-mito-mTq2-HDV-*tADH1, HygR, 2µ, bla, ColE1* | This study |
| pUDE974 | *pPGK1*-HH-y5S-mito-mTq2-HDV-*tADH1, HygR, 2µ, bla, ColE1* | This study |
| pUDE975 | *pPGK1*-HH-hRP-mito-mTq2-HDV-*tADH1, HygR, 2µ, bla, ColE1* | This study |
| pUDE976 | *pPGK1*-HH-yRP-mito-mTq2-HDV-*tADH1, HygR, 2µ, bla, ColE1* | This study |
| pUDE977 | *pPGK1*-HH-hMRP-mito-mTq2-HDV-*tADH1, HygR, 2µ, bla, ColE1* | This study |
| pUDE978 | *pPGK1*-HH-yMRP-mito-mTq2-HDV-*tADH1, HygR, 2µ, bla, ColE1* | This study |
| pUDE979 | *pPGK1*-HH-mito-mTq2-D-HDV-*tADH1, HygR, 2µ, bla, ColE1* | This study |
| pUDE980 | *pPGK1*-HH-mito-mTq2-F-HDV-*tADH1, HygR, 2µ, bla, ColE1* | This study |
| pUDE981 | *pPGK1*-HH-mito-mTq2-h5S-HDV-*tADH1, HygR, 2µ, bla, ColE1* | This study |
| pUDE982 | *pPGK1*-HH-mito-mTq2-y5S-HDV-*tADH1, HygR, 2µ, bla, ColE1* | This study |

| | | |
|---|---|---|
| **pUDE983** | *pPGK1*-HH-mito-mTq2-hRP-HDV-*tADH1, HygR, 2µ, bla, ColE1* | This study |
| **pUDE984** | *pPGK1*-HH-mito-mTq2-yRP-HDV-*tADH1, HygR, 2µ, bla, ColE1* | This study |
| **pUDE985** | *pPGK1*-HH-mito-mTq2-hMRP-HDV-*tADH1, HygR, 2µ, bla, ColE1* | This study |
| **pUDE986** | *pPGK1*-HH-mito-mTq2-yMRP-HDV-*tADH1, HygR, 2µ, bla, ColE1* | This study |
| **pUDE987** | *pPGK1*-HH-mito-mTq2-tracr-HDV-*tADH1, HygR, 2µ, bla, ColE1* | This study |
| **pUDE1230** | *pTEF1-HH-D1F1 -Cox2UTL-mitoARG8-dodecamer-HDV-tENO2, URA3, 2µ, KanR, ColE1* | This study |
| **pUDE1231** | *pTEF1-HH-D-Cox2UTL-mitoARG8-dodecamer-HDV-tENO2, URA3, 2µ, KanR, ColE1* | This study |
| **pUDE1232** | *pTEF1-HH-F -Cox2UTL-mitoARG8-dodecamer-HDV-tENO2, URA3, 2µ, KanR, ColE1* | This study |
| **pUDE1233** | *pTEF1-HH-h5S -Cox2UTL-mitoARG8-dodecamer-HDV-tENO2, URA3, 2µ, KanR, ColE1* | This study |
| **pUDE1234** | *pTEF1-HH-y5S -Cox2UTL-mitoARG8-dodecamer-HDV-tENO2, URA3, 2µ, KanR, ColE1* | This study |
| **pUDE1235** | *pTEF1-HH-hRP -Cox2UTL-mitoARG8-dodecamer-HDV-tENO2, URA3, 2µ, KanR, ColE1* | This study |
| **pUDE1236** | *pTEF1-HH-yRP -Cox2UTL-mitoARG8-dodecamer-HDV-tENO2, URA3, 2µ, KanR, ColE1* | This study |
| **pUDE1237** | *pTEF1-HH-hMRP -Cox2UTL-mitoARG8-dodecamer-HDV-tENO2, URA3, 2µ, KanR, ColE1* | This study |
| **pUDE1238** | *pTEF1-HH-yMRP -Cox2UTL-mitoARG8-dodecamer-HDV-tENO2, URA3, 2µ, KanR, ColE1* | This study |
| **pUDE1239** | *pTEF1-HH-F1 -Cox2UTL-mitoARG8-dodecamer-HDV-tENO2, URA3, 2µ, KanR, ColE1* | This study |
| **pUDE1240** | *pTEF1-HH-tr93 -Cox2UTL-mitoARG8-dodecamer-HDV-tENO2, URA3, 2µ, KanR, ColE1* | This study |
| **pUDE1241** | *pTEF1-HH-GFPdropout-Cox2UTL-mitoARG8-dodecamer-HDV-tENO2, URA3, 2µ, KanR, ColE1* | This study |
| **pUDR514** | *2x gRNA-YPRCtau3, KanMX, 2µ, bla, ColE1* | [11] |
| **pMEL13** | *gRNA-CAN1.Y, KanMX, 2µ, bla, ColE1* | [4] |

Table S2. Primers used for plasmid and strain construction in this study

| # ID | PURPOSE* | SEQUENCE |
|---|---|---|
| 1693 | Fw_pUD518 | GCTGCTACTCATCCTAGTCC |
| 3118 | Rv_pUDC365 | CTTGGCAGCAACAGGACTAGGATGAGTAGCAGCACGTTCC |
| 10320 | Fw_pUDC365 | CATGCGCGGATGACACGAAC |
| 10321 | Rv_pUD518 | CGTCGTGAGTTCGTGTCATC |
| 16310 | fw_DF_pt3aYTK | GCATCGTCTCATCGGTCTCATATGGCGCAATCGGTAGCGCTTCGAGCCCCCTACAGGGCTCCATTCTTGAGACCTGAGACGGCAT |
| 16311 | rv_DF_pt3aYTK | ATGCCGTCTCAGGTCTCAAGAATGGAGCCCTGTAGGGGGCTCGAAGCGCTACCGATTGCGCCATATGAGACCGATGAGACGATGC |
| 16312 | fw_D_pt3aYTK | GCATCGTCTCATCGGTCTCATATGGCGCAATCGGTAGCGCTTCTTGAGACCTGAGACGGCAT |
| 16313 | rv_D_pt3aYTK | ATGCCGTCTCAGGTCTCAAGAAGCGCTACCGATTGCGCCATATGAGACCGATGAGACGATGC |
| 16314 | fw_F_pt3aYTK | GCATCGTCTCATCGGTCTCATATGGAGCCCCCTACAGGGCTCTTTTCTTGAGACCTGAGACGGCAT |
| 16315 | rv_F_pt3aYTK | ATGCCGTCTCAGGTCTCAAGAAAAGAGCCCTGTAGGGGGCTCCATATGAGACCGATGAGACGATGC |
| 16316 | fw_h5SrRNA_pt3aYTK | GCATCGTCTCATCGGTCTCATATGGGCCTGGTTAGTACTTGGATGGGGGACCGCCAAGGAATACCGGGTGTTCTTGAGACCTGAGACGGCAT |
| 16317 | rv_h5SrRNA_pt3aYTK | ATGCCGTCTCAGGTCTCAAGAACACCCGGTATTCCTTGGCGGTCCCCCATCCAAGTACTAACCAGGCCCATATGAGACCGATGAGACGATGC |
| 16318 | fw_y5SrRNA_pt3aYTK | GCATCGTCTCATCGGTCTCATATGGACCGAGTAGTGTAGTGGGTGACCATACGCGAAACTCAGGTGTTCTTGAGACCTGAGACGGCAT |
| 16319 | rv_y5SrRNA_pt3aYTK | ATGCCGTCTCAGGTCTCAAGAACACCTGAGTTTCGCGTATGGTCACCCACTACACTACTCGGTCCATATGAGACCGATGAGACGATGC |
| 16320 | fw_hRP_pt3aYTK | GCATCGTCTCATCGGTCTCATATGTCTCCCTGAGCTTCAGGGAGTTCTTGAGACCTGAGACGGCAT |
| 16321 | rv_hRP_pt3aYTK | ATGCCGTCTCAGGTCTCAAGAACTCCCTGAAGCTCAGGGAGACATATGAGACCGATGAGACGATGC |
| 16322 | fw_yRP_pt3aYTK | GCATCGTCTCATCGGTCTCATATGACTGGGGAACCAGTTTCTTGAGACCTGAGACGGCAT |
| 16323 | rv_yRP_pt3aYTK | ATGCCGTCTCAGGTCTCAAGAAACTGGTTCCCCAGTCATATGAGACCGATGAGACGATGC |
| 16324 | fw_hMRP_pt3aYTK | GCATCGTCTCATCGGTCTCATATGAGAAGCGTATCCCGCTGAGCTTCTTGAGACCTGAGACGGCAT |
| 16325 | rv_hMRP_pt3aYTK | ATGCCGTCTCAGGTCTCAAGAAGCTCAGCGGGATACGCTTCTCATATGAGACCGATGAGACGATGC |
| 16326 | fw_yMRP_pt3aYTK | GCATCGTCTCATCGGTCTCATATGGTACCTCTATTGCAGGGTACTTCTTGAGACCTGAGACGGCAT |
| 16327 | rv_yMRP_pt3aYTK | ATGCCGTCTCAGGTCTCAAGAAGTACCCTGCAATAGAGGTACCATATGAGACCGATGAGACGATGC |
| 16336 | fw_D_pt4aYTK | GCATCGTCTCATCGGTCTCAATCCGCGCAATCGGTAGCGCTGGCTGAGACCTGAGACGGCAT |
| 16337 | rv_D_pt4aYTK | ATGCCGTCTCAGGTCTCAGCCAGCGCTACCGATTGCGCGGATTGAGACCGATGAGACGATGC |
| 16338 | fw_F_pt4aYTK | GCATCGTCTCATCGGTCTCAATCCGAGCCCCCTACAGGGCTCTTTGGCTGAGACCTGAGACGGCAT |
| 16339 | rv_F_pt4aYTK | ATGCCGTCTCAGGTCTCAGCCAAAGAGCCCTGTAGGGGGCTCGGATTGAGACCGATGAGACGATGC |
| 16340 | fw_h5SrRNA_pt4aYTK | GCATCGTCTCATCGGTCTCAATCCGGCCTGGTTAGTACTTGGATGGGGGACCGCCAAGGAATACCGGGTGTGGCTGAGACCTGAGACGGCA<br>T |
| 16341 | rv_h5SrRNA_pt4aYTK | ATGCCGTCTCAGGTCTCAGCCACACCCGGTATTCCTTGGCGGTCCCCCATCCAAGTACTAACCAGGCCGGATTGAGACCGATGAGACGATGC |
| 16342 | fw_y5SrRNA_pt4aYTK | GCATCGTCTCATCGGTCTCAATCCGACCGAGTAGTGTAGTGGGTGACCATACGCGAAACTCAGGTGTGGCTGAGACCTGAGACGGCAT |
| 16343 | rv_y5SrRNA_pt4aYTK | ATGCCGTCTCAGGTCTCAGCCACACCTGAGTTTCGCGTATGGTCACCCACTACACTACTCGGTCGGATTGAGACCGATGAGACGATGC |
| 16344 | fw_hRP_pt4aYTK | GCATCGTCTCATCGGTCTCAATCCTCTCCCTGAGCTTCAGGGAGTGGCTGAGACCTGAGACGGCAT |
| 16345 | rv_hRP_pt4aYTK | ATGCCGTCTCAGGTCTCAGCCACTCCCTGAAGCTCAGGGAGAGGATTGAGACCGATGAGACGATGC |

| | | |
|---|---|---|
| 16346 | fw_yRP_pt4aYTK | GCATCGTCTCATCGGTCTCAATCCACTGGGGAACCAGTTGGCTGAGACCTGAGACGGCAT |
| 16347 | rv_yRP_pt4aYTK | ATGCCGTCTCAGGTCTCAGCCAACTGGTTCCCCAGTGGATTGAGACCGATGAGACGATGC |
| 16348 | fw_hMRP_pt4aYTK | GCATCGTCTCATCGGTCTCAATCCAGAAGCGTATCCCGCTGAGCTGGCTGAGACCTGAGACGGCAT |
| 16349 | rv_hMRP_pt4aYTK | ATGCCGTCTCAGGTCTCAGCCAGCTCAGCGGGATACGCTTCTGGATTGAGACCGATGAGACGATGC |
| 16350 | fw_yMRP_pt4aYTK | GCATCGTCTCATCGGTCTCAATCCGTACCTCTATTGCAGGGTACTGGCTGAGACCTGAGACGGCAT |
| 16351 | rv_yMRP_pt4aYTK | ATGCCGTCTCAGGTCTCAGCCAGTACCCTGCAATAGAGGTACGGATTGAGACCGATGAGACGATGC |
| 16352 | fw_tracrRNA_pt4aYTK | GCATCGTCTCATCGGTCTCAATCCGTTTTAGAGCTAGAAATAGCAAG |
| 16353 | rv_tracrRNA_pt4aYTK | ATGCCGTCTCAGGTCTCAGCCAGCACCACCGACTCGGTG |
| 16643 | fw_mito-mTq2_YTKp3b | GCATCGTCTCATCGGTCTCATTCTATGGTATCAAAAGGTGAAG |
| 16644 | fw_mito-mTq2_YTKp3 | GCATCGTCTCATCGGTCTCATATGGTATCAAAAGGTGAAG |
| 16645 | rv_mito-mTq2 | ATGCCGTCTCAGGTCTCAGGATTTATTTATATAATTCATCCATACCTAATG |
| 16837 | Fw_pUDE929-HDV_YTK4 | GATCGGTCTCAATCCGGCCGGCATGGTCCCAGCC |
| 16838 | Fw_pUDE929-HDV_YTK4b | GATCGGTCTCATGGCGGCCGGCATGGTCCCAGCC |
| 16839 | Rv_pUDE929-HH_YTK4 | GATCGGTCTCACATAATCTAATGACGAGCTTACTCG |
| 16840 | Rv_mito-mTq2_pt3a | ATGCCGTCTCAGGTCTCAAGAATTATTTATATAATTCATCCATACCTAATG |
| 16841 | Fw_mito-mTq2-pt3a | GCATCGTCTCATCGGTCTCAATCCGTATCAAAAGGTGAAG |
| 16842 | Rv_mito-mTq2-pt4a | ATGCCGTCTCAGGTCTCAGCCATTATTTATATAATTCATCCATACCTAATG |
| 18440 | Fw_pUDC329_HDV | TGGGCAACACCTTCGGGTGGCGAATGGGACTAACTCGAGAGTGCTTTTAACTAAGAATTATTAGTCTTTTCTGC |
| 18441 | Rv_pUDC329_HH | GTTTCGTCCTCACGGACTCATCAGATGCGCATTTGTAATTAAAACTTAGATTAGATTGCTATGC |
| 18442 | Fw_mitoARG8-HH | GCGCATCTGATGAGTCCGTGAGG |
| 18443 | Rv_mitoARG8_HDV | GTCCCATTCGCCACCCGAAGGTGTTGC |
| 18444 | Fw_mitoARG8-dg | GGTGAAGGTGGTGTATTTCC |
| 18445 | Rv_mitoARG8-dg | AAAGCACCCATTGTTCTACC |
| 18446 | Fw_F1v2-pt3a | GGTCTCATATGGGTTCGAGCCCCCTACAGGGCTCCATTCTAGAGACC |
| 18447 | Rv_F1v2-pt3a | GGTCTCTAGAATGGAGCCCTGTAGGGGGCTCGAACCCATATGAGACC |
| 18448 | Fw_tRK2-tr93_pt3a | GGTCTCATATGGCCTTGTTAGCTCAGTTGGTAGAGCGTTCGGCTTTTAACCGAAATGTCAGGGGTTCGAGCCCCCTATGAGGCTCCATTCTAG AGACC |
| 18449 | Rv_tRK2-tr93_pt3a | ggtctctagaatggAGCCTCATAGGGGGCTCGAACCCCTGACATTTCGGTTAAAAGCCGAACGCTCTACCAACTGAGCTAACAAGGCcatatgagacc |
| 14022 | Fw_pUDC286_YPRCtau3 | AGAATGATTTACAATCTAGTCGCAAAAACAAGTACAGTGCTGACGTCCCATCTTTTAATGCATGCGCGGATGACACGAACTCA |
| 14023 | rv_pUDC286_YPRCtau3 | CATTACCAATGAATGCTGTTTTTGCAGAAATAACGAGATATCTGCAATAAAAGCAAAAGTCCTTGATCTGTCGGGTGTCTGCT |

* The following consensus is used for primer names: [fw/rv]_[Gene target]_[homology flank or part type]. Fw indicates a forward primer, Rv a reverse primer.  Dg is a diagnostic primer.

Table S3. All non-silent mutations observed in IMS1248 (population bioreactor A) and IMS1249 (population Bioreactor B) and IMS1251-1254 (single colony isolates bioreactor B). A '+' indicates a non-silent mutation was observed in the respective strain.

| location | Systemic name | Gene name | Annotation | Subcellular localization | IMS1248 | IMS1249 | IMS1251 | IMS1252 | IMS1253 | IMS1254 |
|---|---|---|---|---|---|---|---|---|---|---|
| **CHRII** | YBL101C | *ECM21* | Protein ECM21 | Cytosol | - | - | - | - | + | - |
| **CHRII** | YBR102C | *EXO84* | Exocyst complex component EXO84 | Cell Periphery | + | + | + | + | + | + |
| **CHRII** | YBR297W | *MAL33* | Maltose fermentation regulatory protein MAL33 | Nucleus | + | + | + | + | + | + |
| **CHRII** | YBR098W | *MMS4* | Crossover junction endonuclease MMS4 | Nucleus | + | + | + | + | + | + |
| **CHRII** | YBR028C | *YPK3* | AGC kinase YPK3 | Nucleus | + | + | + | - | + | - |
| **CHRIII** | YBR298C | *MAL31* | Maltose permease MAL31 | Cell Periphery | + | + | + | + | + | + |
| **CHRIII** | YCR039C | *MATALPHA2* | Mating-type protein ALPHA2 | Nucleus | + | + | + | + | + | + |
| **CHRIV** | YDL017W | *CDC7* | Cell division control protein 7 | Nucleus | + | + | + | + | + | + |
| **CHRIV** | YDR097C | *MSH6* | DNA mismatch repair protein MSH6 | Nucleus | + | + | + | + | + | + |
| **CHRIV** | YDR006C | *SOK1* | Protein SOK1 | Nucleus | + | + | + | + | + | + |
| **CHRV** | YER017C | *AFG3* | Mitochondrial respiratory chain complexes assembly protein AFG3 | Mitochondria | + | + | + | + | + | + |
| **CHRV** | YER125W | *RSP5* | E3 ubiquitin-protein ligase RSP5 | Nucleus | + | - | - | - | - | - |
| **CHRV** | YHL049C | | Uncharacterized protein YHL049C | | + | + | + | + | + | + |
| **CHRVI** | YFR016C | | Uncharacterized protein YFR016C | | + | + | + | + | + | + |
| **CHRVII** | YGR125W | | Uncharacterized vacuolar membrane protein YGR125W | Vacuole | + | - | + | - | - | - |
| **CHRVII** | YPR202W | | Putative uncharacterized protein YPR202W | | + | + | + | + | + | + |
| **CHRVIII** | YHR055C | *CUP1-2* | Copper metallothionein 1-2 | Cytosol | + | + | + | + | + | + |
| **CHRVIII** | YHR056C | *RSC30* | Chromatin structure-remodeling complex protein RSC30 | Nucleus | + | + | + | + | + | + |
| **CHRIX** | YIL130W | *ASG1* | Activator of stress genes 1 | Nucleus | - | + | - | + | - | - |
| **CHRIX** | YIL128W | *MET18* | DNA repair/transcription protein MET18/MMS19 | Nucleus | + | + | + | + | + | + |

| | | | | | | | | | | |
|---|---|---|---|---|---|---|---|---|---|---|
| **CHRIX** | YJL222W | *VTH2* | VPS10 homolog 2 | Vacuole | + | + | + | + | + | + |
| **CHRIX** | YNR065C | | Uncharacterized membrane glycoprotein YNR065C | ER | + | + | + | + | + | + |
| **CHRX** | YJR005W | *APL1* | AP-2 complex subunit beta | Cell Periphery | + | + | + | + | + | + |
| **CHRX** | YJL090C | *DPB11* | DNA replication regulator DPB11 | Nucleus | + | + | + | + | + | + |
| **CHRXI** | YKL122C | *SRP21* | Signal recognition particle subunit SRP21 | Cytoplasm | + | + | + | + | + | + |
| **CHRXI** | YBL113C | | Uncharacterized protein YBL113C | | + | + | + | + | + | + |
| **CHRXI** | YKR015C | | Uncharacterized protein YKR015C | | + | + | + | + | + | + |
| **CHRXII** | YLR160C | *ASP3-4* | L-asparaginase 2-4 | Cell Periphery | + | + | + | + | + | + |
| **CHRXII** | YLR231C | *BNA5* | Kynureninase | Nucleus | + | + | + | + | + | + |
| **CHRXII** | YLR422W | *DCK1* | DOCK-like protein YLR422W | Mitochondria | + | + | + | + | + | + |
| **CHRXII** | YLR424W | *SPP382* | Pre-mRNA-splicing factor SPP382 | Nucleus | + | + | + | + | + | + |
| **CHRXII** | YLR024C | *UBR2* | E3 ubiquitin-protein ligase UBR2 | Cytoplasm | - | + | + | + | + | + |
| **CHRXII** | YEL074W | | Putative UPF0320 protein YEL074W | | + | + | + | + | + | + |
| **CHRXIII** | YML054C | *CYB2* | Cytochrome b2, mitochondrial | Mitochondria | + | + | + | + | + | + |
| **CHRXIII** | YMR323W | *ERR3* | Enolase-related protein 3 | Cytosol | + | + | + | + | + | + |
| **CHRXIII** | YMR263W | *SAP30* | Transcriptional regulatory protein SAP30 | Nucleus | + | + | + | - | + | + |
| **CHRXIV** | YNL192W | *CHS1* | Chitin synthase 1 | Cell Periphery | + | + | + | + | + | + |
| **CHRXIV** | YFL060C | *SNO3* | Probable glutamine amidotransferase SNO3 | Cytosol | + | + | + | + | + | - |
| **CHRXIV** | YNL333W | *SNZ2* | Probable pyridoxine biosynthesis protein SNZ2 | ND | + | + | + | + | + | + |
| **CHRXIV** | YNL186W | *UBP10* | Ubiquitin carboxyl-terminal hydrolase 10 | Nucleus | + | + | + | + | + | + |
| **CHRXIV** | YNL064C | *YDJ1* | Mitochondrial protein import protein MAS5 | Cytosol | + | + | + | + | + | + |
| **CHRXV** | YOR239W | *ABP140* | Uncharacterized methyltransferase ABP140 | Cell Periphery | + | + | + | + | + | + |

| | | | | | | | | | | |
|---|---|---|---|---|---|---|---|---|---|---|
| **CHRXV** | YOR390W | *FEX1* | UPF0695 membrane protein YOR390W | Cell Periphery | + | + | + | + | - | - |
| **CHRXV** | YOR120W | *GCY1* | Protein GCY | Nucleus | + | + | + | + | + | + |
| **CHRXV** | YOR234C | *RPL33B* | 60S ribosomal protein L33-B | Cytosol | + | + | + | + | + | + |
| **CHRXV** | YOR305W | *RRG7* | Required for respiratory growth protein 7, mitochondrial | Mitochondria | + | + | + | + | + | + |
| **CHRXV** | YOR389W | | Uncharacterized protein YOR389W | | - | - | - | - | + | + |
| **CHRXVI** | YPL160W | *CDC60* | Leucine--tRNA ligase, cytoplasmic | Cytoplasm | + | + | + | + | + | + |
| **CHRXVI** | YPL281C | *ERR2* | Enolase-related protein 2 | Cytosol | + | + | + | + | + | + |
| **CHRXVI** | YOR388C | *FDH1* | Formate dehydrogenase 1 | Cytosol | + | + | + | + | + | + |
| **CHRXVI** | YPR106W | *ISR1* | Serine/threonine-protein kinase ISR1 | ND | + | + | + | + | + | + |
| **CHRXVI** | YPL082C | *MOT1* | TATA-binding protein-associated factor MOT1 | Nucleus | + | + | + | + | + | + |
| **CHRXVI** | YPR075C | *OPY2* | Protein OPY2 | Vacuole | + | + | + | + | + | + |
| **CHRXVI** | YPL085W | *SEC16* | COPII coat assembly protein SEC16 | ER | + | + | + | + | + | + |
| **CHRXVI** | YPL032C | *SVL3* | Styryl dye vacuolar localization protein 3 | Cell Periphery | + | + | + | + | + | + |
| **CHRXVI** | YPR080W | *TEF1* | Elongation factor 1-alpha | Cytosol | + | + | + | + | + | + |
| **CHRXVI** | YPR163C | *TIF3* | Eukaryotic translation initiation factor 4B | Cytoplasm | - | + | - | + | + | - |
| **CHRXVI** | YHR219W | | Putative uncharacterized protein YHR219W | | + | + | + | + | + | + |
| **CHRXVI** | YPL025C | | Putative uncharacterized protein YPL025C | | + | + | - | + | + | + |
| **CHRXVI** | YPL283W-A | | Putative uncharacterized protein YPL283W-A | | + | - | + | + | - | + |

**Table S4. Non-synonymous mutations found in populations IMS1248 and IMS1249 and single colonies IMS1251 - IMS1254.** Mutations that were found in both IMS1248 and IMS1249 were excluded as these could originate from the preculture. A full list of all mutations can be found in Table S3 and SI file 1.

| Location | Position | Systemic name | Gene name | Description | Subcellular localization | Mutation | Effect on protein | Found in strains: |
|---|---|---|---|---|---|---|---|---|
| **CHR II** | 28040 | YBL101C | *ECM21* | Alpha-arrestin, ubiquitin ligase adaptor for Rsp5p; regulates starvation- and substrate-induced Ub-dependent endocytosis of select plasma membrane localized amino acid transporters | Cytosol | Insertion (2 bp) | Early termination (198 aa) | IMS1253 |
| **CHR V** | 412435 | YER125W | *RSP5* | NEDD4 family E3 ubiquitin ligase | Cytosol | $Leu_{518}Ser$ | Amino acid substitution | IMS1248 |
| **CHR IX** | 99639 | YIL130W | *ASG1* | Activator of stress genes 1, Zinc cluster protein proposed to be a transcriptional regulator; regulator involved in the stress response; regulates utilization of fatty acids and accumulation of lipids | Nucleus | Insertion (9 bp) | Insertion 3x Asn | IMS1249, IMS1252 |
| **CHR XII** | 174490 | YLR024C | *UBR2* | E3 ubiquitin-protein ligase component of the Mub1p-Ubr2p-Rad6p ubiquitin ligase complex required for the ubiquitination and degradation of Rpn4p | Cytoplasm | $STOP_{1283}Tyr$ | Elongation (27 aa) | IMS1249, IMS1251, IMS1252, IMS1253, IMS1254 |
| **CHR XV** | 1069766 | YOR389W | | Putative protein of unknown function; expression regulated by copper levels | | $Asn_{437}Asp$ | Amino acid substitution | IMS1253, IMS1254 |
| **CHR XVI** | 871051 | YPR163C | *TIF3* | Translation initiation factor eIF-4B; RNA recognition motif containing single-stranded RNA binding protein that possesses RNA annealing and strand-exchange activities | Cytoplasm | Deletion (1 bp) | Early termination (348 aa) | IMS1249, IMS1252, IMS1253 |

### Supplementary figures

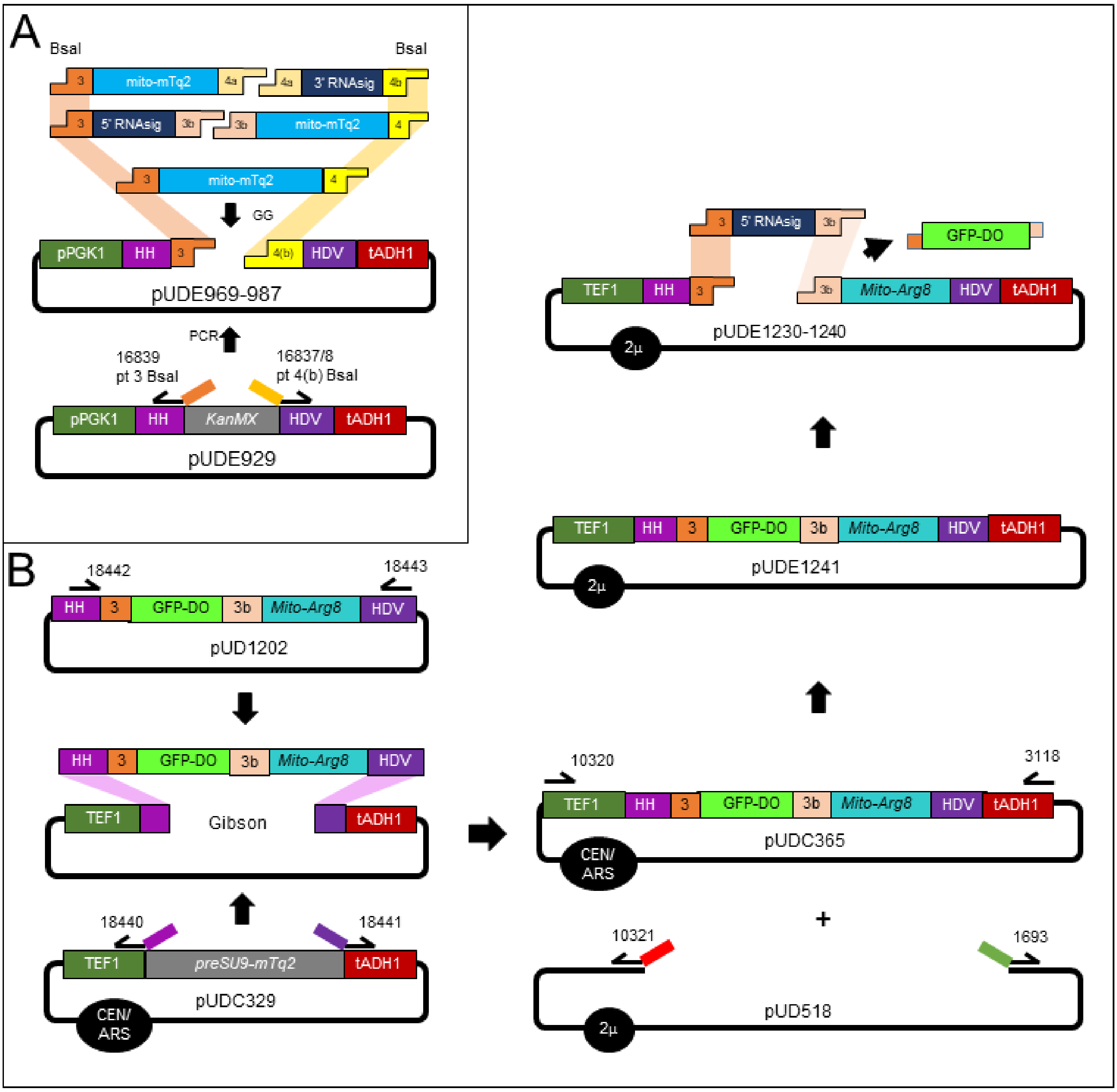


Figure S1. Construction strategy of libraries of mito-mTq2 mRNA (A) and mito-Arg8 mRNA (B) with different RNA import signals.

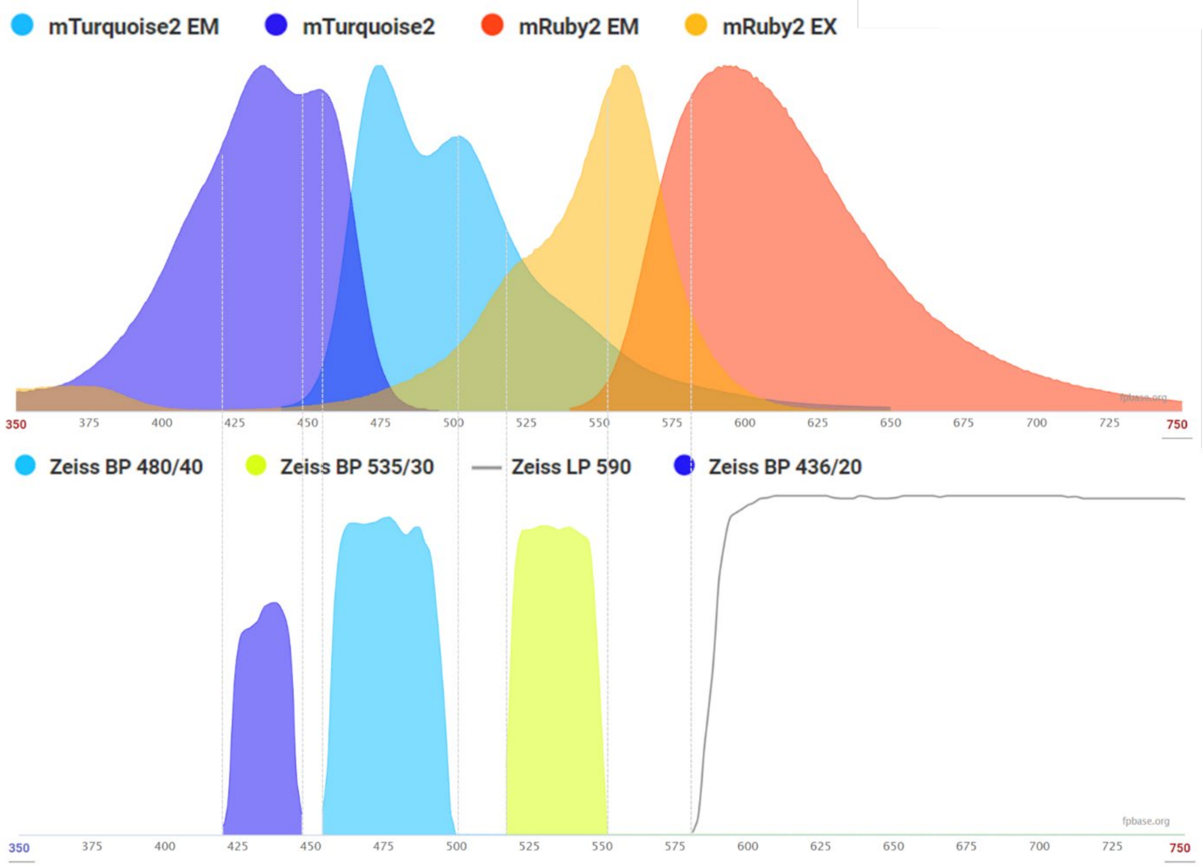


**Figure S2. Spectral overview of fluorophores and filters used for *in vivo* co-localization assays.** The x-axes displays wavelength in nanometers the spectral plot is generated using FPBase.

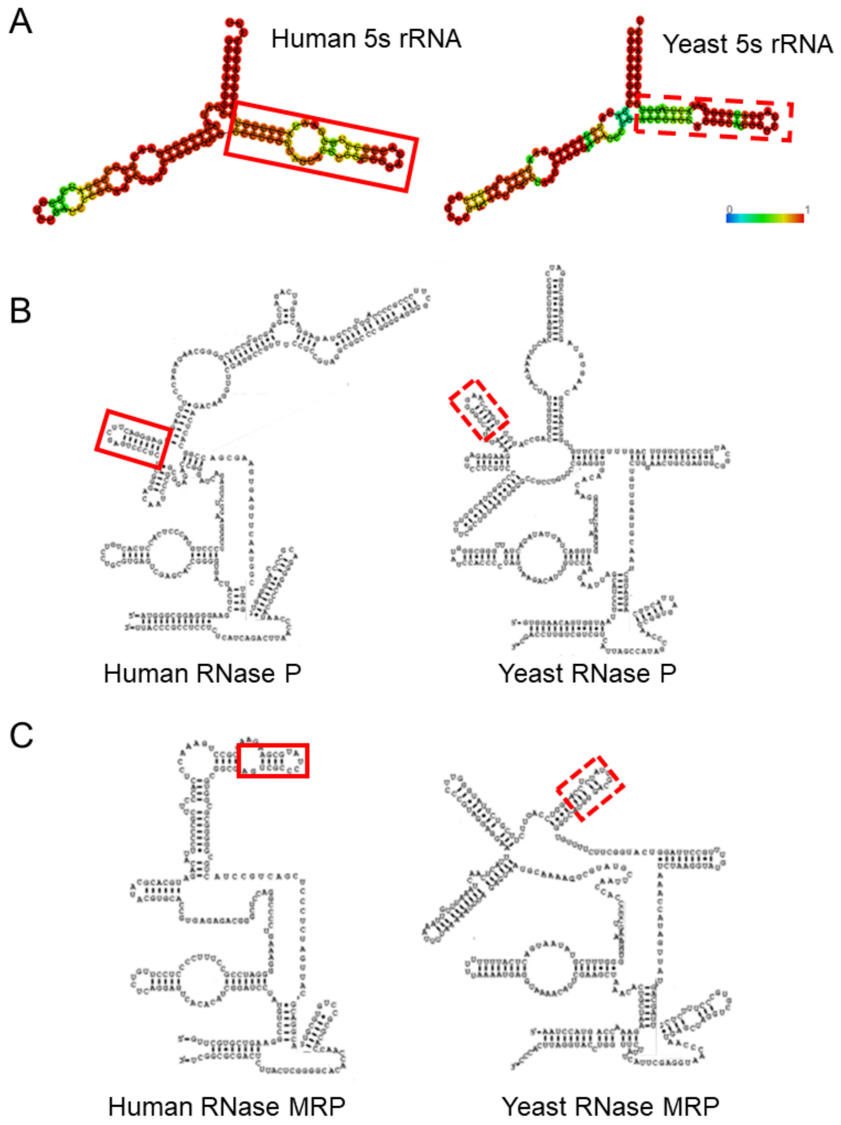


**Figure S3. Structural comparison between human and *S. cerevisiae* putative RNA import signals.** A) human 5S rRNA (accession no. NR_023363) and *S. cerevisiae* 5S rRNA (accession no. NR_132215.1). The putative h5S rRNA MAM as reported Zelenka, Alán (2) is displayed in a solid red box. In order to extrapolate the h5S rRNA stem to a yeast equivalent, a comparative analysis was done between the full 5S rDNA sequences as found in and in yeast. Global alignment of these two exactly 121 nt sequences scored 75% similarity. A structural comparison of these native 5S rRNA components revealed similar structures as well. In h5S rRNA the sequence of the putative RNA signal mapped to a highly similar secondary structure found in yeast 5S rRNA, displayed in a dashed red box. Base-pairing probabilities are denoted according to the color scale as indicated. Structural analysis was done using RNAfold (Vienna RNA secondary structure server *[12, 13]*). B) Structural comparison between RNA components of human RNase P and *S. cerevisiae* RNase P and C) human RNase MRP and *S. cerevisiae* RNase MRP. Putative hRP and hMRP stems as reported by Wang et al. (2010) *[3]* are indicated with a solid red box. Subsequent mapping of this substructure to the yeast RNA variant, allowed for extrapolation of the putative signals. Apart from structural similarity, information on identified conserved intramolecular RNA domains justify the choice of specific regions as extrapolated signals to be examined for ability to stimulate RNA transport to the mitochondria. Based on structural similarity, and identified conserved intramolecular domains, the putative yeast equivalents are displayed in a dashed red box. Adapted from Esakova and Krasilnikov (14).

A - Yeast cytosolic

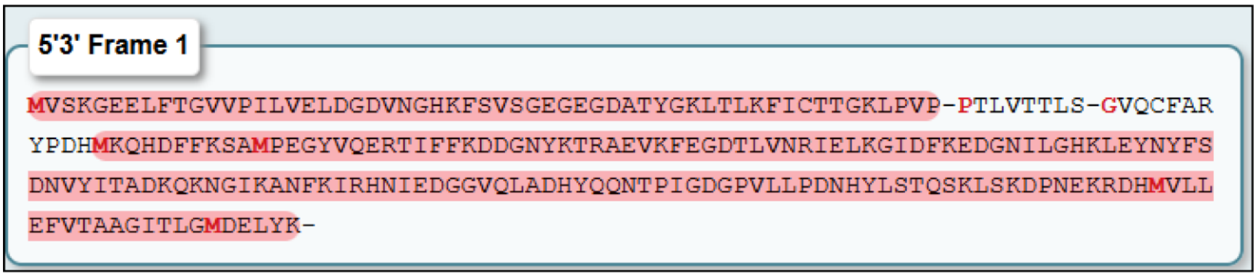


B - Yeast mitochondrial

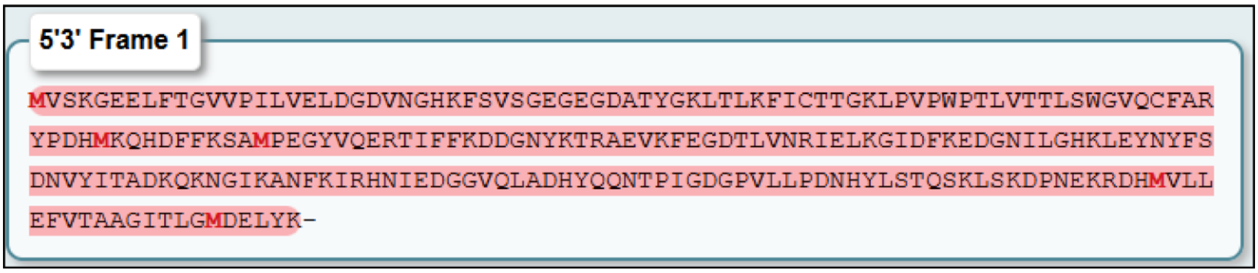


**Figure S4. Translation frames of mito-mTq2 mRNA.** Open reading frames are indicated in red, potential START codons are indicated with a red bold-face M, STOP codons are indicated with a hyphen (-).Top: 5' – 3' Frame 1 translation of mito-mTq2 using yeast cytosolic genetic codon usage. The translation product from the first methionine start codon is 57 amino acids and has a weight of 6 kDa. The translation product starting at the second methionine is 161 amino acids and has a weight of approximately 18.5 kDa. Bottom: 5' – 3' Frame 1 translation of mito-mTq2 using yeast mitochondrial genetic codon usage. The full translation product is 240 amino acids, and has a weight of 27 kDa. Tranlsations were generated using Expasy [15].

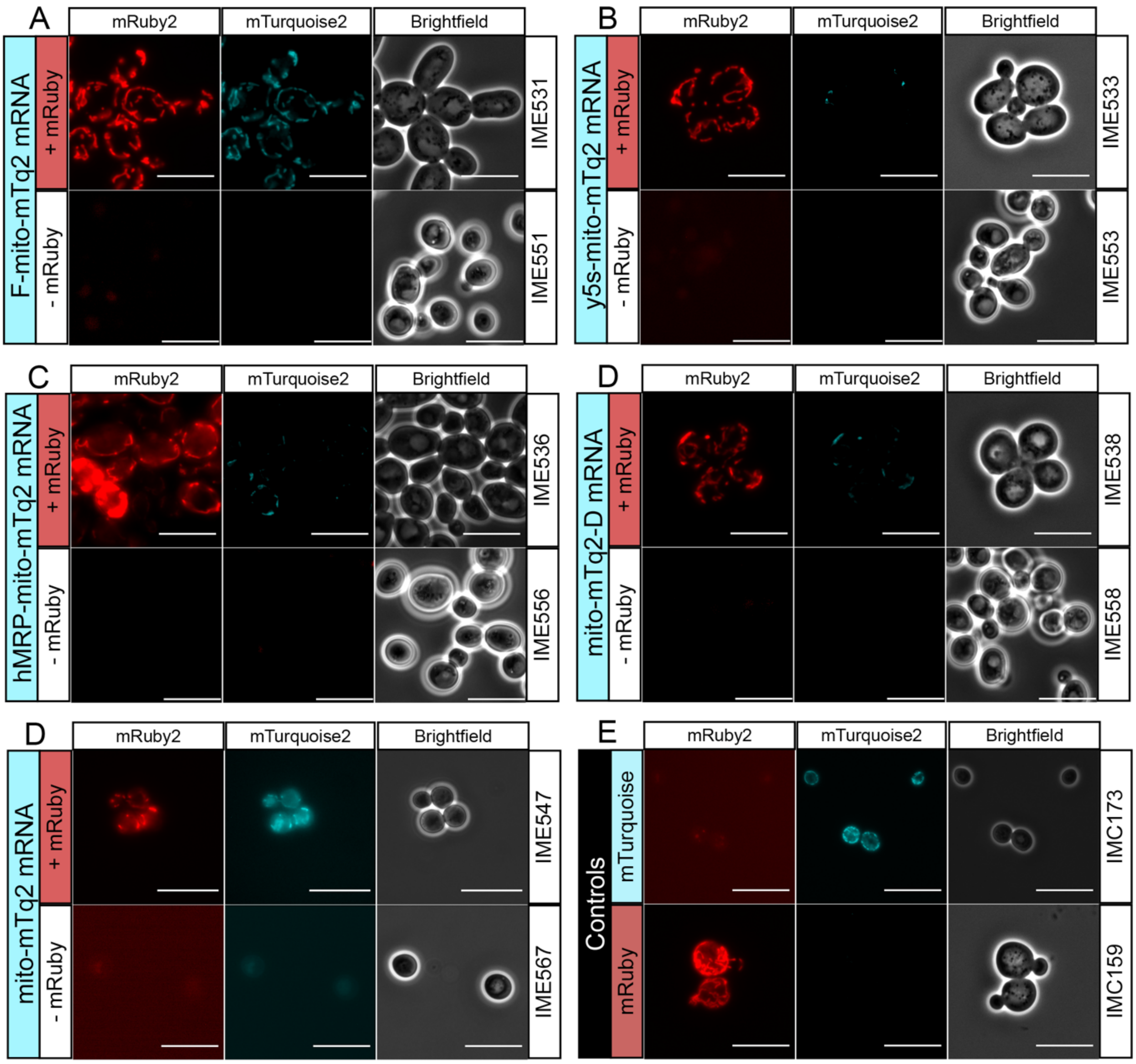


**Figure S5. Fluorescence microscopy images (1000x) in red (mRuby) and blue (mTurquoise) channels and brightfield images of strains expressing mito-mTq2 fused to different mitochondrial RNA import signals.** The fluorescence of strains expressing each signal was measured either in the presence (IME531-IME547, panels A to E, "+mRuby") or absence (IME551-IME567, panels A to E "-mRuby") of mRuby2, along with a positive control targeting the preCOX4-mTq2 protein to the mitochondria (IMC173, panel E, mTurquoise), a negative control omitting the mito-mTq2 RNA (IMC159, panel E, mRuby). All pictures were obtained with the same settings and an exposure times of 1200 ms for mTq2 and 500 ms for mRuby2. Images were processed using the same settings for brightness and contrast, where any pixel values of lower than 1000 were omitted to remove background fluorescence. Scale bars represent 10 µm.

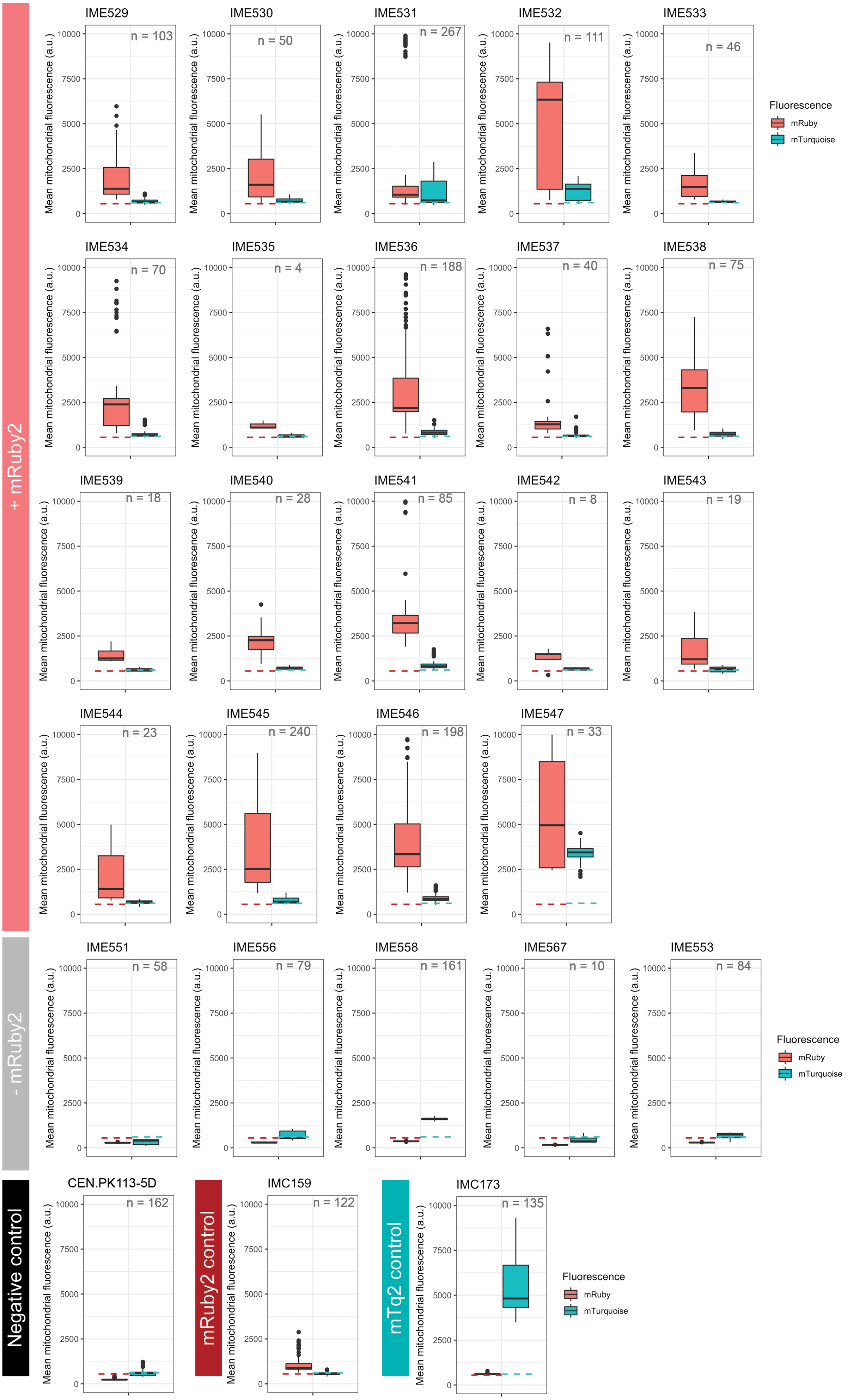
+ mRuby2
- mRuby2
Negative control
mRuby2 control
mTq2 control
IME529
n = 103
IME530
n = 50
IME531
n = 267
IME532
n = 111
IME533
n = 46
IME534
n = 70
IME535
n = 4
IME536
n = 188
IME537
n = 40
IME538
n = 75
IME539
n = 18
IME540
n = 28
IME541
n = 85
IME542
n = 8
IME543
n = 19
IME544
n = 23
IME545
n = 240
IME546
n = 198
IME547
n = 33
IME551
n = 58
IME556
n = 79
IME558
n = 161
IME567
n = 10
IME553
n = 84
CEN.PK113-5D
n = 162
IMC159
n = 122
IMC173
n = 135
Mean mitochondrial fluorescence (a.u.)
0
2500
5000
7500
10000
Fluorescence
mRuby
mTurquoise

**Figure S6. Distribution of the mean mitochondrial fluorescence in the different strains in the mRuby2 (red) and mTurquoise2 (blue) channels, determined from microscopic analysis and measuring mean pixel intensities in the red and blue channels.** The distribution of the mean fluorescence intensity of mRuby (red) and mTurquoise (blue) is presented as boxplots, where the line represents the median value, and the edges of the boxes represent the 25$^{th}$ and 75$^{th}$ percentile. Outliers are shown as dots. The dashed line represents the median fluorescence value of the negative controls as a reference (IMC173 for ruby fluorescence, IMC159 for mTq2 fluorescence). The strains presented contain mito-mTq2 fused to the same RNA targeting signal either in the presence (IME529-IME547) or absence (IME551-IME567) of mRuby2.

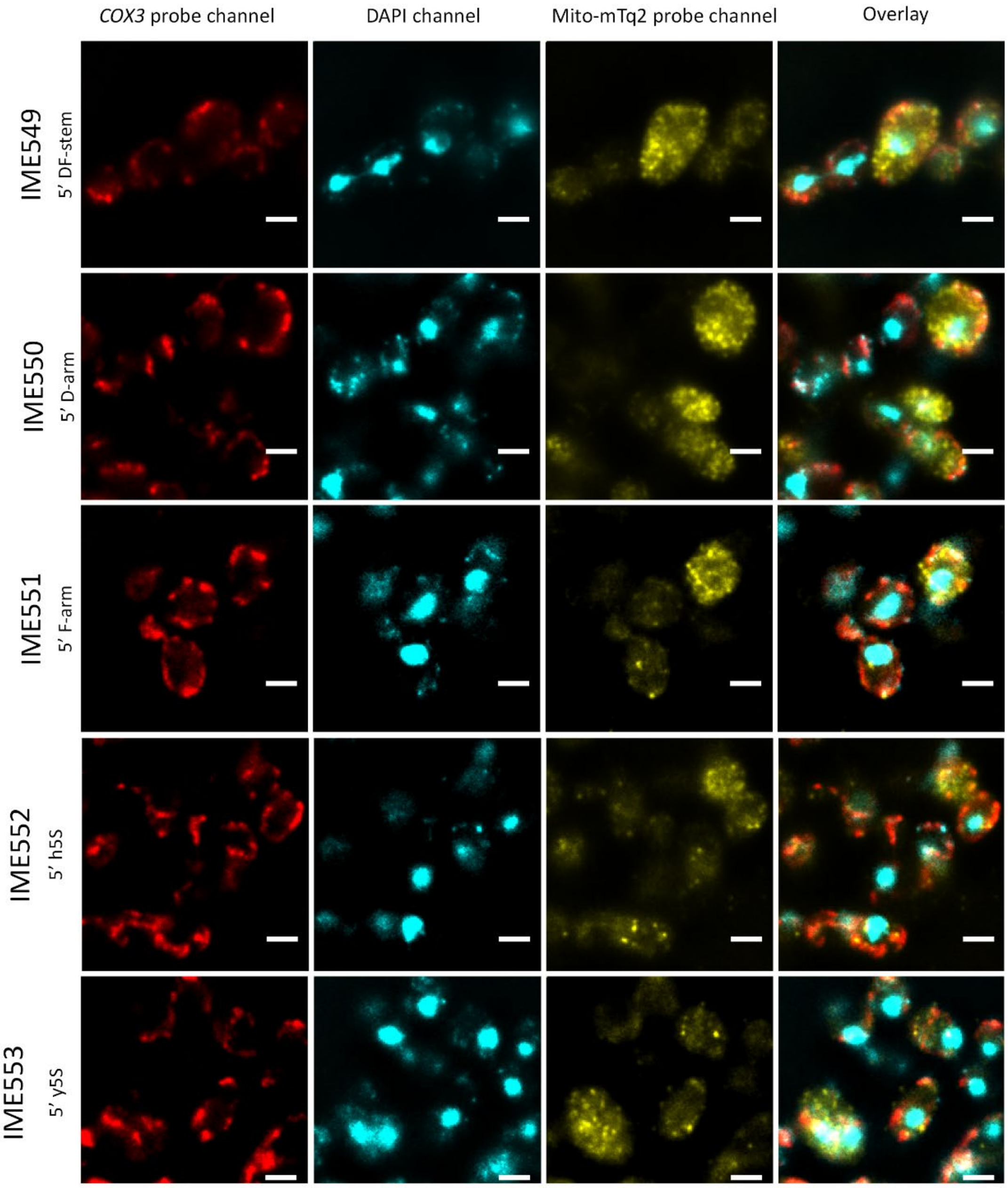


(*Figure continues on next page)*

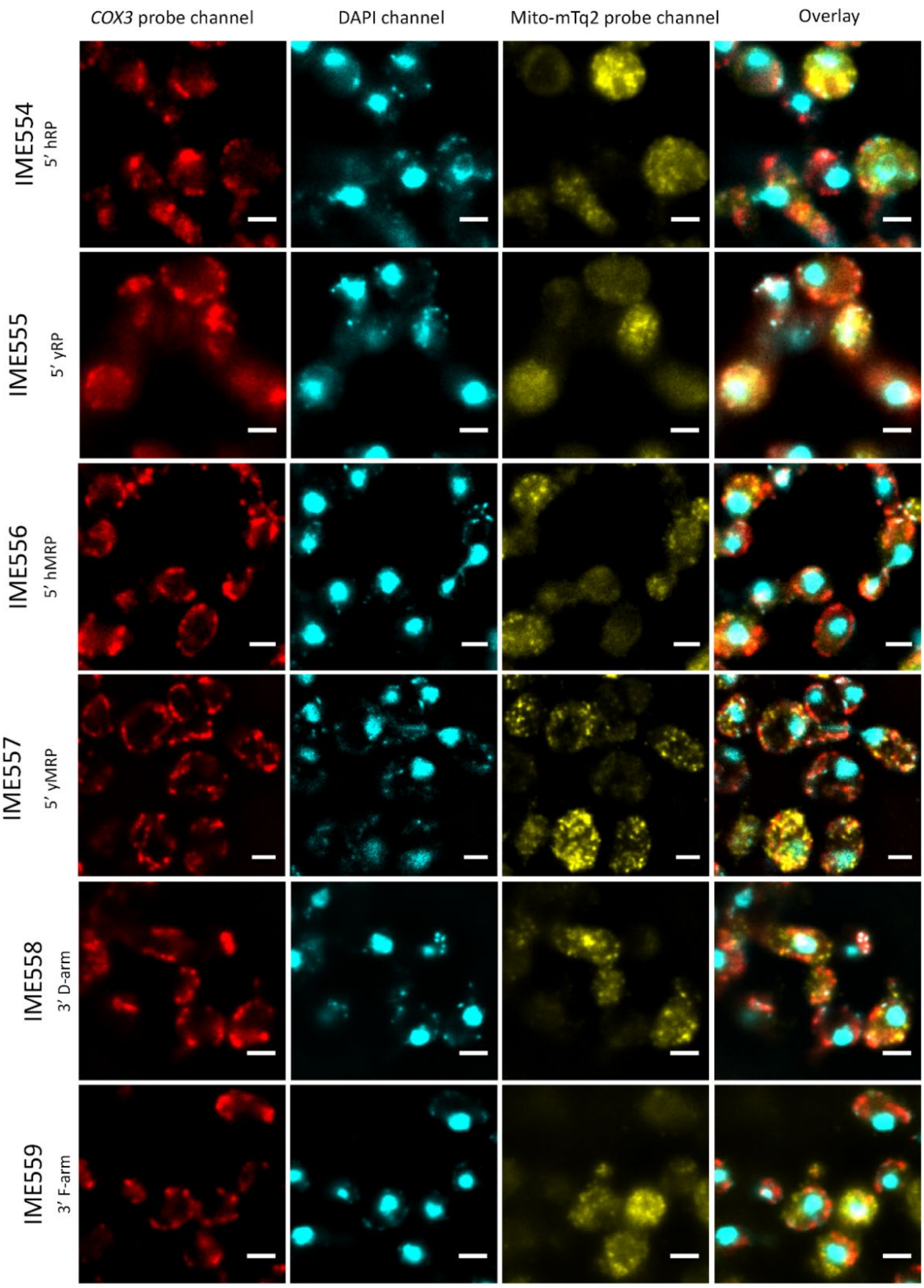


(*Figure continues on next page)*

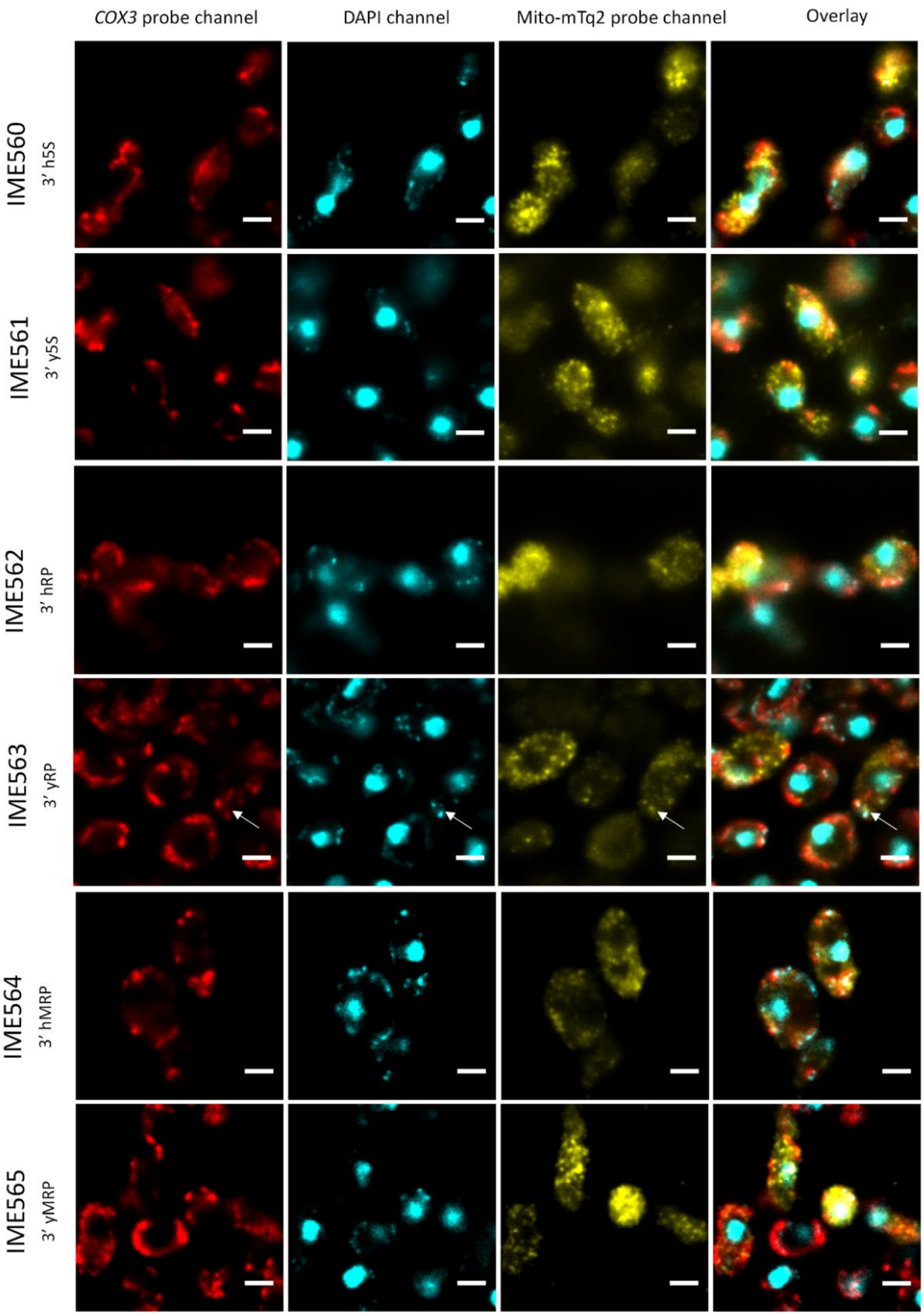


(*Figure continues on next page)*

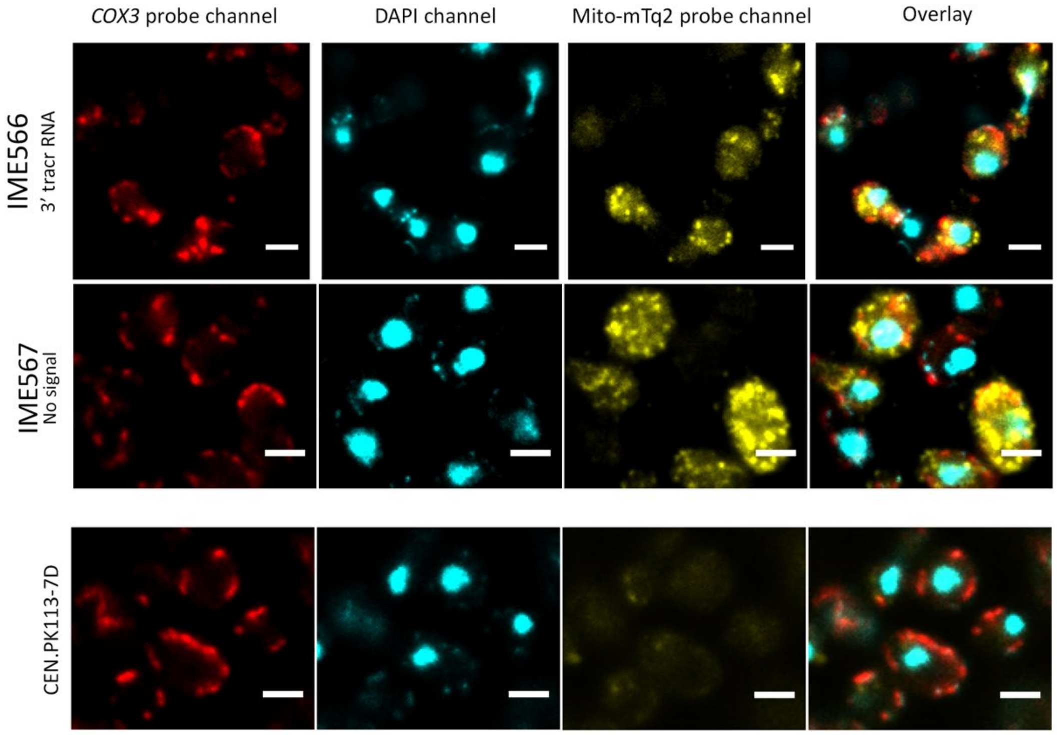


**Figure S7. FISH analysis of strains expressing mito-mTq2 mRNA with different RNA signals and negative control CEN.PK113-7D, omitting mito-mTq2 mRNA.** Mitochondrially transcribed *COX3* mRNA was targeted with a specific probe labeled with CAL Fluor Red 635, while mito-mTq2 mRNA was targeted with a specific probe labeled with Quasar-570. DNA was stained with a fluorescent DNA-specific DAPI stain. Acquisition in the *COX3* probe channel (red), mito-mTq2 probe (yellow) and the DAPI channel (blue) were respectively done with exposure times of 1000, 600 and 100 seconds, at a magnification of 100x. Every image was derived from a single slice in a Z-stack after iterative deconvolution. The white scale bar indicates 5 µm.

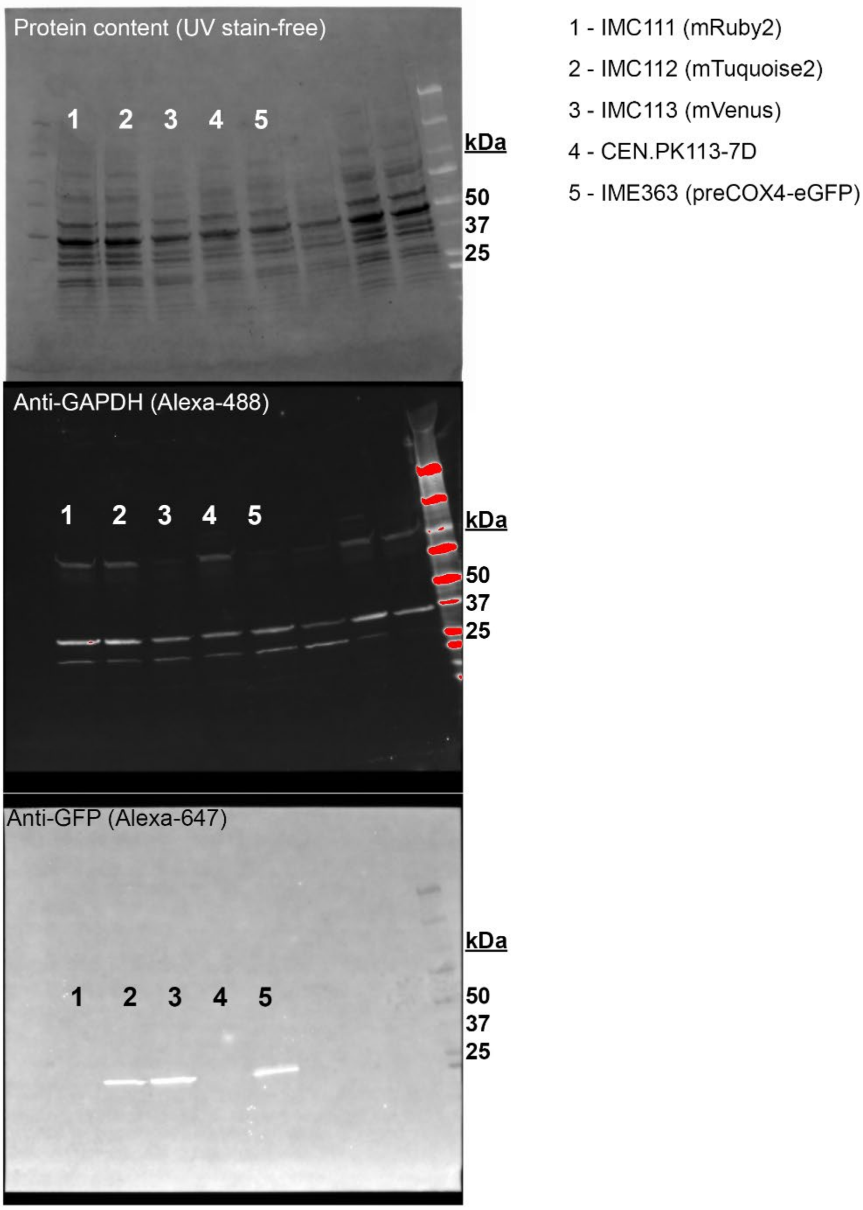


**Figure S8. Western blot of fluorescent proteins using anti-GFP antibody.** Positive control strain IME363 expressing eGFP (26.9 kDa) was compared to control strains IMC112 and IMC113 resp. expressing mTq2 (26.9 kDa) and Venus (26.8 kDa) for capability of anti-GFP to recognize the GFP-derivatives. Negative control strain IMC111 expressing mRuby2 (26.5 kDa) was not recognized by anti-GFP (bottom). 25 µg protein was loaded in each lane at the start of PAGE (Top), anti-GAPDH (middle) was used as a protein loading control ($MW_{GAPDH}$= 36 kDa). The reference ladder displays protein size in kDa.

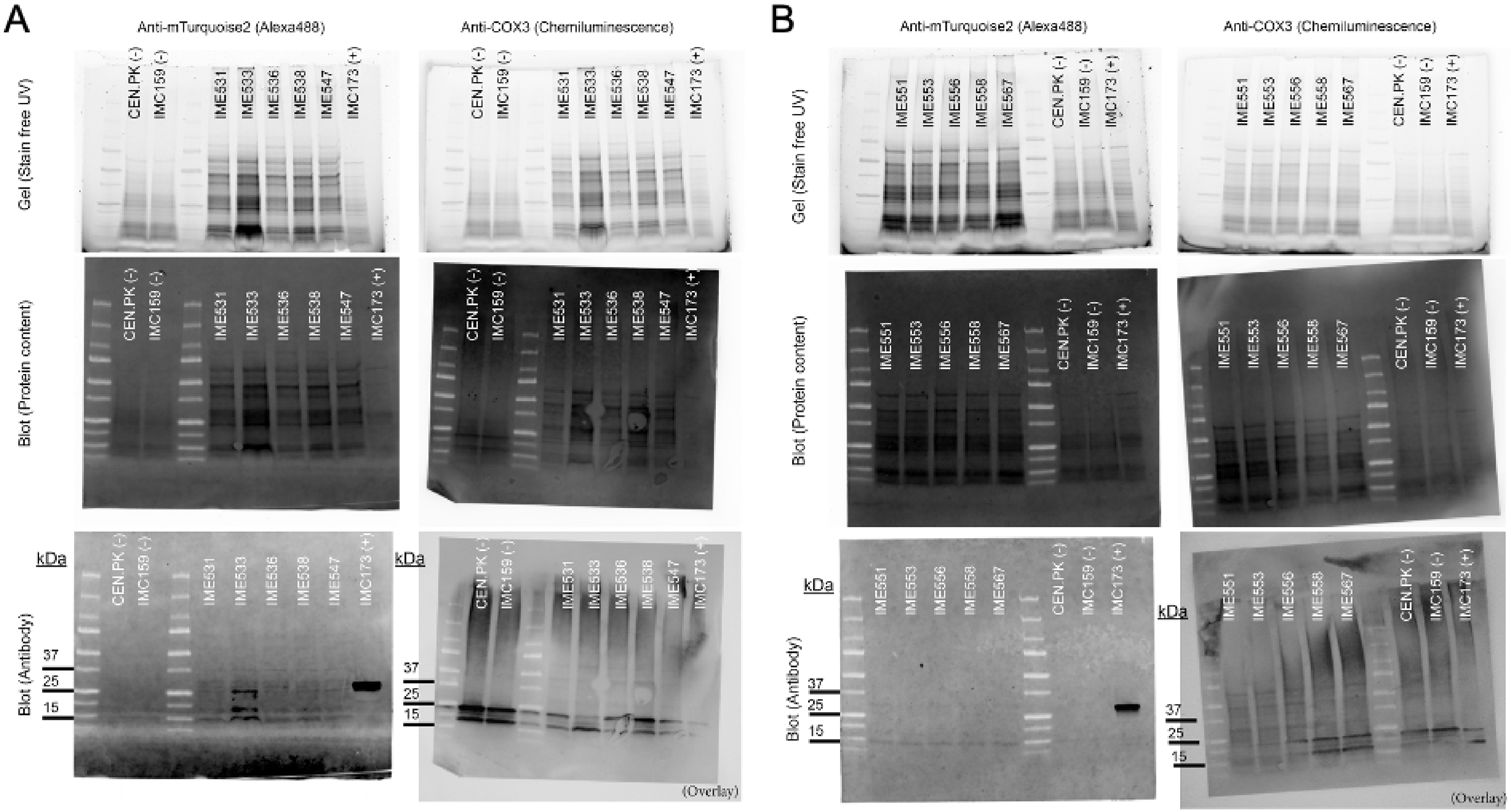


**Figure S9 Western blot analysis of mTurquoise2 expression in strains expressing mito-mTq2 mRNA in with the presence (panel A) or absence (panel B) of the preCox4-mRuby2 protein.** Negative controls (-) include CEN.PK113-7D (no fluorescent protein) and IMC159 (preCOX4-mRuby2 only), and the positive control IMC173 indicated with (+) expresses preSU9-mTq2. Top row: full protein content of isolated mitochondria separated on an SDS-PAGE gel. Middle row: full protein content of isolated mitochondria transferred on a blotting membrane. Bottom row: membranes blotted with antibody and imaged with the corresponding settings. The left column shows isolated mitochondria blotted with an anti-GFP antibody conjugated with an Alexa-488 probe, imaged at 488 nm, the right column shows blotted with an anti-Cox3p (subunit 3 of the cytochrome c oxidase of the respiratory chain) antibody-HRP complex, treated with substrate and imaged using chemiluminescence as a loading control. The chemiluminescence image was overlaid with the blot to allow for visibility of the ladder. Approximate protein sizes of the observed bands in kDa are indicated on the left, the used ladder was the Precision Plus Protein Dual Color Standards (Bio-Rad). Each lane was loaded with 25 μg protein, except for IMC173, that was loaded with 10 μg protein. Expected protein sizes are 26.9 kDa for mTq2 and 30 kDa for Cox3p.

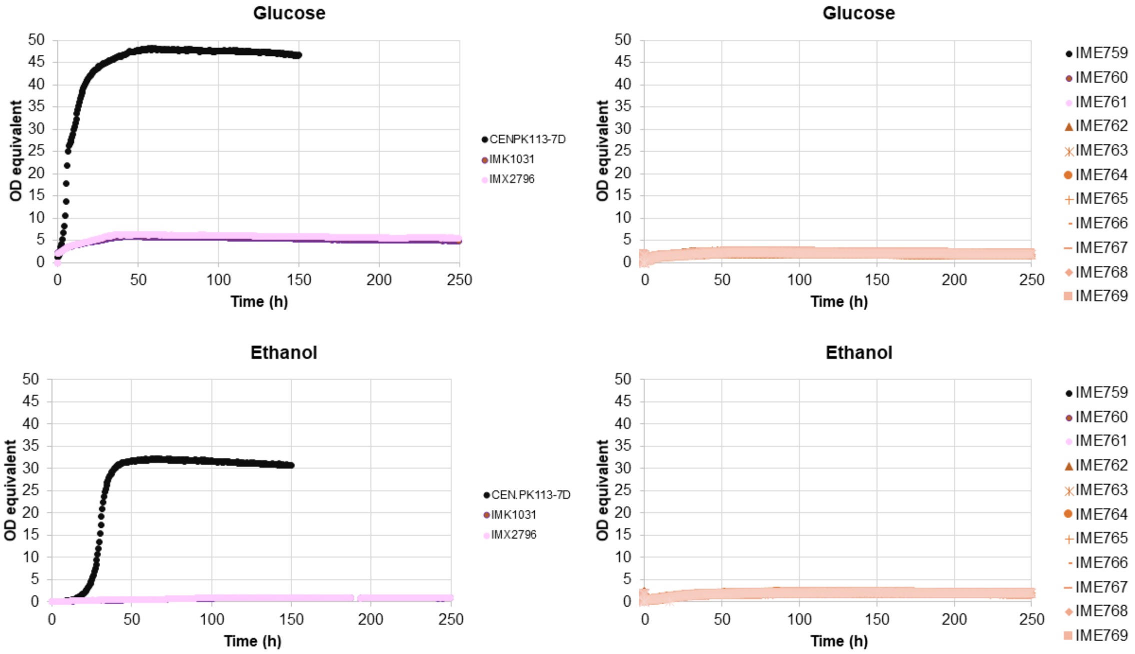


**Figure S10 Growth of strains with mito-ARG8 mRNA targeted to mitochondria, on synthetic medium without arginine in microtiterplates.** A and B, controls strains CEN.PK113-7D (prototrophic reference) and the arginine auxotrophs IMK1031 (*Δarg8 Δcar2*) and IMX2796 (*(Δarg8 Δcar2 YPRCtau3::preCox4-mRuby2*) grown with glucose (A) or ethanol (C) as sole carbon source. C and D, strains IME759-IME769 which are IMX2796 transformed with different plasmids expressing mito-ARG8 mRNA with different targeting signals, grown on glucose (C) or ethanol (D)

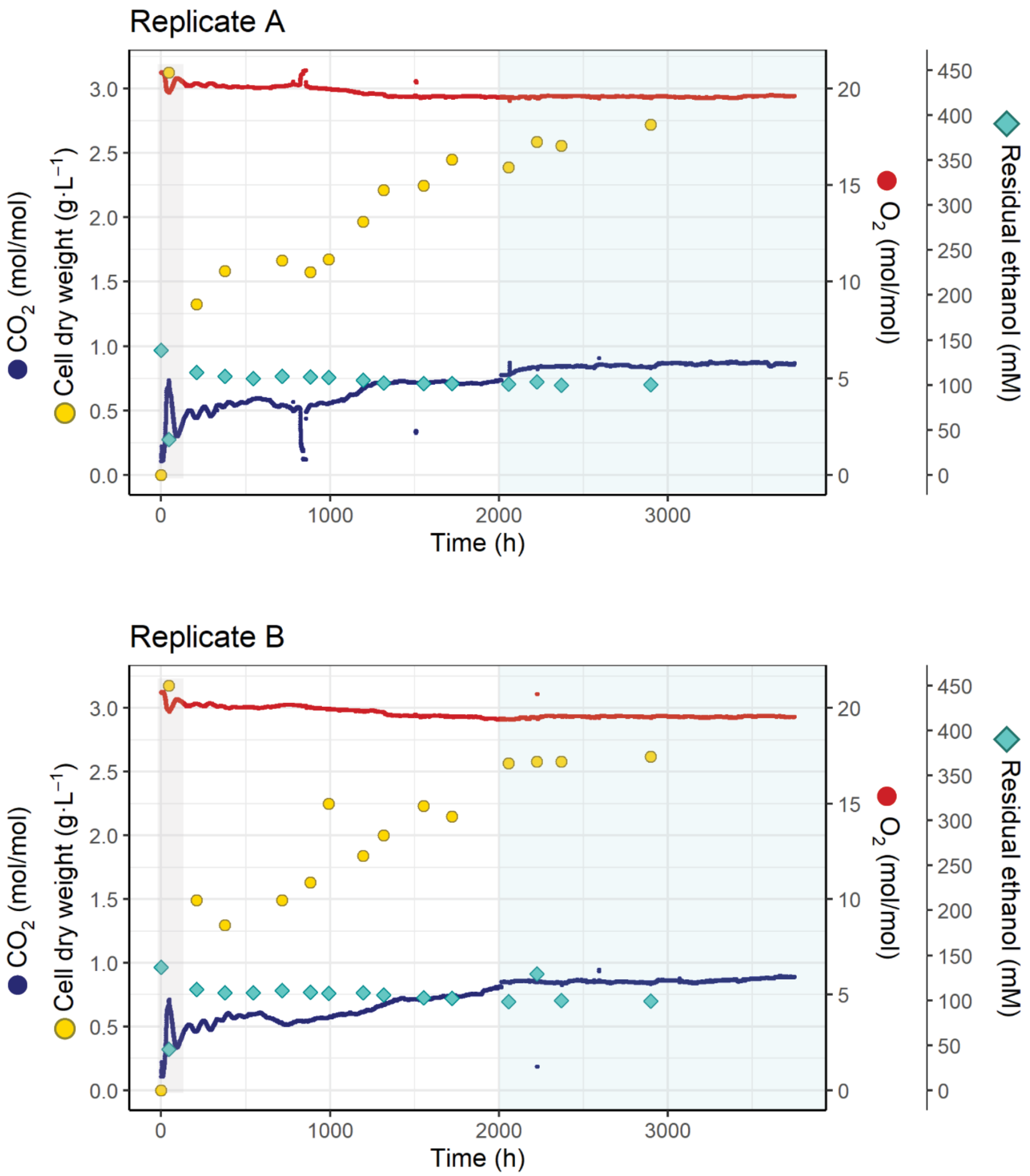


**Figure S11. Prolonged chemostat culture of *S. cerevisiae* to evolve towards mito-ARG8 mRNA import in mitochondria** Profile of two replicate bioreactors, inoculated with a mixture of IMC224-IMC235, arginine auxotrophic strains expressing mito-ARG8 mRNA and the PreCOX4-mRuby2 protein. The chemostats were operated at a dilution rate of 0.1 $h^{-1}$, under nitrogen (arginine) limitation and ethanol as source carbon and energy source. The grey area represents the batch phase, the blue area highlights the phase with relatively steady biomass concentration.

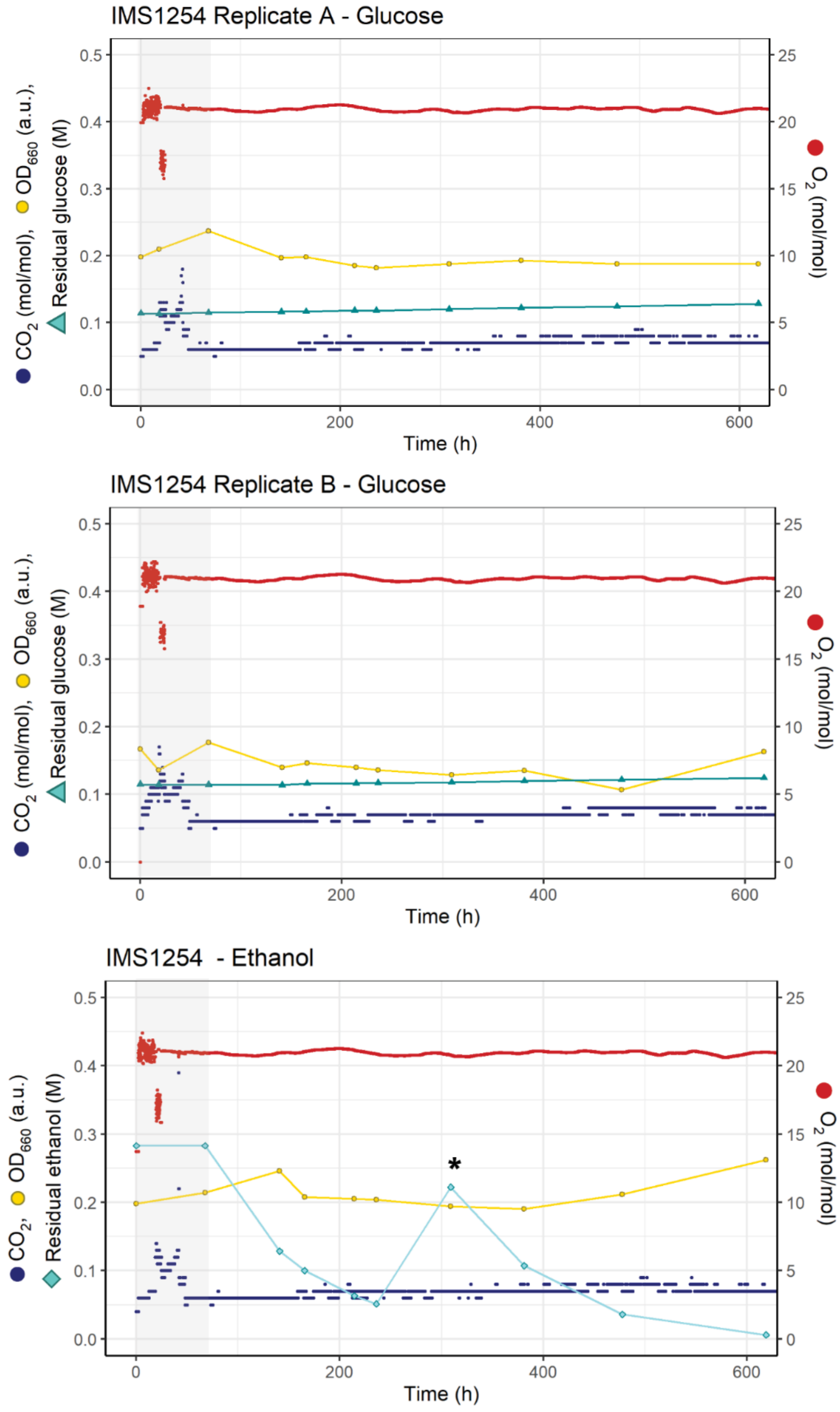


**Figure S12. Physiological characterization of the evolved single colony isolate IMS1254 in batch cultivation in a bioreactor in the absence of arginine.** Growth in synthetic medium supplied with ammonium as sole nitrogen source and either glucose (A and B) or ethanol (C) as sole carbon source. $CO_2$ and $O_2$ concentrations were measured continuously in the off-gas of the bioreactor. Optical density ($OD_{660}$), cell count and the residual ethanol and glucose concentrations were measured offline. The pre-cultures were grown in shake flasks in the presence of arginine. After inoculation of the bioreactor with this pre-culture, a first batch was run to deplete the residual arginine carried over from the pre-culture (grey area). At the end of this first batch, the biomass was let to settle by stopping stirring and aeration and the bioreactor was partially emptied and refilled with fresh, arginine-free medium. The white background reflects growth and metabolic activity of IMS1254 in the complete absence of arginine. The asterisk in panel C indicates additional spiking of ethanol to prevent depletion of the carbon source by evaporation.

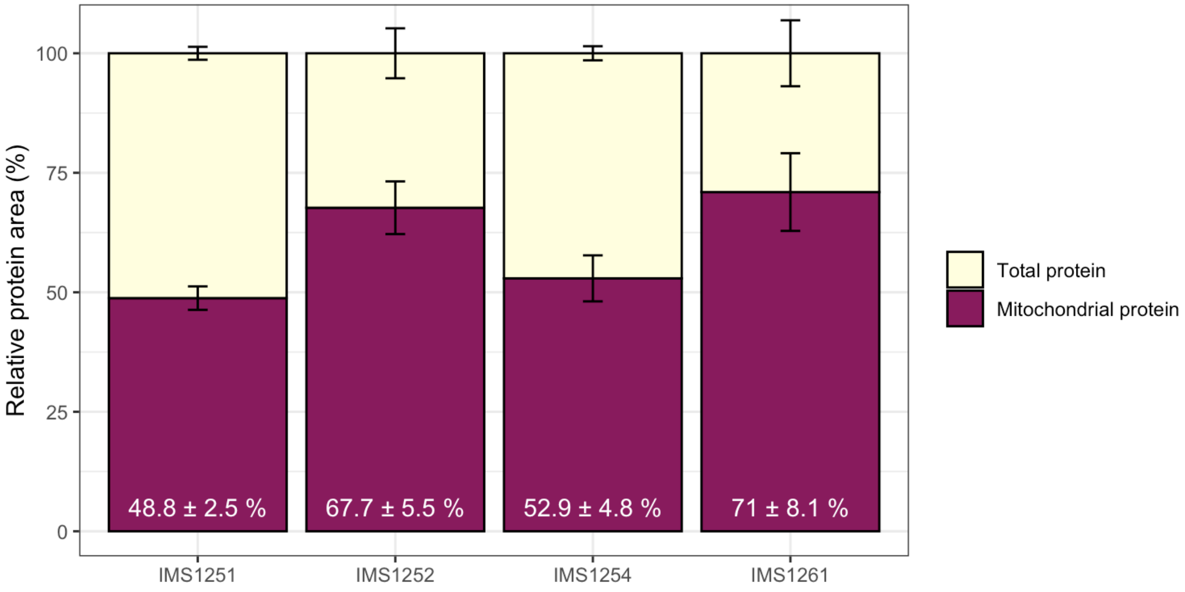


**Figure S13. Fraction of mitochondrial protein detected in each proteome.** 50 to 70% of the detected proteins are mitochondrial, showing a successful enrichment of mitochondria by the isolation protocol. These results were based on the accumulative protein areas of protein that were "mitochondrial" according to their Gene Ontology descriptor versus the total protein areas of each proteome. Error bars were calculated from values obtained for biological duplicates.

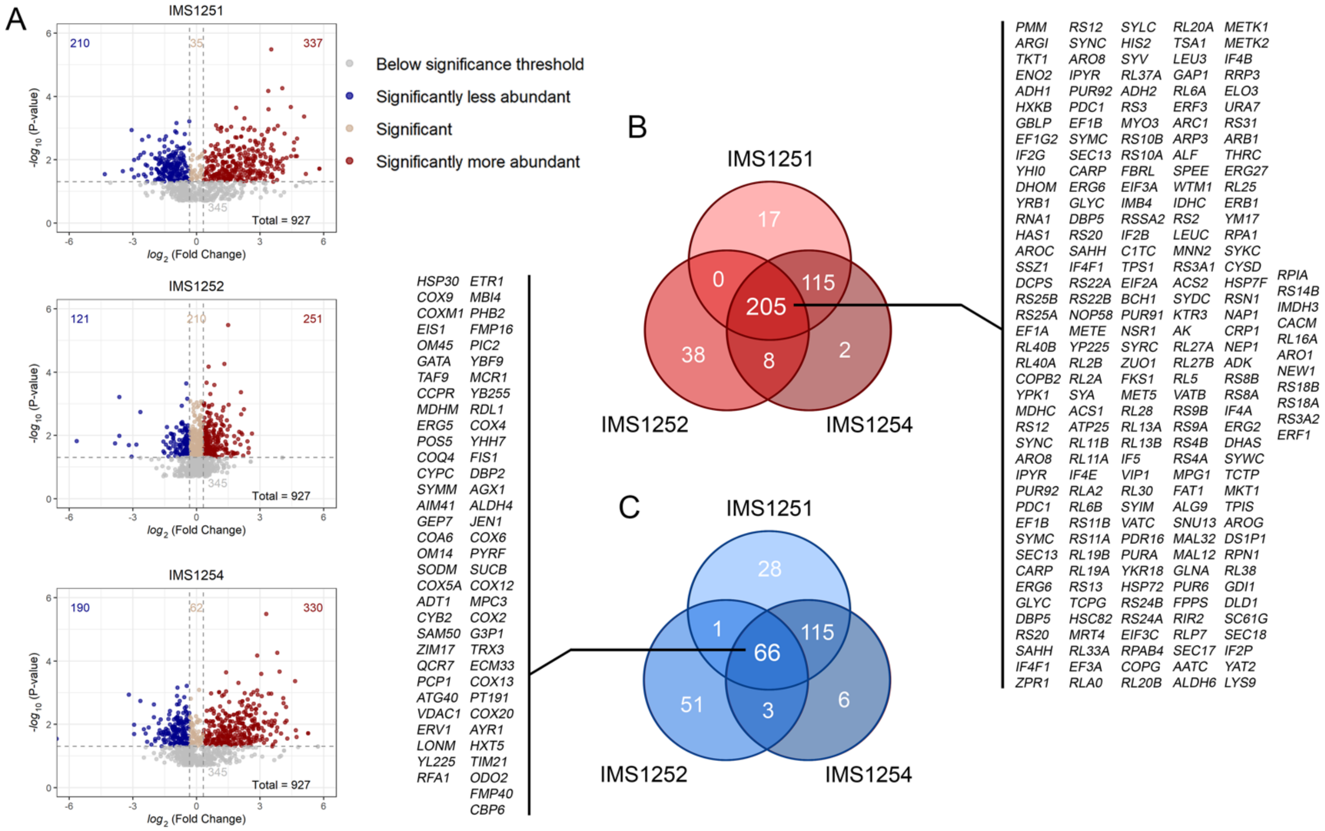


**Figure S14. Global proteome changes between the isolated mitochondria of evolved strains IMS1251, IMS1252 and IMS1254.** A) The fold changes were normalized to the proteome of the unevolved starting population IMS1261. The log2 of the abundance fold change between the two conditions was plotted against the significance (-$\log_{10}$(p)), using a p-value threshold of < 0.05 and a fold change threshold of >1.25 (which corresponds to a log2 fold change threshold +/- 0.32). Red depicts proteins with higher abundance in the evolved strains as compared to the starting population, blue a lower abundance, and beige a similar abundance. B) Venn diagram of the significantly more abundant proteins of each strain (indicated in red in the volcano plots of (A)). Numbers indicate the number of proteins more abundant in one, two or all three strains. The 205 gene names of proteins more abundant in all three strains are listed. C) Venn diagram of the significantly less abundant proteins of each strain (indicated in blue in the volcano plots of (A)). Numbers indicate the number of proteins more abundant in one, two or all three strains. The 66 gene names of proteins less abundant in all three strains are listed.

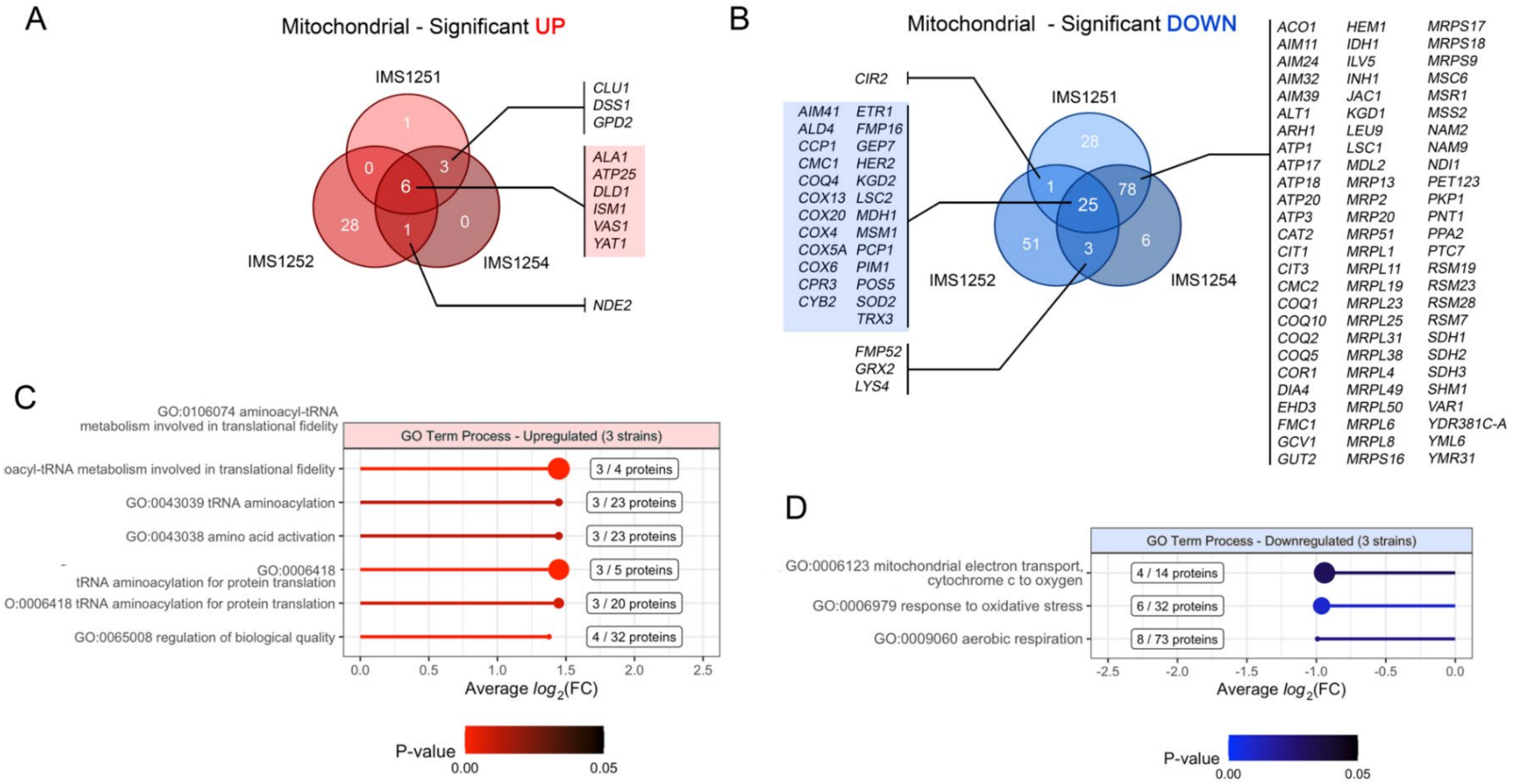


**Figure S15. Proteome response of mitochondrial proteins detected in IMS1251, IMS1252 and IMS1254 compared to unevolved parental population IMS1261.** A) Venn diagram representing the gene names of mitochondrial proteins with significantly higher abundance in the evolved isolates as compared to the unevolved parental population. Numbers indicate the number of proteins more abundant in one, two or all three strains. Proteins more abundant in more than one strain are listed, proteins more abundant in all three strains are indicated in red. B) Venn diagram of the significantly less abundant mitochondrial proteins of each strain. Numbers indicate the number of proteins more abundant in one, two or all three strains. gene names of proteins less abundant in more than one strain are listed, proteins less abundant in all three strains are indicated in blue. C) GO-term analysis of six proteins that were more abundant in all strains (indicated in red in (A)) and D) GO-term analysis of 25 proteins that were less abundant in all strains (indicated in blue in (B)). Proteins were classified according to Gene Ontology (GO) terms of the types "Process". The length of each bar indicates the average measured *log*$_2$(Fold Change) of all proteins associated with the GO-term, The size of the dot indicates the enrichment strength (number of detected proteins in each term/number of proteins of each term in the reference list), which is also indicated next to each bar. The color of each bar indicates the significance of the enrichment, all were below p = 0.05 (Bonferroni corrected).

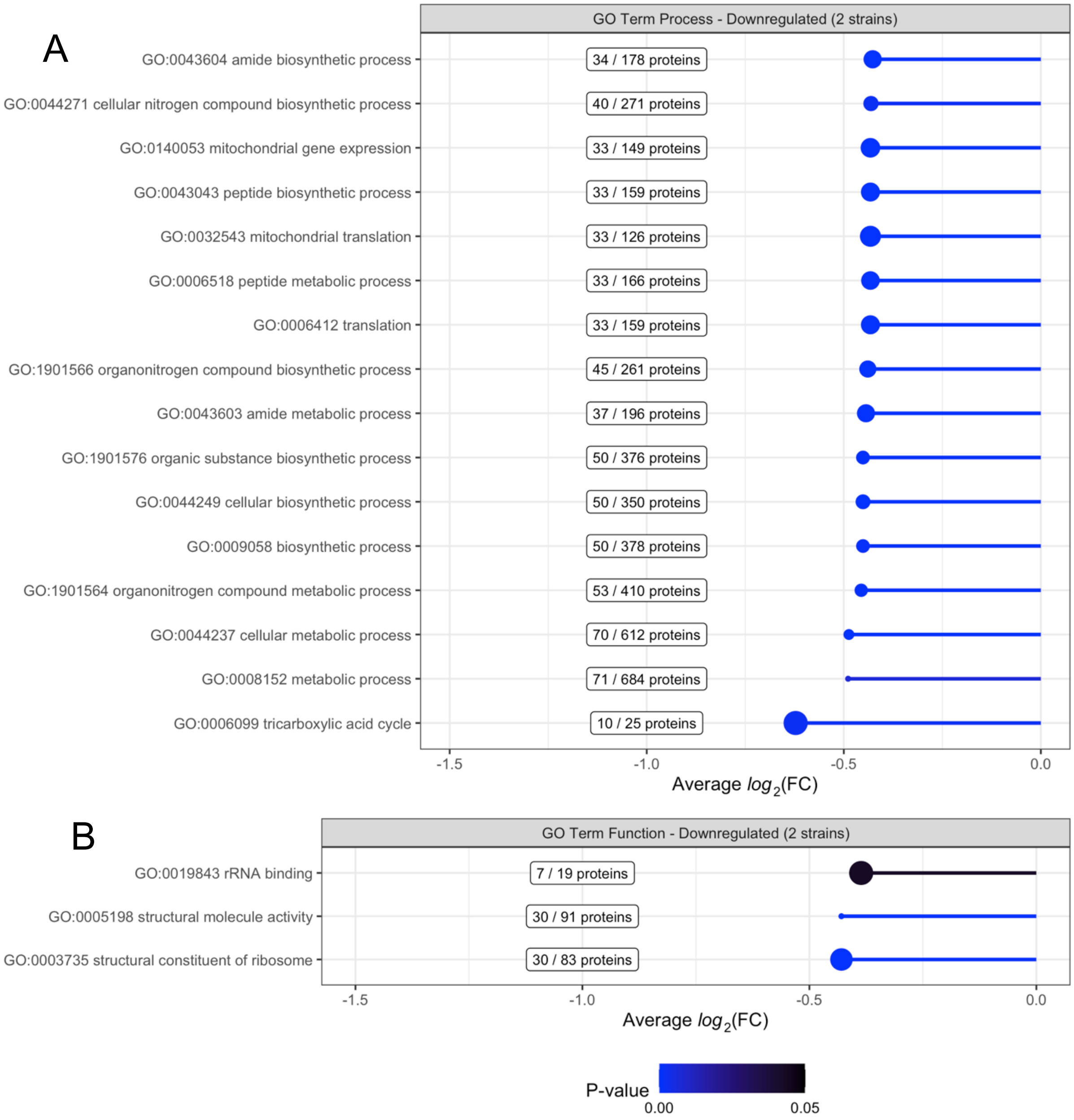


**Figure S16. GO-term analysis of 78 proteins with decreased abundance in both strains IMS1251 and IMS1254 as compared to the unevolved parental population IMS1261**. These data suggest an enrichment for proteins involved in nitrogen metabolism and protein synthesis. Proteins were classified according to Gene Ontology (GO) terms of the type (A) "Process" and (B) "Function". The length of each bar indicates the average measured $log_2$(Fold Change) of all proteins associated with the GO-term, The size of the dot indicates the enrichment strength (number of detected proteins in each term/number of proteins of each term in the reference list), which is also indicated next to each bar. The color of each bar indicates the significance of the enrichment, all were below p = 0.05 (Bonferroni corrected). Gene names associated with each GO-term are listed in SI file 2

### Supplementary files

SI File 1 (*SI_1_WGSdata.xlsx)* – Whole genome sequencing data analysis, containing all annotated mutations found in the evolved strains

SI File 2 (*SI_2_Proteomics_GOterm.xlsx*) – Abundances and annotated proteins found by proteomics analysis of evolved strains, including GO term analysis.

### Supplementary data references


1. Kolesnikova O, Kazakova H, Comte C, Steinberg S, Kamenski P, Martin RP, et al. Selection of RNA aptamers imported into yeast and human mitochondria. RNA. 2010;16(5):926–41. Epub 2010/03/30. doi: 10.1261/rna.1914110. PubMed PMID: 20348443; PubMed Central PMCID: PMCPMC2856887.
2. Zelenka J, Alán L, Jabůrek M, Ježek P. Import of desired nucleic acid sequences using addressing motif of mitochondrial ribosomal 5S-rRNA for fluorescent *in vivo* hybridization of mitochondrial DNA and RNA. Journal of bioenergetics and biomembranes. 2014;46(2):147–56.
3. Wang G, Chen H-W, Oktay Y, Zhang J, Allen EL, Smith GM, et al. PNPase regulates RNA import into mitochondria. 2010;142(3):456–67.
4. Mans R, van Rossum HM, Wijsman M, Backx A, Kuijpers NG, van den Broek M, et al. CRISPR/Cas9: a molecular Swiss army knife for simultaneous introduction of multiple genetic modifications in *Saccharomyces cerevisiae*. FEMS yeast research. 2015;15(2).
5. Goedhart J, Von Stetten D, Noirclerc-Savoye M, Lelimousin M, Joosen L, Hink MA, et al. Structure-guided evolution of cyan fluorescent proteins towards a quantum yield of 93%. Nature communications. 2012;3(1):1–9.
6. Lee ME, DeLoache WC, Cervantes B, Dueber JE. A highly characterized yeast toolkit for modular, multipart assembly. ACS synthetic biology. 2015;4(9):975–86.
7. Boonekamp FJ, Knibbe E, Vieira-Lara MA, Wijsman M, Luttik MAH, van Eunen K, et al. Full humanization of the glycolytic pathway in *Saccharomyces cerevisiae*. Cell Reports. 2022;39(13):111010. doi: https://doi.org/10.1016/j.celrep.2022.111010.
8. Bouwknegt J, Koster CC, Vos AM, Ortiz-Merino RA, Wassink M, Luttik MAH, et al. Class-II dihydroorotate dehydrogenases from three phylogenetically distant fungi support anaerobic pyrimidine biosynthesis. Fungal Biology and Biotechnology. 2021;8(1):10. doi: 10.1186/s40694-021-00117-4.
9. Koster CC, Kleefeldt A, van den Broek M, Luttik M, Daran J-M, Daran-Lapujade P. Long-read direct RNA sequencing of the mitochondrial transcriptome of *Saccharomyces cerevisiae* reveals condition-dependent intron turnover. bioRxiv. 2023:2023.01. 19.524680.
10. Koster C. Exploring the potential of yeast mitochondria for synthetic cell research [PhD dissertation]: Delft University of Technology; 2023.
11. Perli T, Moonen DPI, Broek Mvd, Pronk JT, Daran J-M. Adaptive laboratory evolution and reverse engineering of single-vitamin prototrophies in *Saccharomyces cerevisiae*. Applied and Environmental Microbiology. 2020;86(12):e00388–20. doi: doi:10.1128/AEM.00388-20.
12. Gruber AR, Lorenz R, Bernhart SH, Neuböck R, Hofacker IL. The Vienna RNA Websuite. Nucleic Acids Research. 2008;36(suppl_2):W70–W4. doi: 10.1093/nar/gkn188.
13. Lorenz R, Bernhart SH, Höner zu Siederdissen C, Tafer H, Flamm C, Stadler PF, et al. ViennaRNA Package 2.0. Algorithms for Molecular Biology. 2011;6(1):26. doi: 10.1186/1748-7188-6-26.
14. Esakova O, Krasilnikov AS. Of proteins and RNA: the RNase P/MRP family. Rna. 2010;16(9):1725–47.
15. Duvaud S, Gabella C, Lisacek F, Stockinger H, Ioannidis V, Durinx C. Expasy, the Swiss Bioinformatics Resource Portal, as designed by its users. Nucleic Acids Research. 2021;49(W1):W216–W27. doi: 10.1093/nar/gkab225.
